# Spatio-temporal control of DNA replication by the pneumococcal cell cycle regulator CcrZ

**DOI:** 10.1101/775536

**Authors:** Clement Gallay, Stefano Sanselicio, Mary E. Anderson, Young Min Soh, Xue Liu, Gro A. Stamsås, Simone Pelliciari, Renske van Raaphorst, Julien Dénéréaz, Morten Kjos, Heath Murray, Stephan Gruber, Alan D. Grossman, Jan-Willem Veening

## Abstract

Most bacteria replicate and segregate their DNA concomitantly while growing, before cell division takes place. How bacteria synchronize these different cell cycle events to ensure faithful chromosome inheritance is poorly understood. Here, we identified a conserved and essential protein in pneumococci and related Firmicutes named CcrZ (for <u>C</u>ell <u>C</u>ycle <u>R</u>egulator protein interacting with Fts<u>Z</u>) that couples cell division with DNA replication by controlling the activity of the master initiator of DNA replication, DnaA. The absence of CcrZ causes mis-timed and reduced initiation of DNA replication, which subsequently results in aberrant cell division. We show that CcrZ from *Streptococcus pneumoniae* directly interacts with the cytoskeleton protein FtsZ to place it in the middle of the newborn cell where the DnaA-bound origin is positioned. Together, this work uncovers a new mechanism for the control of the bacterial cell cycle in which CcrZ controls DnaA activity to ensure that the chromosome is replicated at the right time during the cell cycle.

## Main

Most organisms have mechanisms ensuring that their genome is replicated and segregated prior to cell division. In many bacterial species, DNA replication and cell division occur concomitantly^1,2,3^. Different models emerged from the mid-1900’s to explain how bacterial cells handle DNA replication together with cell division in *Escherichia coli* or *Bacillus subtilis*^4,5,67^. The current cell-size control model suggests that cells initiate DNA replication independently from their original size, and grow to a constant size independently from their size at birth (adder model)^8,9,10,11,12^. How cells sense changes in cell size and convert it to trigger replication initiation is not known, but these models imply the existence of regulatory controls^3,13,14,15^. However, no such cell cycle regulator has been reported yet for bacteria. Specific regulatory models have been proposed for *E. coli*^16,17,18^, but these are not applicable to most other organisms, and especially Gram-positive bacteria, that do not contain the proteins proposed to be the regulators. Furthermore, most of the mechanisms known to regulate the initiation of replication and the activity of the replication initiator DnaA in *E. coli* do not exist in other bacteria^19,20,21,22,23^. This pinpoints a divergence between regulatory systems within bacteria. In line with this notion, changes in DNA replication initiation were shown to alter cell size in *E. coli* and *B. subtilis* but the converse was not true for *B. subtilis*^24,25^. Taken together, current data indicates that bacteria evolved different mechanisms to coordinate their cell cycle events.

Although *E. coli* and *B. subtilis* use different systems for regulating their cell cycle, the way they localize their division site is conserved, as both organisms use a variant of the Min system to prevent polymerization of the tubulin-like protein FtsZ away from mid-cell^26,27^. Both species also have a nucleoid occlusion system (Noc) inhibiting Z-ring formation over the chromosome to prevent “cutting” of the chromosome during cell division^28^. Together, the Min and Noc systems ensure that cell division and septation occur when both sister chromatids have been fully replicated and segregated. These systems are however not conserved within all bacteria as the Gram-positive opportunistic human pathogen *S. pneumoniae* lacks homologs of the Min and Noc systems^29^. In contrast to *E. coli* and *B. subtilis*, the pneumococcal Z-ring forms readily over the nucleoid^29,30^. Recently, a pneumococcal specific protein called RocS was identified that might fulfil a similar function as the Noc system by connecting chromosome segregation with capsule production^31^. Another *S. pneumoniae* specific protein, called MapZ was shown to guide Z-ring formation, analogous to the Min system in other bacteria^32,33^. During cell growth, nascent MapZ rings are pushed apart by septal peptidoglycan synthesis, allowing for FtsZ polymers to continuously assemble at the newly formed septum^34^. Importantly, the position of the origin of replication (*oriC*) was shown to be crucial for division site selection in *S. pneumoniae* and the origins mark the approximate positions of future division sites^35^. In *S. pneumoniae*, cell division and DNA replication are thus intimately connected. Critically however, it remains unknown how the cell senses when a new round of replication should be initiated.

We hypothesized that an unknown factor could be responsible for coordination of cell division and DNA replication in the pneumococcus. Using high throughput gene silencing with CRISPRi of all essential genes of *S. pneumoniae*^36^, we examined proteins leading to DNA content defects upon depletion. Here, we describe the identification of CcrZ, a conserved protein that activates DnaA to trigger initiation of DNA replication. Pneumococcal CcrZ localizes at the division site in a FtsZ-dependent manner and its inactivation leads to division defects. Together, our findings show that CcrZ acts as a novel spatio-temporal link between cell division and DNA replication in *S. pneumoniae*.

### CcrZ is a conserved bacterial cell cycle protein

We previously generated a knock-down library using CRISPRi (clustered regularly interspaced short palindromic repeats interference) targeting 348 conditionally essential genes of the serotype 2 strain *S. pneumoniae* D39V that were identified by Tn-Seq (transposon-insertion sequencing)^36^. Here, we investigated the function of *spv_0476*, encoding a protein of unknown function that is conserved in most Firmicutes (>30% identity) (Extended Data Fig. 1a). Silencing of *spv_0476* by CRISPRi led to a drastic reduction of the growth rate as well as appearance of anucleate cells as visualized by DAPI staining (Fig. 1a,b). We renamed SPV_0476 to CcrZ (for <u>C</u>ell <u>C</u>ycle <u>R</u>egulator protein interacting with Fts<u>Z</u>) for reasons explained below. *ccrZ* is in an operon with *trmB*, which encodes a tRNA methyltransferase and this operon structure is conserved across Firmicutes (Extended Data Fig. 1a). To exclude the possibility that the observed phenotypes of *ccrZ* silencing were caused by polar effects on *trmB* expression, we constructed a deletion of *trmB*. This deletion did not lead to any growth defect (Extended Data Fig. 1b left panel). While Tn-seq indicated that *ccrZ* is essential^36^, we were able to generate a Δ*ccrZ* deletion mutant, although cells grew very slowly. We therefore constructed a depletion of CcrZ by ectopically expressing CcrZ under control of either an IPTG- or a ZnCl_2_-inducible promoter (*P_lac_* and *P_Zn_* respectively) and deleted *ccrZ* from its native location (*ccrZ*^−/+^ and *P_Zn_*-*ccrZ*^−/+^ respectively). Depletion of CcrZ led to a significant growth delay at 37°C and 30°C, confirming the CRISPRi observations (Extended Data Fig. 1b). Immunoblotting using a specific antibody raised against purified CcrZ confirmed CcrZ depletion (Extended Data Fig. 1c).

**Fig. 1:**
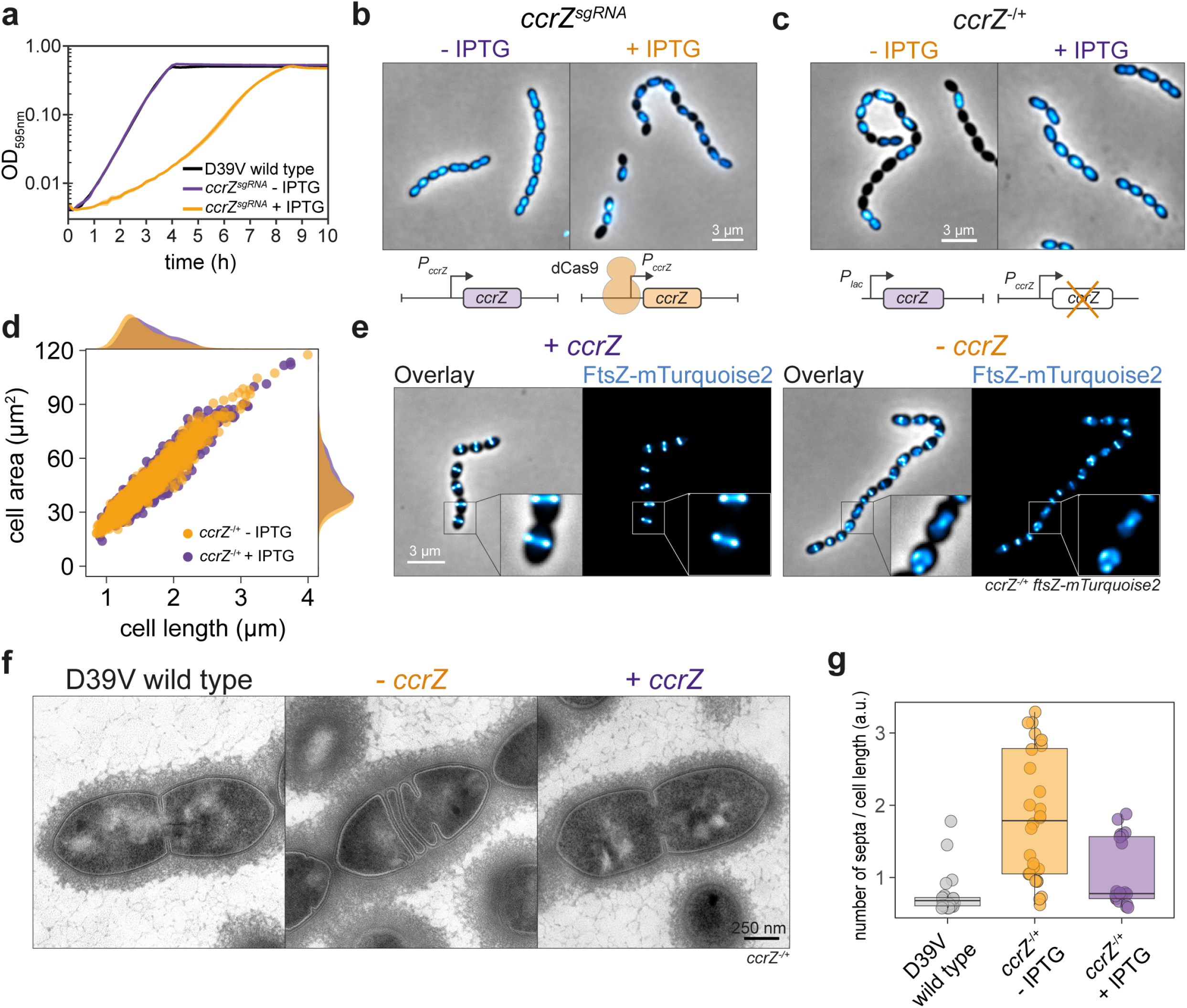
Depletion of CcrZ leads to anucleate cells and division defects. **a**, Growth curve of cells with *ccrZ* targeted by CRISPRi (*ccrZ^sgRNA^* + IPTG) indicates a clear growth defect when *ccrZ* is silenced. **b**, *ccrZ* silencing leads to appearance of anucleate cells, as visualized by DAPI staining. **c**, *ccrZ* depletion by ectopic expression via the IPTG-inducible Plac promoter also leads to cells lacking a nucleoid, as observed by DAPI staining. **d**, Distribution of cell area of *ccrZ*-depleted cells, *ccrZ* depletion leads to a slight decrease in cell length and cell area (*P* value = 2.2 x 10^−16^, Wilcoxon rank sum test). **e**, When a deletion of *ccrZ* is complemented (left panel), FtsZ-mTurquoise2 shows a clear mid-cell localization, while it appears as a blurry signal in several cells upon *ccrZ* depletion (right panel). **f**, Transmission electron microscopy (TEM) indicates that cells depleted for *ccrZ* form multiple, often incomplete, septa. **g**, Distribution of number of septa per cell length as determined by TEM for 22 wild type cells, 28 CcrZ-depleted cells and 17 CcrZ-complemented cells. Each dot represents a measurement. (*P* value = 1 x 10^−6^ for wild type vs *ccrZ*-depleted cells and *P* value = 0.0013 for *ccrZ*-complemented vs *ccrZ*-depleted cells, Wilcoxon rank sum test with Bonferroni adjustment).

In line with the CRISPRi observations, DNA staining of cells depleted for CcrZ showed that 20% of cells lacked a nucleoid (Fig. 1c, 442 cells counted). To test whether the *ccrZ*-deletion phenotype was conserved in other Gram-positive bacteria, we silenced *ccrZ* (*SAOUHSC_01866*, here *ccrZ_Sa_*) in *Staphylococcus aureus* SH1000 using CRISPRi and deleted the *Bacillus subtilis* 168 *ccrZ* homolog (*ytmP*, here *ccrZ_Bs_*). Upon *ccrZ_Sa_* silencing in *S. aureus*, we observed a high proportion of anucleate cells, as well as a delay in growth. In contrast, no anucleate cells were observed for *B. subtilis* (Extended Data Fig. 1d). However, cells deleted for *ccrZ_Bs_* were slightly thinner and longer although they grew with a growth rate similar to the wild type (Extended Data Fig. 1d). Interestingly, the *S. pneumoniae ccrZ* deletion could not be complemented by expression of *ccrZ* from either *B. subtilis* or *S. aureus* as only very small colonies were present on agar plates. In contrast, depletion of *S. aureus* CcrZ was rescued by expression of CcrZ from *B. subtilis* (Extended Data Fig. 1d).

In addition to an increase of the number of anucleate cells, CcrZ depletion in *S. pneumoniae* also led to slight morphological defects and modest changes in cell size when analyzed by phase contrast microscopy (slight decrease in length and increase in width) (Fig. 1d). Polysaccharide capsule production has previously been linked to the pneumococcal cell cycle^37^, but capsule production was not impacted as the amount of capsule was similar between a CcrZ mutant and wild type (Extended Data Fig. 1e). To visualize division sites in live cells, we constructed a translational fusion of mTurquoise2 to FtsZ (as the only copy of FtsZ, expressed from its native genetic location), which assembles into distinct rings at new division sites where it recruits the machinery required to form septa^38^. As shown in Figure 1e, Z-rings were clearly mis-localized upon CcrZ depletion for 3h, with the presence of several aberrant Z-rings in 43% of the cells (Fig. 1e). To obtain more insights into the morphological defects caused by CcrZ depletion and verify that the increased number of septa are not due to the fluorescent protein fused to FtsZ, we employed transmission electron microscopy (TEM) in untagged cells. While not evident by phase contrast microscopy, when *ccrZ* was depleted we observed frequent aberrant septum formation using TEM, in line with the FtsZ localization data, and many cells harbored two (18 %) to four (4 %) septa while only one septum is observed in 91% of wild type cells (Fig. 1f,g).

### *S. pneumoniae* CcrZ is part of the divisome

As CcrZ seems to be involved in both chromosome biology and cell division, we examined its subcellular localization. Strikingly, immunofluorescence on fixed cells using a CcrZ-specific antibody demonstrated a clear mid-cell localization (Extended Data Fig. 2a). To assess the localization of CcrZ in live cells, we created several functional fusions of a green fluorescent protein to the N-terminus of CcrZ (*gfp-ccrZ*) or a red fluorescent protein to the C-terminus (*ccrZ-mKate2*) and inserted either construct at the native *ccrZ* locus (Extended Data Fig. 1c). Visualization of fluorescently tagged CcrZ by epifluorescence microscopy in live bacteria showed that CcrZ localizes at mid-cell (Fig. 2a). This localization was also conserved in both the TIGR4 and unencapsulated R6 strains (Extended Data Fig. 2b). Interestingly, CcrZ_Sa_ and CcrZ_Bs_ did not localize as clear rings at mid-cell in *S. aureus* and *B. subtilis* (Extended Data Fig. 2b), indicating that the activity and/or localization of CcrZ in these organisms is regulated differently. In order to get higher spatial resolution of *S. pneumoniae* CcrZ, 240 images (16 stacks) on live cells were acquired using 3D-structured illumination microscopy (3D-SIM) and reconstructed to generate a super resolution image and a 3D fluorescence image of GFP-CcrZ. As shown in Fig. 2b and Supplementary Video 1, CcrZ forms a patchy ring at mid-cell. Furthermore, time-lapse microscopy showed that CcrZ disassembles from the old septum to assemble at the newly formed division site (Supplementary Video 2).

**Fig. 2:**
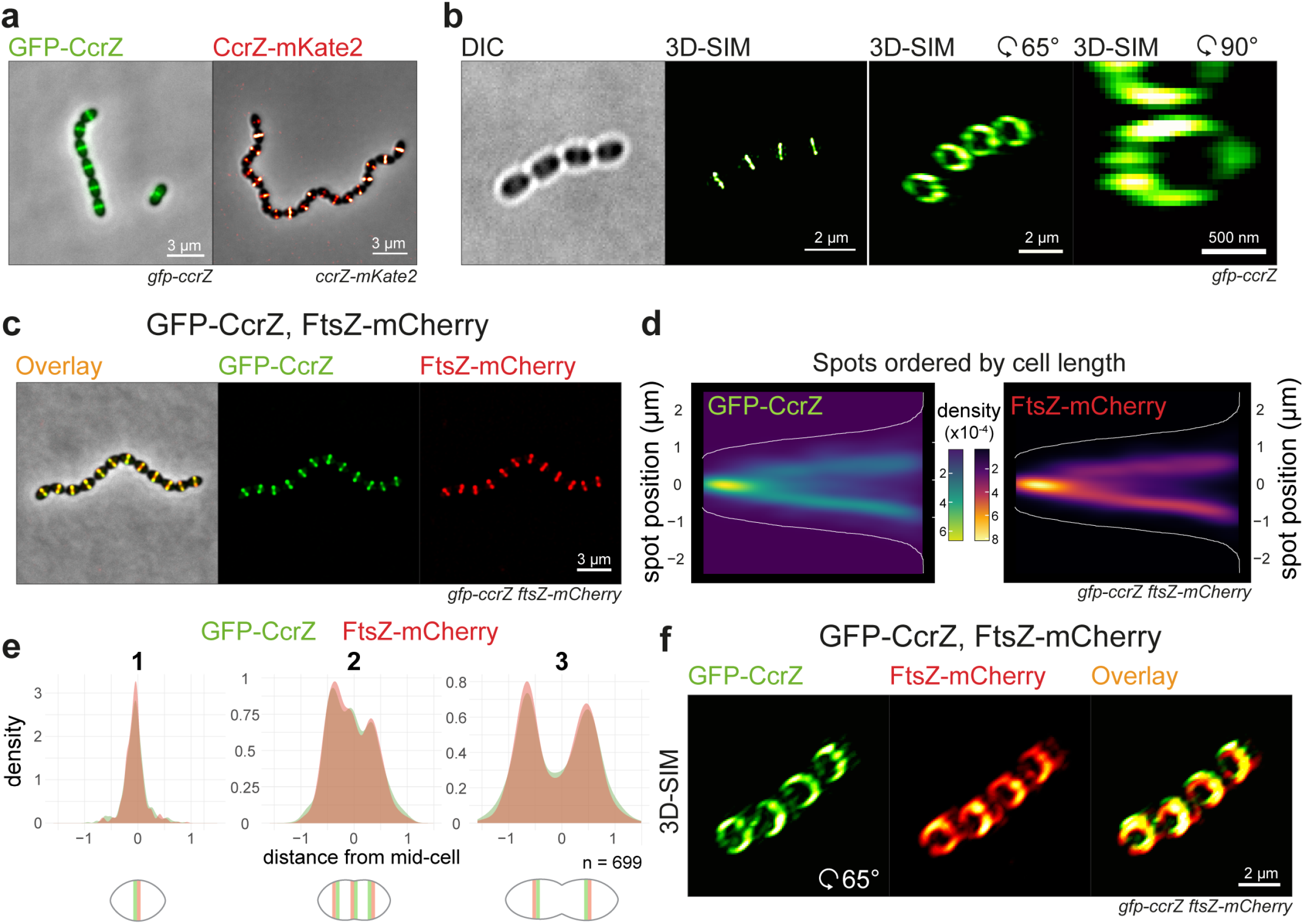
CcrZ co-localizes with FtsZ at new division sites. **a**, CcrZ localizes at mid-cell in live wild type *S. pneumoniae* cells as observed by epifluorescence microscopy of GFP-CcrZ and CcrZ-mKate2. **b**, 3D-SIM of GFP-CcrZ and reconstructed volume projection (right) indicate that CcrZ forms a patchy ring at mid-cell. **c**, GFP-CcrZ and FtsZ-mCherry co-localize in wild type cells. **d**, Localization signal of GFP-CcrZ and FtsZ-mCherry in 699 cells of a double labelled *gfp-ccrZ ftsZ-mCherry* strain, ordered by cell length and represented by a heatmap. **e**, GFP-CcrZ and FtsZ-mCherry co-localize during the entire cell cycle, as visualized when signal localization over cell length is grouped in three quantiles. **f**, 3D-SIM co-localization of GFP-CcrZ and FtsZ-mCherry shows a clear co-localizing ring with identical patchy pattern. Note that for clarity, we did not correct for chromatic shift in the overlay.

To test whether the mid-cell localization of *S. pneumoniae* CcrZ coincides with FtsZ, we constructed a CcrZ / FtsZ double-labelled strain (*gfp-ccrZ ftsZ-mCherry*). As shown in Fig. 2c, CcrZ co-localized with FtsZ and analysis of still images from exponentially growing cells corroborated this observation (Fig. 2c-e and Supplementary Video 3). Note that the FtsZ-mCherry fusion did affect the growth or morphology of the cells^39^. 3D-SIM also indicated an overlap of GFP-CcrZ and FtsZ-mCherry as well as a similar circular co-localizing pattern at mid-cell (Fig. 2f, Extended Data Fig. 2c and Supplementary Video 4).

Prediction of CcrZ’s topology using TMHMM^40^ did not indicate the presence of a transmembrane domain; CcrZ’s septal localization might then rely on another partner. To identify possible partners, we purified GFP-CcrZ expressed from *S. pneumoniae* and untagged cytosolic sfGFP as a control using anti-GFP nanobodies (GFP-Trap) without cross-linking, and directly analyzed the purified fraction by liquid chromatography□tandem mass spectrometry (LC□MS/MS). Interestingly, we found an enrichment (> 5-fold change) for several proteins from the divisome (*e.g.,* FtsZ, PBP2X and EzrA) (Supplementary Table 3). To determine which of the candidates might interact directly with CcrZ, we used the NanoBit complementation reporter assay^41,42^, which uses an enhanced luciferase separated into two different fragments (large bit (LgBit) and small bit (SmBit), respectively). Fusion of two different interacting proteins to each fragment can restore the activity of the luciferase and, in presence of a furimazine-based substrate, produce light^41^. Accordingly, we fused the C-terminal extremity of CcrZ to LgBit (*ccrZ-LgBit*) and putative partners to SmBit and integrated the different constructs at their respective loci under native control. We also fused SmBit to other proteins known to localize at the septum (Cps2E, FtsA, FtsW and ZapA), or to the highly abundant histone-like protein HlpA localizing at the nucleoid and used a strain expressing both HlpA-LgBit and HlpA-SmBit as a positive control of interaction. After addition of the substrate, we could detect a strong and reproducible signal when FtsZ was fused to SmBit and CcrZ to LgBit, as well as a weaker signal for FtsA, EzrA and ZapA, and no detectable signal for any of the other proteins (Fig. 3a). This result indicates that FtsZ and CcrZ in *S. pneumoniae* are in very close proximity in space. Interestingly, using a strain expressing both CcrZ-LgBit and CcrZ-SmBit, a weak signal was observed indicating that CcrZ might also self-interact (Fig. 3a).

**Fig. 3:**
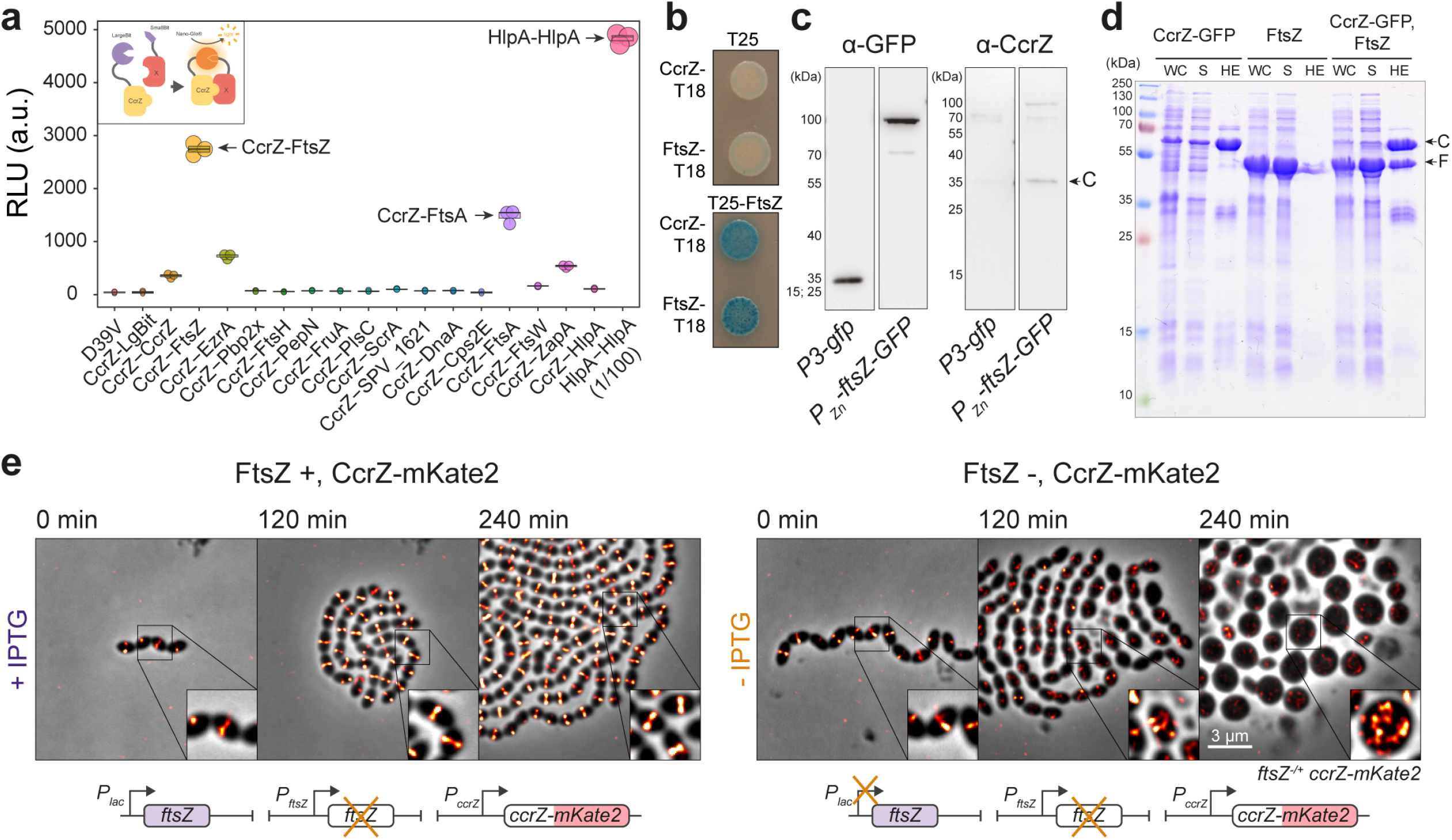
CcrZ directly interacts with FtsZ. **a**, Ssplit-luciferase assay using several combinations with CcrZ-LgBit reveals that CcrZ and FtsZ are in very close proximity, as indicated by a high luminescence signal. FtsA, EzrA and ZapA, all three interacting directly with FtsZ, also gave a slight signal. *hlpA-LgBit hlpA-SmBit* (HlpA-HlpA), here diluted 100 times, is used as positive control. Each dot represents the average of 15 measurements of a technical replicate, with the size of the dot representing the SEM. **b**, FtsZ-CcrZ interaction confirmation by bacterial two-hybrid assay. T25 is the empty vector pST25 and T25-FtsZ corresponds to vector pST25-FtsZ used in combination with pUT18-CcrZ (CcrZ-T18) and pUT18-FtsZ (FtsZ-T18). **c**, Affinity purification of FtsZ-GFP from *S. pneumoniae* cells (2^nd^ lane) also pulls down untagged CcrZ (4^th^ lane). Purification of GFP alone (first lane) did not pull CcrZ down (3^rd^ lane). **d**, FtsZ from *S. pneumoniae* expressed in *E. coli* co-purifies with CcrZ_Sp_-GFP by affinity purification. WC: whole cell extract, S: supernatant, HE: heat eluted products, C: CcrZ-GFP, F: FtsZ. **e**, Epifluorescence time-lapse microscopy of CcrZ-mKate2 at 37°C in presence (left panel) or absence (right panel) of FtsZ. When FtsZ amounts are reduced, cells increase their size and CcrZ is de-localized from mid-cell.

To confirm the observed interaction with FtsZ, we used a bacterial two-hybrid assay in *E. coli*^43^. Again, we observed a robust interaction between CcrZ and FtsZ, while T25-FtsZ did not interact with the empty vector alone, strongly suggesting that CcrZ directly binds to FtsZ (Fig. 3b). Co-immunoprecipitation of FtsZ-GFP from *S. pneumoniae* cells confirmed the *in vivo* interaction with CcrZ (Fig. 3c). Affinity purification of CcrZ_Sp_-GFP when over-expressing FtsZ_Sp_ in *E. coli* also confirmed this interaction as we were able to co-purify FtsZ in large amounts (Fig. 3d). To test whether the localization of CcrZ depends on FtsZ, we constructed a strain expressing CcrZ-mKate2 as well as a second copy of FtsZ under the control of an IPTG-inducible promoter and deleted the native *ftsZ* gene (*ftsZ*^−/+^). As expected, FtsZ depletion led to aberrant cell morphology and, consistent with a FtsZ-CcrZ interaction, CcrZ-mKate2 was rapidly mis-localized and the signal was spread throughout the cytoplasm (Fig. 3e and Supplementary Video 5). In total, we conclude that CcrZ localizes to new division sites via a direct interaction with FtsZ.

### CcrZ controls DNA replication

As shown in Fig. 1c, when cells are depleted for CcrZ, a large proportion of cells become anucleate. To investigate the consequences of lack of CcrZ on chromosome segregation in live cells, we introduced a translational fluorescent fusion of HlpA^44^ and deleted *ccrZ*. Localization of HlpA-mKate2 in this slow growing Δ*ccrZ* mutant showed similar results to DAPI stained cells depleted for CcrZ and we observed that 19 % of cells lacked a nucleoid signal (Extended Data Fig. 3a, 4855 cells counted). Time-lapse imaging indicated that cells with defective DNA content had either no DNA at all or chromosomes “guillotined” during septum closure suggesting reduced nucleoid occlusion control in Δ*ccrZ* (Fig. 4a and Supplementary Video 6). We also co-localized FtsZ-CFP with HlpA-mKate2 while depleting CcrZ for a short period of time (2h). Interestingly, we observed many cells with a chromosome localized at only one half of the cell, at one side of the Z-ring (Fig. 4b). The absence of DNA in the other half of the cell could be explained by defective DNA segregation, by impaired replication or by DNA degradation.

**Fig. 4:**
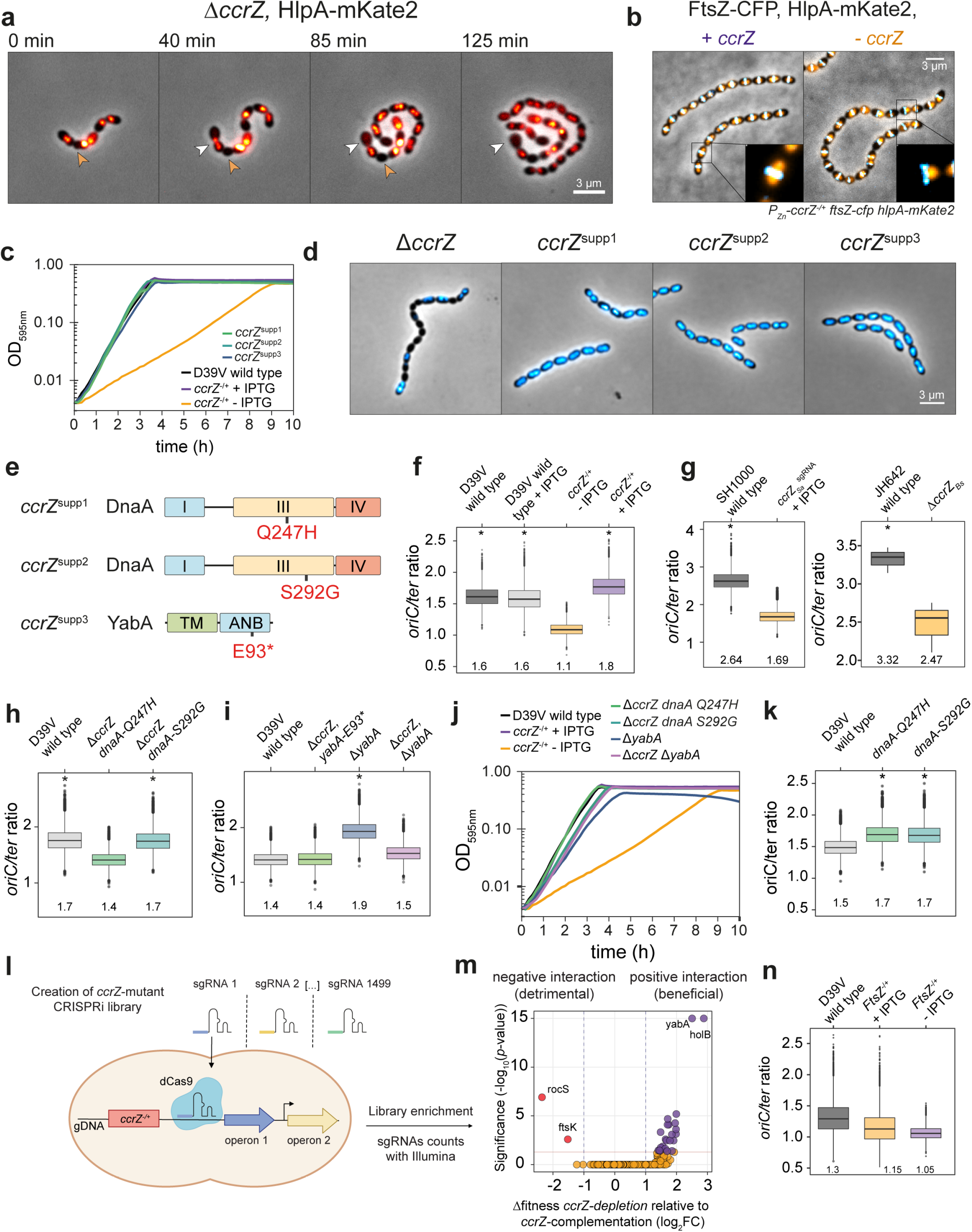
CcrZ-depleted cells under-replicate. **a**, Time-lapse microscopy of HlpA-mKate2 at 30°C in a Δ*ccrZ* mutant shows several cells with defective DNA content. Orange arrows indicate a cell with no nucleoid after cell division; white arrows indicate a cell with “guillotined” DNA. **b**, Co-localization of FtsZ-CFP and HlpA-mKate2 when depleting *ccrZ* indicates that several cells have a nucleoid located only on one side of the Z-ring. **c**, Three isolated *ccrZ* mutants (*ccrZ*^supp1-3^) restore wild type growth to Δ*ccrZ*. **d**, DAPI staining of the three selected *ccrZ* suppressors mutants shows a restoration of DNA content. **e**, Schematic representation of the localization of suppressor mutations in the domain III of DnaA and in the DnaA/DnaN binding motif (ANB) of YabA. TM: tetramerization domain. **f**, *oriC/ter* ratios as determined by RT qPCR for D39V wild type and *ccrZ* depleted cells. Average values are indicated under the boxes. *ccrZ* depletion leads to a clear reduction in *oriC/ter* ratio. See Methods for statistical tests. **g**, *oriC/ter* ratios for *S. aureus* upon *ccrZ_Sa_* depletion (left) and for *B. subtilis* with *ccrZ_Bs_* deletion (right). **h**, *oriC/ter* ratios of strains with *dnaA* mutations re-inserted into a Δ*ccrZ* background show that these mutations restore replication initiation rates. **i**, *yabA* deletion leads to an increase in *oriC/ter* ratios, while suppressor mutation *ccrZ^supp3^* (Δ*ccrZ*, *yabA-E93\**) as well as co-deletion of *yabA* together with *ccrZ* (Δ*yabA* Δ*ccrZ*) restore a wild type ratio. **j**, While *yabA* deletion alters the growth rate, a Δ*yabA* Δ*ccrZ* double mutant grows like wild type. *dnaA Q247H* and *dnaA S292G* mutation also restore a wild type rate in a Δ*ccrZ* mutant. **k**, *dnaA* mutation in a wildtype background increases the *oriC/ter* ratios. **l**, Schematic overview of CRISPRi-seq. *ccrZ*-depletion strain (*ccrZ^−/+^*) was transformed with 1,499 different sgRNAs targeting 2,111 genetic elements of *S. pneumoniae*. These sgRNAs are expressed constitutively. The resulting library is then grown in presence or absence of inducer for production of dCas9 and the genomic DNA isolated. After sequencing of the sgRNA region, reads counts are compared between *ccrZ* depleted induced and complemented induced, indicating which sgRNAs were enriched or deprived. This fold change directly informs whether the gene targeted by the sgRNA becomes more (beneficial) or less (detrimental) essential in a specific genetic background. **m**, CRISPRi-seq of *ccrZ*-depletion vs *ccrZ-*expression shows a positive interaction between *ccrZ* and *yabA / holB* (these two genes are in the same operon) and a negative interaction between *ccrZ* and *ftsK / rocS*. **n**, *oriC/ter* ratios for FtsZ depletion strain (FtsZ^−/+^) shows a reduced ratio when FtsZ is depleted (-IPTG) compared to when it is complemented (+ IPTG).

When attempting to make clean *ccrZ* deletions, in addition to small colonies typical of slow growing mutants, there were also spontaneous large, wild type-sized colonies. Growth analysis of cells from three of these large colonies (*ccrZ^supp1-3^*) showed that cells behaved like wild type and DAPI staining revealed a restoration of wild type DNA content (Fig. 4c,d). To verify whether these wild type-like phenotypes were caused by suppressor mutations, the genomes of these fast-growing strains were sequenced. All three strains still contained the *ccrZ* deletion and, in addition, contained a single nucleotide polymorphism elsewhere in the genome (Fig. 4e). Two missense mutations were found in *dnaA* (DnaA-Q247H and DnaA-S292G) and one nonsense mutation in *yabA* (YabA-E93*). Since DnaA promotes initiation of DNA replication and YabA hampers it by preventing interaction of DnaA with DnaN^45^, we wondered whether the frequency of DNA replication initiation was changed in a *ccrZ* mutant.

To test this hypothesis, we quantified the copy number ratio between chromosomal origin and terminus regions (*oriC/ter* ratios) using real-time quantitative PCR. In a wild type situation, during exponential growth, the *oriC/ter* ratio varies between 1.3 – 1.8, as most cells have started a round of DNA replication (note that in contrast to *E. coli* and *B. subtilis*, multifork replication does not occur in *S. pneumoniae*)^46^. Remarkably, depletion of CcrZ resulted in a significantly decreased DNA replication initiation rate with an *oriC/ter* ratio of 1.1 vs 1.8 for complemented cells (*P* value < 0.05) (Fig. 4f). Interestingly, the same observation was made for both *B. subtilis* and *S. aureus*, where deletion or depletion of CcrZ caused a clear reduction in *oriC/ter* ratios (Fig. 4g). As the identified *ccrZ*-bypass mutations were found in DNA replication initiation regulators, we tested whether they would restore the *oriC/ter* ratio in a fresh *ccrZ* deletion background in *S. pneumoniae*. Indeed, the *oriC/ter* ratios for Δ*ccrZ dnaA-S292G,* Δ*ccrZ dnaA-Q247H* and for *yabA-E93\** (*ccrZ^supp3^*) were like wild type (Fig. 4h,i).

The point mutation found in *yabA* causes premature translation termination at the C-terminus of YabA. When *yabA* alone was replaced by an antibiotic resistance cassette, we observed an increase of replication initiation as well as a reduced growth rate; but when *ccrZ* was co-deleted, wild type like growth and a wild type *oriC/ter* ratio was restored (Fig. 4i,j). DnaA suppressor mutations were located in the AAA+ ATPase domain of DnaA^47^ (Extended Data Fig. 3b) and it was previously reported that specific mutations in this domain could increase the initiation rate in *B. subtilis*^48^. To determine if those mutations alone were able to induce over-initiation, we inserted each *dnaA* mutation into a wild type background strain. Marker frequency analysis detected an increase in the *oriC/ter* ratio for both *dnaA* alleles (Fig. 4k). We conclude that mutations that increase the rate of initiation of DNA replication can rescue the Δ*ccrZ* phenotype.

To gain additional insights into CcrZ function, we performed a genome-wide genetic interaction screen in cells depleted for CcrZ using CRISPRi-seq^49^ (Fig. 4l). This technique relies on the expression of dCas9, controlled by an anhydrotetracycline (aTc) -inducible promoter, and constitutive expression of a specific sgRNAs that together form a roadblock for RNAP and thereby downregulated transcription of the targeted operon. We created a CRISPRi library by transforming *S. pneumoniae P_tet_-dCas9, P_lac_-ccrZ, ΔccrZ* with 1,499 different sgRNAs targeting 2,111 out of 2,146 genetic elements of *S. pneumoniae*. The resulting library was grown in presence or absence of aTc to repress, or not, every operon, and in presence or absence of IPTG to express or deplete *ccrZ*, and the sgRNAs were then sequenced by Illumina sequencing. If an operon becomes more essential in a *ccrZ*-depletion background than in an induced *ccrZ* background, the corresponding sgRNA will therefore be under-represented. After analyzing the fold change for every sgRNA between *ccrZ-*depletion and *ccrZ*-complementation, we found an enrichment of sgRNAs targeting the operon of YabA / HolB (*tmk-holB-yabA-spv_0828*), indicating that depletion of this operon becomes beneficial for growth of Δ*ccrZ* (Fig. 4m and Supplementary Table 4). This confirms that deletion of YabA can complement a *ccrZ* deletion. Interestingly, we also found that inactivation of two genes coding for FtsK and RocS, worsened the fitness of a *ccrZ* mutant. RocS is a regulator of chromosome segregation in *S. pneumoniae* interacting with the DNA and the chromosome partitioning protein ParB^50^ and FtsK is thought to couple segregation of the chromosome terminus during cell division^51^. These interactions reinforce a role of CcrZ in chromosome integrity and replication and that CcrZ acts in a distinct pathway from these chromosome segregation factors. Finally, to test whether the midcell localization of CcrZ is important for well-timed replication of the chromosome, we abrogated CcrZ’s mid-cell localization by depleting cells for FtsZ (Fig. 3e). After 2h of FtsZ depletion, chromosomal DNA was isolated and *oriC*/*ter* ratios were determined. This showed that upon mis-localization of CcrZ, cells under replicate (Fig. 4n).

### CcrZ is a conserved regulator of DnaA

The results so far suggest that the division defects observed in the absence of CcrZ are due to perturbed Z-ring formation caused by under-replication of the chromosome. To examine whether disruption of DNA replication in general could lead to division defects similar to those of a *ccrZ* mutant, we took advantage of a thermosensitive *dnaA* mutant (*dnaA^TS^*) in which DNA replication initiation is drastically reduced when cells are grown at the non-permissive temperature (40°C)^50^. As expected, when shifted to the non-permissive temperature, many cells were anucleate (Extended Data Fig. 4a). Strikingly, localization of FtsZ-mTurquoise2 in the *dnaA^TS^* strain at 40°C phenocopied the Δ*ccrZ* mutant, and FtsZ was frequently mis-localized (Fig. 5a). Examination by time-lapse microscopy following a temperature shift from 30°C to 40°C showed that FtsZ-mTurquoise2 mis-localization occurs after four to five generations (Supplementary Video 7). Furthermore, examination by TEM at 40°C showed many cells with aberrant septa like CcrZ-depleted cells (Fig. 5b). As DnaA inactivation leads to strikingly similar phenotypes, these data are consistent with the idea that CcrZ exerts a control on DNA replication initiation.

**Fig. 5:**
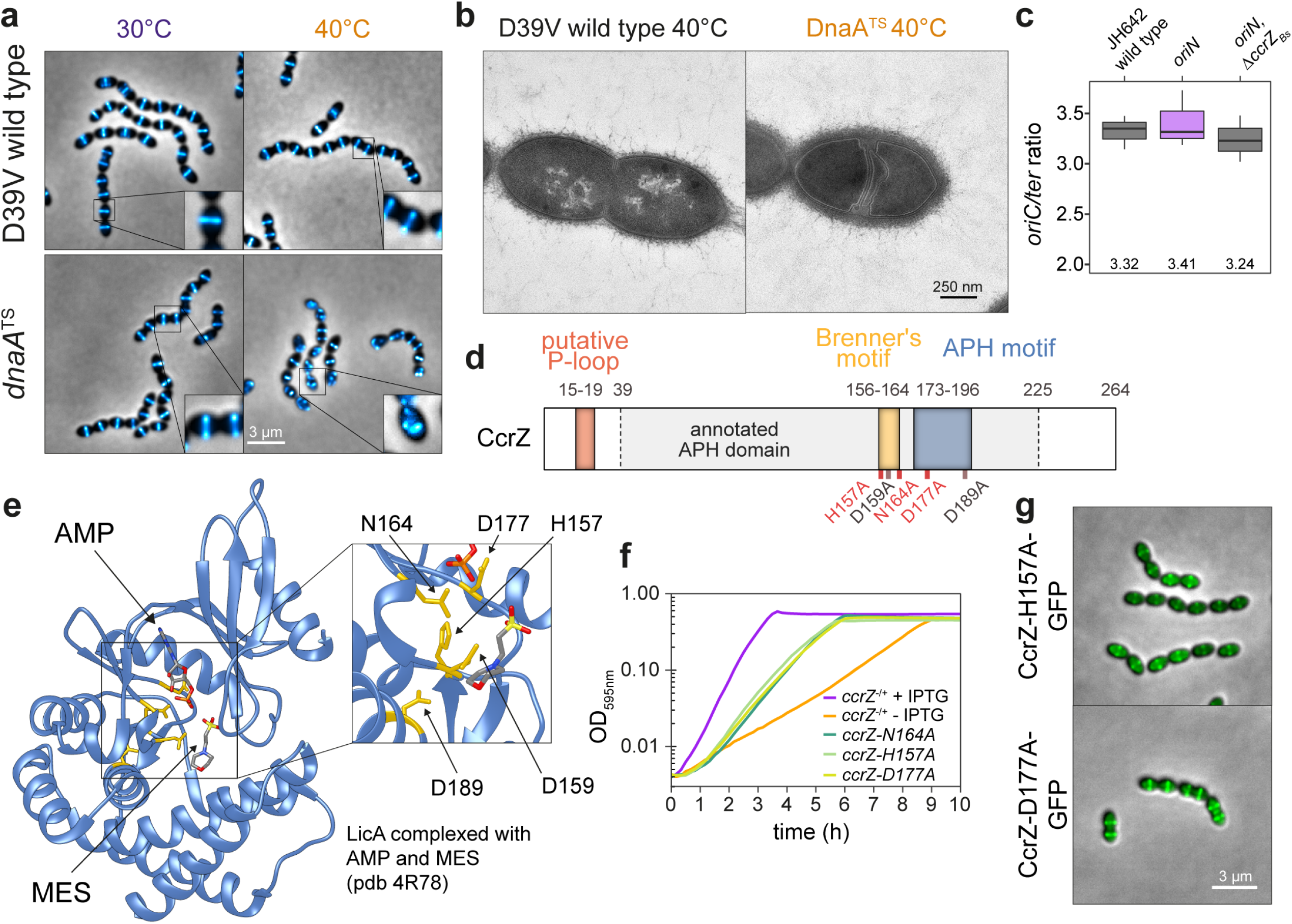
CcrZ activates DnaA-dependent replication initiation. **a**, Localization of FtsZ-mTurquoise2 in a thermo-sensitive DnaA strain (*dnaA^TS^*) at permissive (30°C) and non-permissive (40°C) temperatures shows that *dnaA* inactivation leads to a similar phenotype as *ccrZ* inactivation. **b**, TEM of DnaA^TS^ at non-permissive temperature (40°C) indicates the presence of multiple septa, similarly to a Δ*ccrZ* mutant. **c**, When replication is driven in a RepN-dependent manner in *B. subtilis* (*oriN*), no decrease in *ori/ter* ratio can be observed in absence of *ccrZ_Bs_* (*oriN,* Δ*ccrZ_Bs_*). **d**, Schematic representation of CcrZ motifs. CcrZ has one putative domain annotated APH (Phosphotransferase enzyme family; PFAM01636). Sequence alignment with several kinases revealed the presence of a conserved P-loop, APH and Brenner’s motifs, found in most phosphotransferases. Locations of mutations made for 3 essential (red) and 2 non-essential (black) conserved residues are shown underneath. **e**, LicA choline kinase structure complexed with AMP and MES (2-(N-morpholino)ethanesulfonic acid). The 5 residues indicated in yellow are conserved between CcrZ and LicA (and highly conserved within Firmicutes). **f**, Mutation of three of these five conserved residues in the putative ATP binding pocket leads to growth defects. **g**, Localization of CcrZ-H157A-GFP and CcrZ-D177A-GFP is not impaired.

To test whether CcrZ controls DNA replication via regulating DnaA activity, we made use of the fact that a *B. subtilis* Δ*ccrZ_Bs_* mutant also under-initiates (Fig. 4g) and a strain was constructed in which DNA replication was driven in a RepN-dependent manner (from a plasmid origin of replication *oriN*) rather than from DnaA-dependent initiation (from *oriC*). This showed no significant *ori-ter* ratio differences when *ccrZ* was deleted (Fig. 5c), suggesting that CcrZ is an activator of DnaA-dependent initiation of replication in *B. subtilis*. We therefore tested whether CcrZ interacts directly with DnaA to trigger DNA replication and employed bacterial two-hybrid assays and the Split-luc system using pneumococcal CcrZ and DnaA (Fig. 3a and Extended Data Fig. 4b). However, none of these assays revealed a direct protein-protein interaction. In line with our genetic data, we also did not find a direct interaction of CcrZ with YabA, while YabA clearly interacts with DnaA (Extended Data Fig. 4c). It is still possible that CcrZ interacts directly with DnaA, but that we cannot detect it with these assays. Alternatively, another factor might be required for CcrZ’s function or CcrZ indirectly affects the activity of DnaA in replication initiation.

### CcrZ’s conserved residues are essential for its function

*S. pneumoniae* CcrZ is 264 amino acids long and is predicted to have a single APH (aminoglycoside phosphotransferase enzyme family) domain (Fig. 5d). Sequence alignment using Psi-BLAST showed homology with phosphotransferase enzyme family proteins, while pairwise comparisons of profile-hidden Markov models (HMMs) using HHpred^52^ identified homologies with ethanolamine- and choline kinases. Despite various attempts, we have not been able to establish any biochemical activity or nucleotide binding for recombinant purified CcrZ and were unable to produce protein crystals, probably because of its rapid precipitation in solution. Nevertheless, as CcrZ is highly conserved in Firmicutes, we aligned CcrZ protein sequence with 1000 protein sequences from UniRef50 and identified three residues conserved in more than 95% of the proteins (D159, N164 and D177) and two other residues (H157 and D189) in more than 80% (Fig. 5d and Extended Data Fig. 4d). To determine the position of these residues, the *S. pneumoniae* CcrZ protein sequence was mapped onto the crystal structure of the best hit from the HMM alignment, the choline kinase LicA, in complex with adenosine monophosphate (AMP) (pdb 4R78). Interestingly, these five conserved residues appear to be in spatial proximity to AMP and thus to a putative nucleotide-binding pocket (Fig. 5e). Comparison of CcrZ and LicA sequences shows a conserved Brenner’s motif [HXDhX3N] (residues CcrZ H157 – N164) found in most phosphotransferases (Fig. 5d). In this motif, LicA-N187 (CcrZ-N164) was shown to interact with the α-phosphate moiety of AMP^53^ and LicA-D176 (CcrZ-D159) was shown to be crucial for hydrogen bond formation with the hydroxyl moiety of choline. Furthermore, it also has a conserved motif found in phosphotranspherases (APH), in which LicA-D194 (CcrZ-D177) was shown to interact with the α-phosphate moiety of AMP. CcrZ-D189 corresponds to residue D313 of the choline kinase A (cka-2) of *Caenorhabditis elegans*, a residue which was proposed to stabilize the cka-2 dimer as well as the catalytic site^54^. CcrZ however does not possess the conserved hydrophobic residues specific to choline- and ethanolamine-kinases necessary for choline binding, but instead has several polar amino acids at these positions (*e.g.,* the crucial residues LicA-Y197 and V178 corresponding to CcrZ-S180 and R161). Mutational analysis of the five conserved residues of CcrZ showed that at least H157, N164 and D177 are essential for CcrZ’s function in *S. pneumoniae* (Fig. 5f), while mutating CcrZ-D159 or CcrZ-D189 did not lead to any growth defect. All three essential mutants were properly produced (Extended Data Fig. 1c) and CcrZ-H157A and CcrZ-D177A could still localize at the septum (Fig. 5g). Therefore, these three residues are crucial for the function of CcrZ. It is interesting to note that none of the three mutants was dominant negative when expressed together with a wild type CcrZ. Given the high similarity with LicA, it is very likely that CcrZ can bind an as of yet unknown nucleotide.

### A model for CcrZ-controlled DNA replication in *S. pneumoniae*

We showed above that CcrZ is fundamental for DnaA-dependent DNA replication initiation in *B. subtilis* and that *S. pneumoniae* CcrZ localizes at mid-cell for most of the cell cycle. In *S. pneumoniae*, once DNA replication initiates at mid-cell, the origins localize at both future division sites, while the replication machinery stays near the Z-ring until completion of replication and closure of the septum^35^. We therefore hypothesized that CcrZ is brought to mid-cell by FtsZ to promote initiation of DNA replication. To map the hierarchy of events that take place during the pneumococcal cell cycle, we constructed a triple-labeled strain (strain *ccrZ-mKate2 dnaN-sfTQ^OX^ parB_p_-mYFP*) in which CcrZ is fused to a red fluorescent protein, DNA replication is visualized by a DnaN fusion to a cyan fluorescent protein, and the origin of replication is marked with a yellow fluorescent reporter (see Methods). Imaging of this strain by short time-lapse fluorescence microscopy revealed that DNA replication initiates once CcrZ is assembled at mid-cell, rapidly followed by segregation of the newly replicated origins as cells elongate (Fig. 6a-d and Supplementary Video 8). The replication machinery remains near the old division site together with CcrZ, only to move to the new cell division sites once DNA replication is complete. This data supports a model in which FtsZ brings CcrZ to *oriC* to stimulate DnaA to fire a new round of replication ensuring that DNA replication only commences after the origins are well segregated and positioned at the new mid-cell position. Indeed, DnaA co-localizes with CcrZ in new-born cells (Extended Data Fig. 5). In the absence of CcrZ, initiation of DNA replication is mis-timed and occurs too late relative to cellular growth and Z-ring formation, frequently leading to futile division events, mis-segregated chromosomes and anucleate cells (Fig. 6e).

**Fig. 6:**
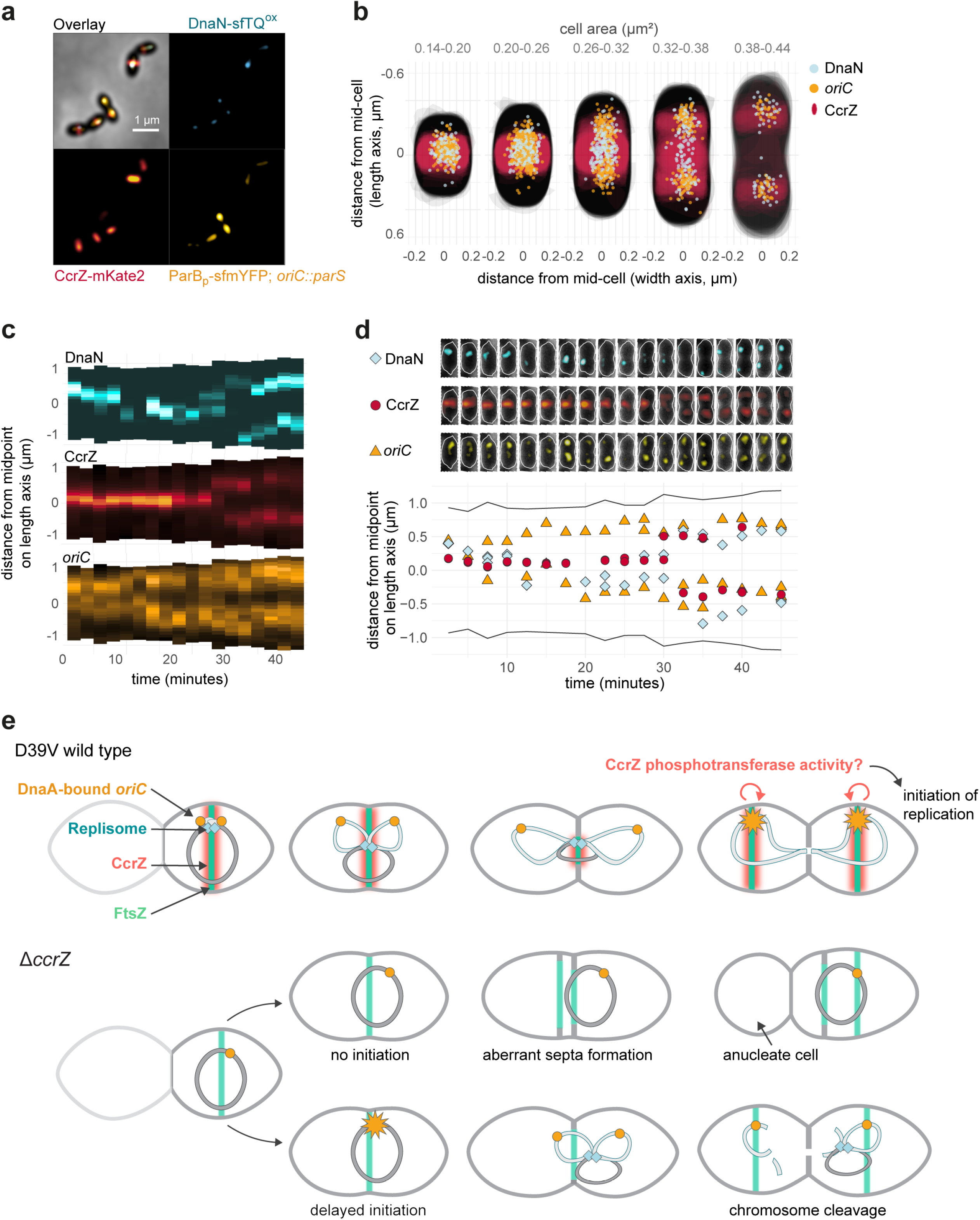
Spatio-temporal localization of CcrZ via FtsZ ensures proper timing of DNA replication in *S. pneumoniae*. **a**, Microscopy of the origin of replication (yellow), replication fork (cyan) and CcrZ (red) in live *S. pneumoniae* wild type background cells. **b**, DnaN, *oriC* and CcrZ localizations grouped by cell area (µm^2^) in five equally sized groups. Analyzed from snapshots of exponentially growing cells. **c**, Single-cell kymographs of DnaN, CcrZ and *oriC* localizations in a 2:30 minute interval time-lapse movie. **d**, Tracked DnaN, *oriC* and CcrZ over time in a single cell. Top: overlay of fluorescence, cell outline and phase-contrast of the cell displayed in panel c and bottom: fluorescence localization on the length axis of the same single cell over time. **e**, Model for spatio-temporal control of replication by CcrZ. In *S. pneumoniae*, CcrZ is brought to the middle of the cell where the DnaA-bound origin of replication is already positioned. CcrZ then stimulates DnaA to trigger DNA replication by an as of yet unknown activity, possibly involving a phosphor-transfer event. While the precise regulation and localization of CcrZ seems diverse between different organisms, CcrZ’s activity to stimulate DNA replication is conserved, at least in *S. pneumoniae, S. aureus* and *B. subtilis*.

## Discussion

The principal contribution of this work is the identification and initial functional characterization of a new mechanism for cell cycle regulation in *S. pneumoniae* via the CcrZ protein. We show that CcrZ’s septal localization occurs via a direct interaction with FtsZ. Our data is consistent with a model in which, once positioned at mid-cell where the DnaA-bound origin of replication is located, CcrZ stimulates DnaA, likely by phosphorylation of an intermediate molecule, to initiate DNA replication (Fig. 6). Importantly, CcrZ’s function of controlling DnaA seems conserved in *S. aureus* and *B. subtilis*, and likely in many other Gram-positive bacteria (Extended data Fig. 1a).

Besides the production of anucleate cells and cells with cleaved chromosomes, *ccrZ* mutants contain multiple aberrant division septa (Fig. 6e). Notably, this is phenocopied by a temperature sensitive DnaA allele. This indicates that chromosome replication itself, and correct localization of the chromosome has an important role in nucleoid occlusion: when initiation is too late and the new daughter chromosomes are not fully segregated, division can take place over the DNA resulting in dissected chromosomes. We also observed multiple division septa in cells depleted for CcrZ that are likely caused by mis-timed chromosome segregation whereby Z-rings are formed adjacent to the nucleoid. These phenotypes are reminiscent of observations made for *E. coli* and *B. subtilis* that showed that after arrest of DNA replication, many cells continued to elongate without dividing, but FtsZ rings continued to form and almost always were located to the side of nucleoids^55,56^. In this respect, it is interesting to note that the *S. aureus* Noc system also controls DNA replication, as Δ*noc* cells over-initiate DNA replication^57^. In support of our findings, a lethal Δ*noc* Δ*comEB* double mutant in *S. aureus* could be rescued by a suppressor mutation in *ccrZ_Sa_*, further indicating that CcrZ_Sa_ is also involved in the control of DNA replication in *S. aureus*.

This work uncovers a novel mechanism in which a single protein links cell division with DNA replication control. In this model, Z-ring formation is used as a timer for the initiation of DNA replication. When cell division terminates, leading to the formation of another Z-ring at the new division site, CcrZ is brought along and can activate a new round of DNA replication. This simple system ensures that DNA replication only commences a single time per cell cycle in newborn cells. It will be interesting to see how CcrZ controls the cell cycle in other bacteria, what the involved biochemical activities are and whether CcrZ will prove as a new target for innovative antibiotics.

## Methods

### Bacterial strains and culture conditions

All strains, plasmids and primers used are listed in Supplementary Table 1 and Supplementary Table 2.

All pneumococcal strains in this study are derivate of *S. pneumoniae* D39V^58^, unless specified otherwise, and are listed in Supplementary Table 1. Strains were grown in liquid semi-defined C+Y medium^59^ at 37°C from a starting optical density (OD_600nm_) of 0.01 until the appropriate OD. Induction of the zinc-inducible promoter (*P_Zn_*) was carried out by supplementing the medium with 0.1 mM ZnCl_2_ and 0.01 mM MnCl_2_ (Sigma-Aldrich) and the IPTG-inducible promoter (*P_lac_*) was activated with 1 mM IPTG (β-D-1-thiogalactopyranoside, Sigma-Aldrich). For all related experiments, depletion strains where first grown without inducer until OD_600nm_ = 0.3 and then diluted 100 times in fresh medium and grown until the desired OD. Transformation of *S. pneumoniae* was performed as described before^59^ with cells taken at exponential growth phase (OD_600nm_ = 0.1). When necessary, the medium was supplemented with the following antibiotics: chloramphenicol (0.45 µg.mL^−1^), erythromycin (0.2 µg.mL^−1^), kanamycin (250 µg.mL^−1^), spectinomycin (200 µg.mL^−1^) and tetracycline (0.5 µg.mL^−1^).

*S. aureus* strains are listed in Supplementary Table 1. Cells were grown in brain heart infusion (BHI) medium (Oxoid) with shaking at 37°C. When appropriate, 5 µg.mL^−1^ erythromycin and / or 10 µg.mL^−1^ chloramphenicol was added to the growth medium. All *S. aureus* plasmids were initially made in *E. coli* strain IM08B^60^. *E. coli* IM08B was grown in LB medium at 37°C with shaking; 100 µg.mL^−1^ ampicillin was added when appropriate. Plasmids were then transformed into *S. aureus* by electroporation, as described previously^61^.

*B. subtilis* strains are listed in Supplementary Table 1. Cells were grown with shaking at 37°C in Luria-Bertani (LB) medium or S7 defined minimal medium with MOPS (3-(N-morpholino) propanesulfonic acid) buffer at a concentration of 50 mM rather than 100 mM supplemented with 1 % glucose, 0.1 % glutamate, trace metals, 40 μg.mL^−1^ phenylalanine, and 40 μg.mL^−1^ tryptophan^62^. Standard concentrations of antibiotics were used when appropriate. *B. subtilis* strains were derived from 1A700 or JH642 (*pheA1 trpC2*)^63^.

### Strain construction

Construction of strains is described in the Supplementary Methods.

### Microtiter plate-based growth assay

For *S. pneumoniae* growth assays, cells were first grown in C+Y medium pH = 7.4 until mid-exponential growth phase (OD_595nm_ = 0.3) with no inducer at 37°C, after which they were diluted 100 times in fresh C+Y medium supplemented with IPTG or ZnCl_2_ when appropriate. Cellular growth was then monitored every 10 min at either 37°C or 30°C in a microtiter plate reader (TECAN Infinite F200 Pro). Each growth assay was performed in triplicate. The lowest OD_595nm_ of each growth curve was normalized to 0.004 (detection limit of the reader and initial OD_595nm_ of the inoculum) and the average of the triplicate values were plotted, with the SEM (Standard Error of the Mean) represented by an area around the curve.

For assessment of *S. aureus* growth, CRISPRi knockdown strains were grown overnight in BHI medium. Cultures were then diluted 100-fold and grown until OD_600nm_ = 0.4. The cultures were then re-diluted 200-fold in medium with or without inducer 500 µM IPTG. Growth analysis was performed on a Synergy H1 Hybrid (BioTek) microtiter plate reader at 37°C with measurement of OD_600nm_ every 10 min. Average of the triplicate values were plotted, with the SEM (Standard Error of the Mean) represented by an area around the curve.

### Phase contrast and fluorescence microscopy

*S. pneumoniae* cells were grown in C+Y medium pH = 7.4 at 37°C to an OD_595nm_ = 0.1 without any inducer and diluted 100 times in fresh C+Y medium supplemented when appropriate with IPTG (for activation of dCas9, complementation of CcrZ and FtsZ, or expression of fluorescent fusions) or ZnCl_2_ (for CcrZ complementation or expression of fluorescent fusions). At OD_595nm_ = 0.1, 1 mL of culture was harvested by centrifugation 1 min at 9,000 x g. For DAPI staining, 1 µg.mL^−1^ DAPI (Sigma-Aldrich) was added to the cells and incubated for 5 min at room temperature prior to centrifugation. For imaging of bulk exponentially growing cultures, cells were washed twice with 1 mL ice-cold PBS and re-suspended into 50 µL ice-cold PBS; for time-lapse microscopy, cells were washed and re-suspended into 1 mL of fresh pre-warmed C+Y medium. 1 µL of cells were then spotted onto PBS- or C+Y-polyacrylamide (10 %) pads. For time-lapse microscopy, pads were incubated twice for 30 min in fresh C+Y medium at 37°C prior to spotting. Pads were then placed inside a gene frame (Thermo Fisher Scientific) and sealed with a cover glass as described before^64^. Microscopy acquisition was performed either using a Leica DMi8 microscope with a sCMOS DFC9000 (Leica) camera and a SOLA light engine (Lumencor), or using a DV Elite microscope (GE Healthcare) with a sCMOS (PCO-edge) camera and a DV Trulight solid state illumination module (GE Healthcare), and a 100x/1.40 oil-immersion objective. Phase contrast images were acquired using transmission light (100 ms exposure). Still fluorescence images were usually acquired with 700 ms exposure, and time-lapses with 200-300 ms exposure. Leica DMi8 filters set used are as followed: DAPI (Leica 11533333, Ex: 395/25 nm, BS: LP 425 nm, Em: BP 460/50 nm), CFP (Ex: 430/24 nm Chroma ET430/24x, BS: LP 455 Leica 11536022, Em: 470/24 nm Chroma ET470/24m), GFP (Ex: 470/40 nm Chroma ET470/40x, BS: LP 498 Leica 11536022, Em: 520/40 nm Chroma ET520/40m), YFP (Ex: 500/20 nm Chroma ET500/20x, BS: LP 520 Leica 11536022, Em: 535/30 nm Chroma ET535/30m) and mCherry (Chroma 49017, Ex: 560/40 nm, BS: LP 590 nm, Em: LP 590 nm). DeltaVision microscope used a DV Quad-mCherry filter set: GFP (Ex: 475/28 nm, BS: 525/80 nm, Em: 523/36 nm) and mCherry (Ex: 575/25 nm, BS: 605/50, Em: 632/60 nm). Images were processed using either LAS X (Leica) or SoftWoRx (GE Healthcare). For *S. aureus* microscopy, cells were induced as described above, grown until OD_600nm_ = 0.2 and analyzed on a Zeiss AxioObserver with an ORCA□Flash4.0 V2 Digital CMOS camera (Hamamatsu Photonics) through a 100x PC objective. HPX 120 Illuminator (Zeiss) was used as a light source for fluorescence microscopy. Images were processed using ZEN (Zeiss). Signals was deconvolved, when appropriate, using Huygens (SVI) software.

### Transmission Electron Microscopy (TEM)

Strains were grown in C+Y medium at either 37°C, or at 30°C for *dnaA^TS^,* until an OD_595nm_ = 0.3, with or without addition of ZnCl_2_ (for *ccrZ* complementation or depletion, respectively) and diluted 100 times into 10 mL of fresh C+Y medium. Cells were then grown either at 37°C or at 40°C, for *dnaA* depletion in the *dnaA^TS^* strain, until OD_595nm_ = 0.15. 5 mL of each sample was then fixed with 2.5 % glutaraldehyde solution (EMS) in phosphate buffer (PB 0.1 M pH = 7.4) (Sigma Aldrich) for 1h at room temperature, followed by 16 h incubation at 4°C. Cells were then post-fixed by a fresh mixture of osmium tetroxide 1 % (EMS) with 1.5 % potassium ferrocyanide (Sigma Aldrich) in PB buffer for 2 h at room temperature. Samples were then washed three times with distilled water and spun down in low melting agarose 2 % (Sigma Aldrich) and solidified in ice. Solid samples were then cut in 1 mm^3^ cubes and dehydrated in acetone solution (Sigma Aldrich) at graded concentrations (30 % for 40 min; 50 % for 40 min; 70 % for 40 min and 100 % for 3 x 1 h). This step was followed by infiltration in Epon (Sigma Aldrich) at graded concentrations (Epon 1/3 acetone for 2 h; Epon 3/1 acetone for 2 h, Epon 1/1 for 4 h and Epon 1/1 for 12 h) and finally polymerized for 48 h at 60°C. Ultra-thin sections of 50 nm were then cut on a Leica Ultracut (Leica Mikrosysteme GmbH) and placed on a copper slot grid 2 x 1 mm (EMS) coated with a polystyrene film (Sigma Aldrich). Sections were subsequently post-stained with 4 % uranyl acetate (Sigma Aldrich) for 10 min, rinsed several times with water, then with Reynolds lead citrate (Sigma Aldrich) for 10 min and rinsed several times with distilled water. Micrographs were taken using a transmission electron microscope Philips CM100 (Thermo Fisher Scientific) equipped with a TVIPS TemCam-F416 digital camera (TVIPS) and using an acceleration voltage of 80 kV. Number of septa and cell length were manually measured on TEM images of cells in the correct focal plane: 22 wild type cells, 28 CcrZ-depleted cells and 17 CcrZ-complemented cells.

### 3D Structured Illumination Microscopy (3D-SIM)

Samples for 3D-SIM were prepared as described previously by spotting 1 µL onto PBS-10 % acrylamide pads. Acquisition was performed on a DeltaVision OMX SR microscope (GE Healthcare) equipped with a 60x/1.42 NA objective (Olympus) and 488 nm and 568 nm excitation lasers. 16 Z-sections of 0.125 μm each were acquired in Structure Illumination mode with 20 ms exposure and 20 % laser power. The 240 images obtained were reconstructed with a Wiener constant of 0.01, and the volume reconstructed using SoftWoRx.

### Image analysis and cells segmentation

All microscopy images were processed using Fiji (fiji.sc). Cell segmentation based on phase contrast images was performed either on Oufti^65^, MicrobeJ^66^ or Morphometrics^67^ and fluorescent signals where analyzed using Oufti (for CcrZ and FtsZ), MicrobeJ^66^ (for CcrZ) or iSBatch^68^ (for DnaN and *oriC*). Fluorescence heat-maps were generated using BactMAP^69^.

### Small-scale expression and GFP resin pull-down of FtsZ and CcrZ-GFP

For affinity purification of CcrZ_Sp_-GFP while expressing FtsZ_Sp_, *ccrZ_Sp_* was amplified from D39V genomic DNA with primers 213/214 and the resulting fragment was assembled using Golden Gate allelic replacement strategy (BsaI) with plasmid pET-Gate2 ccdB (pSG436), pSG366, pSG367 and pSG2562, resulting in plasmid pSG2950. *ftsZ* was amplified by PCR 215/216 on D39V genomic DNA and cloned into plasmid pJet1.2, resulting in plasmid pSG4227. The later was then assembled with pSG1694 using Golden Gate assembly, leading to plasmid pSG4268. BL21 DE3 Gold competent cells were co-transformed with plasmids containing one of each *S. pneumoniae* FtsZ and CcrZ-GFP. Expression was ZYM-5052 autoinduction media^70^. Cells were sonicated in buffer containing 50 mM Tris pH 7.5, 150 mM potassium acetate, 5 % glycerol, and 5 mM β-mercaptoethanol (lysis buffer). Supernatant was then mixed with GFP resin which was produced by crosslinking nanobody^71^ to NHS-Activated Sepharose 4 Fast Flow beads (GE Healthcare) according to the manufacturer’s instructions. After 1 hour of batch binding, resin was washed 10 column volume (CV) with lysis buffer. Beads were then re-suspended in 50 µL of lysis buffer mixed with SDS-PAGE loading dye containing 5 % w/v β-mercaptoethanol and heat treated at 95 □C for 15 minutes. Supernatant was collected and labelled heat elution (HE) samples. Whole cell lysate (WC), supernatant after sonication (S), and HE were loaded on 15 % SDS-PAGE gels and visualized by Coomassie staining.

### Large-scale purification of CcrZ-CPD for antibody production

In order to express a fusion of *S. pneumoniae* CcrZ with a C-terminal cysteine protease domain (CPD), *ccrZ* was amplified by PCR from D39V genomic DNA with primers 213/214 and assembled using Golden Gate allelic replacement strategy (BsaI) with plasmid pET-Gate2 ccdB (pSG436), pSG366, pSG367 and pSG2559. The resulting pSG2949 plasmid was then transformed into BL21 DE3 Gold cells using ZYM-5052 auto-induction media^70^. Cells were sonicated in buffer containing 300 mM NaCl, 50 mM Tris pH 7.5, 5 mM β-mercaptoethanol, and protease inhibitor cocktail (PIC). Supernatant was loaded onto a gravity flow column containing HisPur^TM^ Cobalt Resin (Thermo Scientific). Column was washed 5 CV with buffer containing 100 mM NaCl, 20 mM Tris pH 7.5, 5 mM β-mercaptoethanol. Because CcrZ had affinity to the resin even without the CPD, instead of on column tag cleavage, elution was collected with buffer containing 150 mM Imidazole, 100 mM NaCl, 20 mM Tris pH 7.5, 5 mM β-mercaptoethanol, and tag cleavage was performed for 1 hour at 4 □C by adding 1 mM inositol hexakisphosphate. The sample was further purified using a HitrapQ column and Superdex 200 16/600 pg column (GE). The final storage buffer contained 100 mM NaCl, 20 mM Tris pH 7.5, 1 mM DTT. For antibody production, sample was loaded onto a 15 % SDS PAGE gel. Edge wells were cut out and stained with Coomassie to determine position of CcrZ on the gel. Gel portions containing CcrZ was sent for antibody production by Eurogentec.

### Western blot analysis

Cells were grown in C+Y medium until OD_595nm_ = 0.2 and harvested by centrifugation at 8000 x g for 2 min at room temperature from 1 mL of culture. Cells were re-suspended into 150 µL of Nuclei lysis buffer (Promega) containing 0.05 % SDS, 0.025% deoxycholate and 1 % Protease Inhibitor Cocktail (Sigma Aldrich), and incubated at 37°C for 20 min and at 80°C for 5 min in order to lyse them. One volume of 4X SDS sample buffer (50 mM Tris-HCl pH = 6.8, 2 % SDS, 10 % glycerol, 1 % β-mercaptoethanol, 12.5 mM EDTA and 0.02 % Bromophenol blue) was then added to three volumes of cell lysate sample and heated at 95°C for 10 min. Protein samples were separated by SDS–PAGE (4-20%) and blotted onto polyvinylidene fluoride membranes (Merck Millipore). Membranes were blocked for 1 h with Tris-buffered saline (TBS) containing 0.1 % Tween 20 (Sigma Aldrich) and 5 % dry milk and further incubated for 1 h with primary antibodies diluted in TBS, 0.1 % Tween 20, 5 % dry milk. Polyclonal CcrZ-antiserum concentration used was 1:5000 and commercial monoclonal GFP-IgG (Thermo Fisher Scientific) were used at 1:5000. Membranes were washed four times for 5 min in TBS, 0.1 % Tween 20 and incubated for 1 h with the secondary IgG (HRP-conjugated donkey anti-rabbit antibodies, Promega) diluted 1:20,000 in TBS, 0.1 % Tween 20 and 5 % dry milk. Membranes were then washed four times for 5 min in TBS, 0.1 % Tween 20 and revealed with Immobilon Western HRP substrate (Merck Millipore).

### ccrZ-GFP purification with anti-GFP nanobodies

*gfp-ccrZ* and *P3-gfp* (negative control) strains were grown in C+Y medium at 37°C until OD_595nm_ = 0.2 and cells were harvested by centrifugation 15 min at 3000 x g at 4°C. Cells were then incubated in sucrose buffer (0.1 M Tris-HCl pH = 7.5, 2 mM MgCl_2_, 1 M sucrose, 1 % Protease Inhibitor Cocktail (Sigma Aldrich), 200 μg.mL^−1^ RNase A and 10 μg.mL^−1^ DNase (Sigma Aldrich)) for 30 min at 30°C, then incubated in hypotonic buffer (0.1 M Tris-HCl pH = 7.5, 1 mM EDTA, 1 % Triton, 1 % Protease Inhibitor Cocktail, 200 μg.mL^−1^ RNase A and 10 μg.mL^−1^ DNase) for 15 min at room temperature and cell debris were eliminated by centrifugation 30 min at 15,000 x g at 4°C. Cell lysate was then incubated with equilibrated GFP-Trap resin (Chromotek) at 4°C for 2 h. After several washes with wash buffer (10 mM Tris-HCl pH = 7.5, 150 mM NaCl, 0.5 mM EDTA, 1 % Protease Inhibitor Cocktail), beads were resuspended in 20 µL 8 M Urea, 50 mM triethylammonium bicarbonate (TEAB), pH = 8.0 and reduced with 5 mM DTT for 30 min at 37°C. Cysteines were alkylated by adding 20 mM iodoacetamide and incubated for 30 min at room temperature in the dark. Samples were diluted 1:1 with TEAB buffer and digested by adding 0.1 µg of modified Trypsin (Promega) and incubated overnight at 37°C, followed by a second digestion for 2 h with the same amount of enzyme. The supernatant was collected, diluted 2 times with 0.1 % formic acid and desalted on strong cation exchange micro-tips (StageTips, Thermo Fisher scientific) as described^72^. Peptides were eluted with 1.0 M ammonium acetate (100 µL). Dried samples were resuspended in 25 µL 0.1 % formic acid, 2 % acetonitrile prior being subjected to nano LC-MS/MS.

### LC-MS/MS analysis

Tryptic peptide mixtures (5 µL) were injected on a Dionex RSLC 3000 nanoHPLC system (Dionex, Sunnyvale, CA, USA) interfaced via a nanospray source to a high resolution QExactive Plus mass spectrometer (Thermo Fisher Scientific). Peptides were separated on an Easy Spray C18 PepMap nanocolumn (25 or 50 cm x 75 µm ID, 2 µm, 100 Å, Dionex) using a 35 min gradient from 4 to 76 % acetonitrile in 0.1 % formic acid for peptide separation (total time: 65 min). Full MS survey scans were performed at 70,000 resolution. In data-dependent acquisition controlled by Xcalibur software (Thermo Fisher), the 10 most intense multiply charged precursor ions detected in the full MS survey scan were selected for higher energy collision-induced dissociation (HCD, normalized collision energy NCE = 27 %) and analysis in the orbitrap at 17,500 resolution. The window for precursor isolation was of 1.6 m/z units around the precursor and selected fragments were excluded for 60 sec from further analysis.

MS data were analyzed using Mascot 2.5 (Matrix Science, London, UK) set up to search the UniProt (www.uniprot.org) protein sequence database restricted to *S. pneumoniae* D39 / NCTC 7466 taxonomy (339 SWISSPROT sequences + 1586 TrEMBL sequences). Trypsin (cleavage at K,R) was used as the enzyme definition, allowing 2 missed cleavages. Mascot was searched with a parent ion tolerance of 10 ppm and a fragment ion mass tolerance of 0.02 Da (QExactive Plus). Iodoacetamide derivative of cysteine was specified in Mascot as a fixed modification. N-terminal acetylation of protein, oxidation of methionine and phosphorylation of Ser, Thr, Tyr and His were specified as variable modifications. Scaffold software (version 4.4, Proteome Software Inc., Portland, OR) was used to validate MS/MS based peptide and protein identifications, and to perform dataset alignment. Peptide identifications were accepted if they could be established at greater than 90.0 % probability as specified by the Peptide Prophet algorithm^73^ with Scaffold delta-mass correction. Protein identifications were accepted if they could be established at greater than 95.0 % probability and contained at least 2 identified peptides. Protein probabilities were assigned by the Protein Prophet algorithm^74^. Proteins that contained similar peptides and could not be differentiated based on MS/MS analysis alone were grouped to satisfy the principles of parsimony. Proteins sharing significant peptide evidence were grouped into clusters.

### Split luciferase assay

*S. pneumoniae* cells were grown in C+Y medium at 37°C until OD_595nm_ = 0.2 and washed once with fresh C+Y medium. 1 % NanoGlo Live Cell substrate (Promega) was then added, and luminescence was measured 20 times at 37°C every 30 sec in plate reader (TECAN Infinite F200 Pro). Measurements were performed in triplicate and the average values were plotted, with the SEM (Standard Error of the Mean) represented by the dot size.

### Bacterial two-hybrid assay

The bacterial two-hybrid assay was based on the method from Karimova *et al*.^43^ with the following modifications. *dnaA*, *ccrZ* and *ftsZ* genes from *S. pneumoniae* D39V were cloned both into the low copy-number vector pUT18 and into the high copy-number vector pST25^75^ using the enzymes BamHI and KpnI. *Escherichia* coli strain HM1784 (BTH101 Δ*rnh*::*kan*) was transformed using each combination of plasmids. Chemically competent cells were incubated on ice for 60 min, heat shocking at 42□ for 90 sec and then inoculated at 37□ in 3 mL of LB media supplemented with ampicillin (100 µg/mL) and spectinomycin (50 µg/mL) with mild agitation for 16 hours. The A_600 nm_ was adjusted to 0.5, cultures were diluted 1:1000 and a 5 µL aliquot was spotted on a nutrient agar plate containing antibiotics (as above) containing 0.006 % X-gal. Plates were incubated at 30□ for 48 hours and the images were captured using a digital camera.

### Co-immunoprecipitation of CcrZ and FtsZ-GFP with anti-GFP nanobodies

*S. pneumoniae* cells were grown in C+Y medium at 37°C until OD_595nm_ = 0.2 and harvested by centrifugation 15 min at 3,000 x g at 4°C. Cells were lysed using GFP-Trap_A Lysis buffer (Chromotek), 0.25 % Deoxycolate, 1 % Protease Inhibitor Cocktail incubated at room temperature for 10 min followed by incubation at 4°C for 20 min. Cell lysate was incubated with equilibrated GFP-Trap resin (Chromotek) at 4°C for 2 h. The resin was then washed 3 times in GFP-Trap_A Wash buffer (Chromotek) and GFP-proteins were eluted using SDS sample buffer at 95°C for 10 min and analyzed by immunoblotting.

### Genome resequencing of ccrZ suppressors by NGS

Strains *hlpA-mKate2* Δ*ccrZ*, *ccrZ^supp1^, ccrZ^supp2^* and *ccrZ^supp3^* were grown in C+Y medium at 37°C until OD_595nm_ = 0.3 and cells were harvested by centrifugation 1 min at 10,000 x g. Pellet was then re-suspended into Nuclei lysis buffer (Promega) containing 0.05 % SDS, 0.025% deoxycholate and 200 µg.mL^−1^ RNase A at 37°C for 20 min to lyse the cells and Protein Precipitation Solution (Promega) was added. DNA was then precipitated using isopropanol. The extracted genomes were then analyzed by Illumina sequencing by GATC Biotech (Eurofins Genomics). Mutations were mapped onto D39V genome using breseq pipeline^76^. Genomes sequences are available at SRA (project PRJNA564501).

### oriC/ter ratios determination by RT-qPCR

Determination of *S. pneumoniae oriC/ter* ratios was performed as followed. Cells were pre-grown until OD_600nm_ = 0.4 in C+Y medium at 37°C, with or without inducer (ZnCl_2_ or IPTG) for complementation and depletion conditions, respectively. Cells were then diluted 100 times in fresh C+Y medium supplemented when appropriate with inducer and harvested for genomic DNA isolation when they reached OD_600nm_ = 0.1 (exponential phase). For normalization (*oriC/ter* ratio of 1), *dnaA* thermosensitive strain was grown for 2h at non-permissive temperature (40°C) in C+Y medium and harvested for chromosomal DNA isolation. As a negative (overinitiating) control, wild type *S. pneumoniae* was incubated 2h with 0.15 μg.mL^−1^ HPUra (DNA replication inhibitor) at 37°C prior to harvesting. Primers pairs OT1/OT2 and OT3/OT4 were used to amplify the *oriC* and *ter* regions respectively. Amplification by Real-Time qPCR was performed using SYBR Select Master Mix (Applied Biosystems) on a StepOne Plus Real-Time PCR System (Applied Biosystems), in triplicate. For *S. aureus oriC/ter* ratio determination, overnight cultures were diluted 100-fold and grown until OD_600nm_ = 0,4. These cultures were then re-diluted 200-fold in medium with 500 µM IPTG and grown until OD_600nm_ = 0.2. As reference samples with assumed *oriC/ter* ratio of 1, wild type *S. aureus* SH1000 cells at OD_600nm_ = 0.15 were supplemented with 50 µg.mL^−1^ rifampicin (inhibiting replication initiation) and incubated for 2 hours for replication run-out. Cells were then harvested and lysed enzymatically by addition of 0.2 mg.mL^−1^ lysostaphin and 10 mg.mL^−1^ lysozyme, and genomic DNA was isolated using the Wizard Genomic DNA Purification Kit (Promega). qPCR reactions of 10 µL were set up with 5 µL PowerUpTM SYBR^TM^ Green Master Mix (Applied Biosystems), 500 nM of each primer OT5/OT and OT7/OT8 and 20 ng of DNA. In both cases, amplification efficiencies of the primers and *oriC/ter* ratios were determined as described previously ^46^. Data were plotted as whiskers plot where whiskers represent the 10^th^ and 90^th^ percentile of data from Monte Carlo simulations, * P value < 0.05, significantly up. For *B. subtilis oriC/ter* ratios determination, cultures were grown to mid-exponential phase in LB medium and diluted back to OD_600nm_ = 0.05 and grown to mid-exponential phase (OD_600nm_ = 0.2 - 0.4) at 37°C. Cells were harvested in ice-cold methanol (1:1 ratio) and pelleted. Genomic DNA was isolated using Qiagen DNeasy kit with 40 μg.mL^−1^ lysozyme. The copy number of the origin (*oriC*) and terminus (*ter*) were quantified by qPCR to generate the *oriC/ter* ratio. qPCR was done using SSoAdvanced SYBR master mix and CFX96 Touch Real-Time PCR system (Bio-Rad). Primers used to quantify the origin region were OT9/OT10. Primers used to quantify the terminus region were OT11/OT12. Origin-to-terminus ratios were determined by dividing the number of copies (as indicated by the Cp values measured through qPCRs) of the origin by the number of copies quantified at the terminus. Ratios were normalized to the origin-to-terminus ratio of a temperature sensitive mutant, *dnaB134* (KPL69), that was grown to have synchronized replication initiation, resulting in 1:1 ratio of the *oriC/ter*. Data were plotted as whiskers plot. *P value < 0.05 (t-test), significantly up.

### Genetic interactions determination by CRISPRi-seq

Protocol for CRISPRi library construction, sequencing and analysis was performed as described before^49^. Briefly, 1,499 plasmids containing a different sgRNA were transformed into strain *P_tet_- dCas9, P_lac_-ccrZ, ΔccrZ* in presence of 1 mM IPTG to ensure the expression of wild type *ccrZ*, resulting in a pooled library containing the inducible CRISPRi system under control of an anhydrotetracycline (aTc) -inducible promoter and combined with a depletion of *ccrZ* under control of an IPTG-inducible promoter. Colonies were harvested and stored at −80°C. To ensure sufficient induction of the library, cells were grown for 8 generations in triplicates. The pooled libraries were diluted 1:100 from stock in 10 mL of CY medium supplemented or not with aTc 50 ng.mL^−1^ and 1 mM IPTG and grown at 37°C. At OD_600nm_ = 0.4, cells were harvested and their gDNA isolated and prepared for MiniSeq (Illumina) sequencing with a custom sequencing protocol (www.veeninglab.com/crispri-seq). sgRNA counts were retrieved and analyzed using DESeq2 package in R to evaluate the fitness cost of each sgRNA as previously described^49^. The effect of interaction between aTc treatments, across *ccrZ* complementation and depletion was compared with a log_2_FC of 1 and an alpha of 0.05.

### Quantifications and statistical analysis

Data analysis was performed using R and Prism (Graphpad). When comparing wild type phenotypes with *ccrZ* depletion/complementation, a Wilcoxon rank sum test with Bonferroni adjustment was used as we did not assume a normal distribution, since some mutant cells can behave like wild type because of the variable time of depletion or possible leakiness of *P_lac_* or *P_Zn_*. When using whiskers plot, the lower and upper whiskers represent, respectively, the minimum and maximum values of the data; the median is represented as a solid line and the lower and upper quartiles respectively represent the 25^th^ and 75^th^ percentiles. When plotting the *oriC/ter* ratios in Fig. 4, the outliers are also depicted by gray dots.

Data shown are represented as mean of at least three replicates ± SEM if data came from one experiment with replicated measurement, and ± SD if data came from separate experiments.

### Data availability

The data that support the findings of this study are available from the corresponding author upon request. Genomes sequences data are available at NCBI Sequence Read Archive (SRA) under the following accession number <u>PRJNA564501</u>.

## Acknowledgements

We appreciate the assistance from the Electron Microscopy Facility (EMF) and the Protein Analysis Facility (PAF) at the University of Lausanne (UNIL) and thank them for their support. We thank Wiep Klaas Smits (LUMC) for the Split-luc sequences and Tanneke den Blaauwen (UVA) for the mTQ^ox^ sequence prior to publication, Arnau Domenech (UNIL) for construction of strain *hlpA-LgBit hlpA-SmBit* and Zhian Salehian (NMBU) for help with cloning. Work in the Kjos lab is supported by a FRIMEDBIO grant (project number 250976) and a JPIAMR grant (project number 296906) from the Research Council of Norway. Work in the Murray lab was supported by a Wellcome Trust Senior Research Fellowship (204985/Z/16/Z) and a grant from the Biotechnology and Biological Sciences Research Council (BB/P018432/1). Work in the Grossman lab was supported, in part, by the National Institute of General Medical Sciences of the National Institutes of Health under award number R37 GM041934 and R35 GM122538. Work in the Veening lab is supported by the Swiss National Science Foundation (SNSF) (project grant 31003A_172861), a JPIAMR grant (40AR40_185533) from SNSF, a Novartis Foundation grant (#17B064) and ERC consolidator grant 771534-PneumoCaTChER.

## Author contributions

C.G. and J.W.V. wrote the paper with input from all authors. C.G., S.S., M.E.A., Y.M.S., X.L., G.A.S., S.P., R.R., J.D. and M.K. performed the experiments. C.G., S.S., M.E.A., M.K., H.M., S.G., A.D.G. and J.W.V designed, analyzed and interpreted the data.

## Extended Data

**Extended Data Fig. 1.**
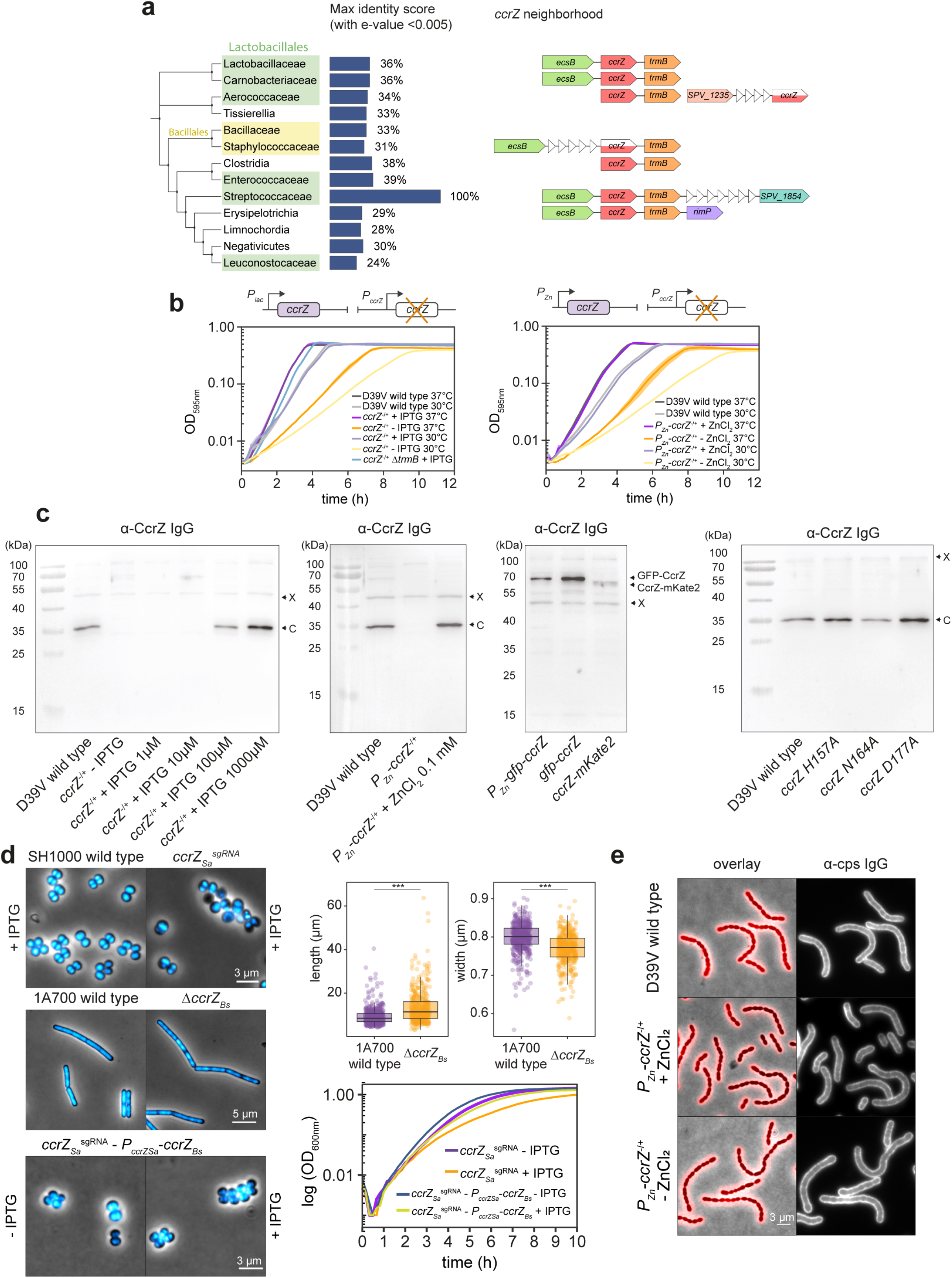
*ccrZ* deletion phenotype is conserved in *S. aureus.* **a**, Left: CcrZ conservation in firmicutes. Percentages indicate the highest percent identity, for each class, obtained using PSI-BLAST with NIH sequences. Right: genes co-occurrence in several genomes (data obtained from https://string-db.org; see Supplementary methods). Horizontal section indicate complexity in neighborhood score assignment; white triangles indicate missing annotation. **b**, Growth curves at 37°C and 30°C of *ccrZ* depletion mutants using *P_lac_* (left) or *P_Zn_* (right). **c**, Western blot from different pneumococcal strains. C: native CcrZ size; X: unknown protein recognized by α-ccrZ IgG. **d**, Microscopy of DAPI-stained *S. aureus* upon *ccrZ* silencing shows anucleate cells, while *B. subtilis* Δ*ccrZ* mutant did not present nucleoid defects. However, Δ*ccrZ* _Sa_ cells were longer (or less well separated) and thinner (top right; wild type: 483 cells, Δ*ccrZ_Bs_* 399 cells; each dot represents a measurement) ***: *p*-value <0.001 Wilcoxon rank sum test. Bottom-left: chromosome defects upon *ccrZ_Sa_* silencing can be rescued by expression of *ccrZ_Bs_* (*ccrZ_Sa_^sgRNA^*-*P_ccrZSa_*-*ccrZ_Bs_* + IPTG). Associated growth curves (bottom-right) also confirmed the complementation of *ccrZ_Sa_* by *ccrZ_Bs_*. **e**, Immunostaining of the polysaccharide capsule of *S. pneumoniae* wild type and upon *ccrZ* depletion.

**Extended Data Fig. 2.**
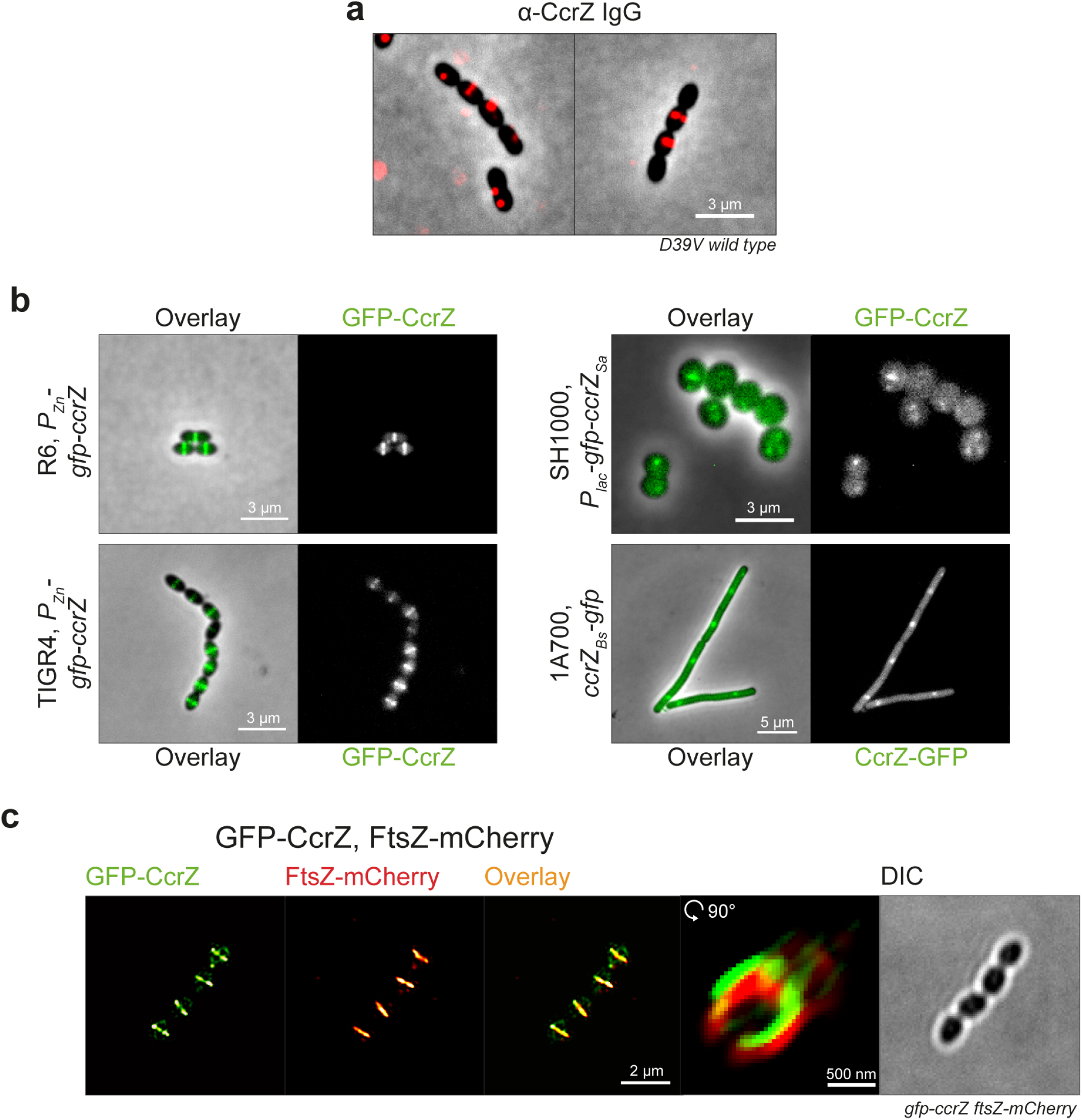
Septal localization of CcrZ in *S. pneumoniae*. **a**, Immunostaining of CcrZ in wild type *S. pneumoniae* shows a septal localization. **b**, Localization of CcrZ in other pneumococcal strains (un-encapsulated R6 strain and capsular serotype 4 TIGR4) and in *S. aureus* SH1000, as well as in *B. subtilis* 1A700. **c**, 3D-SIM of GFP-CcrZ (green) and FtsZ-mCherry (red) and reconstructed volume projection of both.

**Extended Data Fig. 3.**
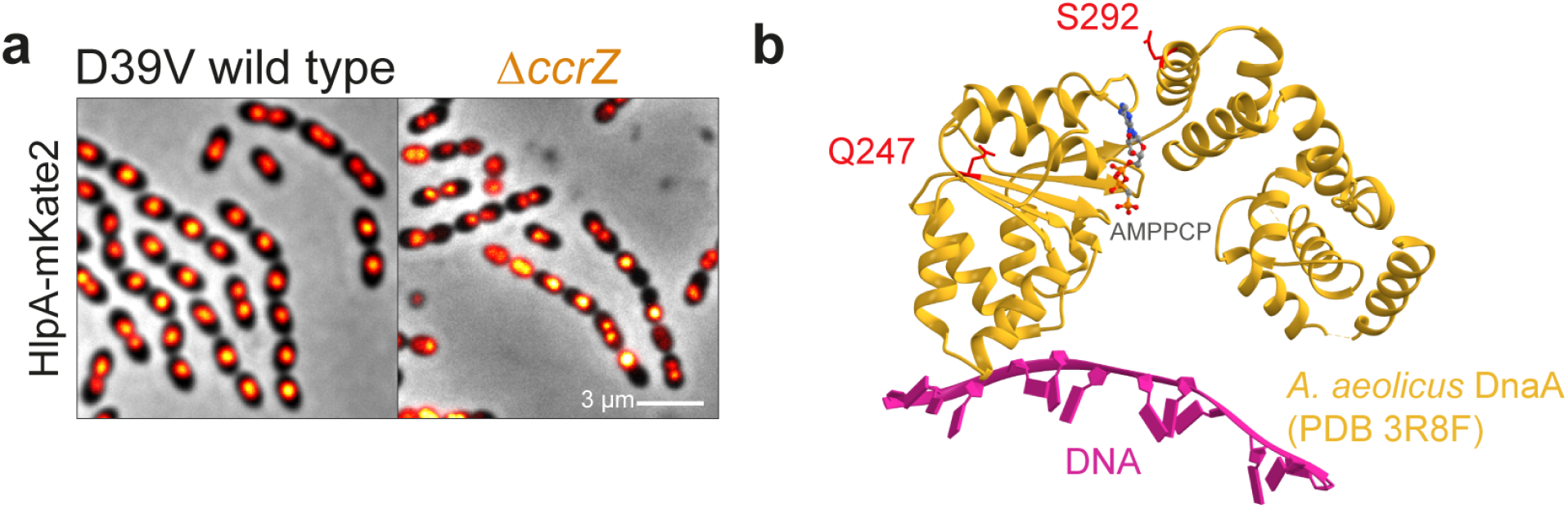
Deletion of *ccrZ* leads to anucleate cells. **a**, Localization of the nucleoid associated protein HlpA in a *ccrZ* mutant shows anucleate cells. **b**, Mapping of DnaA Q247 and S292 residues onto the crystal structure of DnaA’s AAA+ and duplex-DNA-binding domains from *Aquifex aeolicus*. Both residues are predicted to be in the AAA+ domain. DnaA Q247 and S292 correspond to *A. aeolicus* DnaA Q208 and E252 respectively.

**Extended Data Fig. 4.**
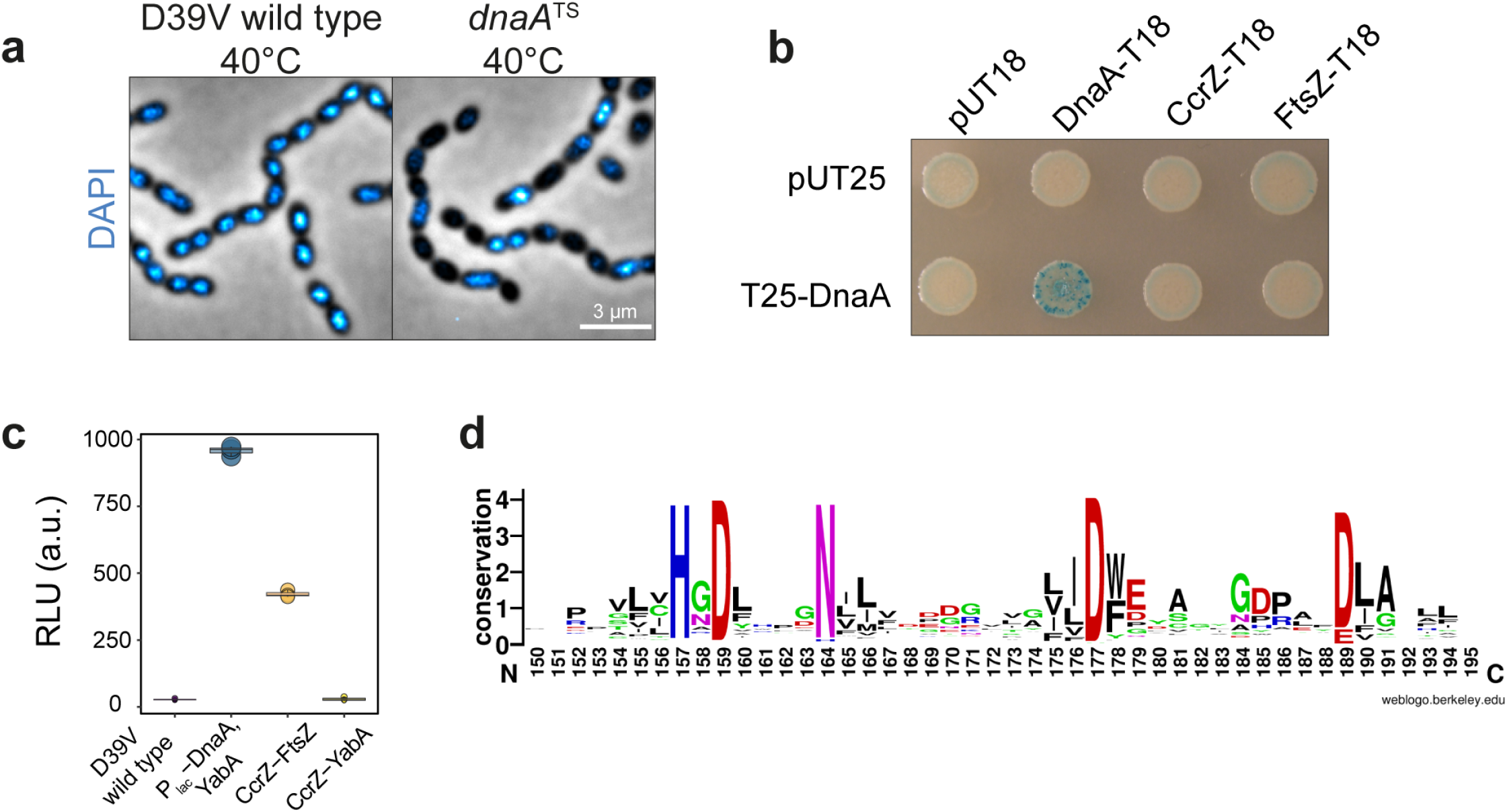
CcrZ activity is crucial for proper replication initiation as a *dnaA^TS^* mutant phenocopied a *ccrZ* deletion. **a**, Microscopy of DAPI-stained DnaA thermosensitive strain at non-permissive temperature (40°C) indicates several anucleate cells, compared to a wild type grown in identical conditions. **b**, No interaction was detected between DnaA and CcrZ using bacterial-2-hybrid, while a positive DnaA-DnaA self-interaction is visible. **c**, Using split-luc assay, no interaction between CcrZ-YabA was detected, while a strong signal was obtained for DnaA-YabA. DnaA level was controlled by P_lac_ to avoid toxicity. Each circle represents the average of 15 measurements of a technical replicate, with the size of the dot representing the SEM. **d**, Five (H157, D159, N164, D177 and D189) most conserved residues between 1000 different CcrZ sequences from different bacterial species; sequences obtained from UniRef50 database.

**Extended Data Fig. 5.**
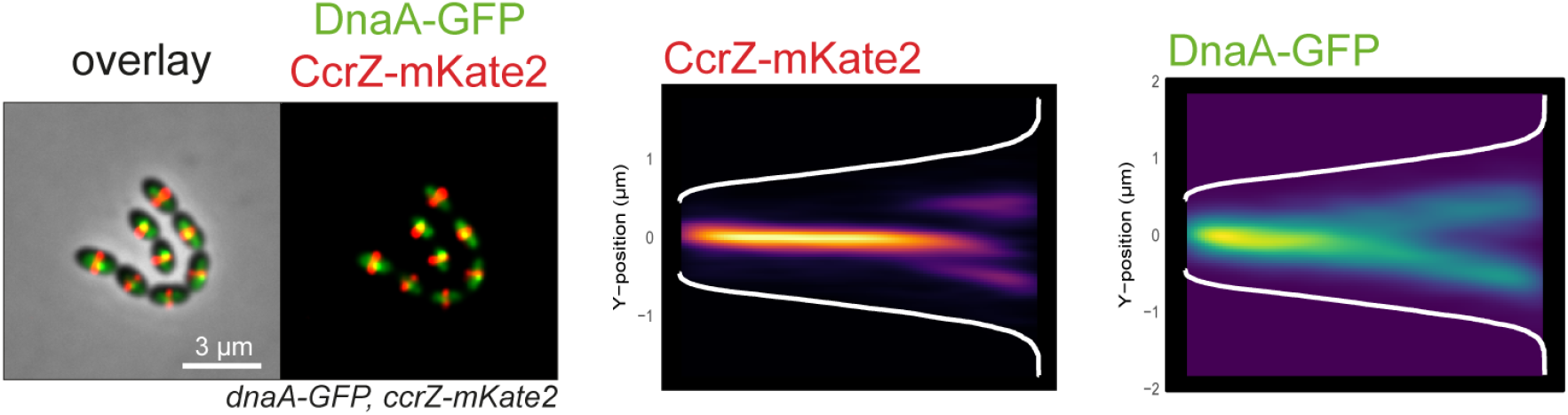
Transient co-localization of CcrZ and DnaA. Co-localization of CcrZ-mKate2 with DnaA-GFP (left) and corresponding heatmap of signal distribution over cell length (right) show that DnaA and CcrZ co-localize at the beginning of the cell cycle.

## Supplementary information

Supplementary Videos legends, Supplementary Methods, Supplementary Tables 1-3, and Supplementary References.

**Supplementary Table 4**

CRISPRi-seq results for ccrZ-complementation vs ccrZ-depletion

