## Supplemental methods and Supplemental figures for "Spatio-temporal control of DNA replication by the pneumococcal cell cycle regulator CcrZ"

**Contents**:

**Supplementary Videos legends**
**1**. 3D-SIM of GFP-CcrZ in wild type cells.
**2**. Time lapse microscopy of GFP-CcrZ in live cells.
**3**. Time lapse analysis of GFP-CcrZ and FtsZ-mCherry in live cells.
**4**. 3D-SIM of GFP-CcrZ and FtsZ-mCherry in wild type cells.
**5**. Time lapse microscopy of CcrZ-mKate2 in FtsZ depleted cells.
**6**. Time lapse analysis of HlpA-mKate2 in Δ*ccrZ* cells.
**7**. Time lapse microscopy of FtsZ-mTurquoise2 in DnaATS following temperature shift.
**8**. Time lapse analysis of CcrZ-mKate2, DnaN-sfTQOX and ParBp-sfmYFP in live cells.

**Supplementary Methods**

**Supplementary Tables 1-3**
**1**. Strains and plasmids used in this study
**2**. Oligonucleotides used in this study
**3**. Ratio of spectral counts between GFP-CcrZ and GFP from LC-MS/MS

**Supplementary References**

**Supplementary Video 1: 3D-SIM of GFP-CcrZ in wild type cells.**
Volume projection of 240 reconstructed 3D-SIM images from a chain of four live *S. pneumoniae* cells expressing msfGFP-CcrZ shows CcrZ forming patchy rings.

**Supplementary Video 2: Time lapse microscopy of GFP-CcrZ in live cells.**
Localization overtime of msfGFP-CcrZ (green) at 30°C overlaid with phase contrast (gray) shows that CcrZ localizes exclusively at the division site overtime. Time interval: 10 min.

**Supplementary Video 3: Time lapse analysis of GFP-CcrZ and FtsZ-mCherry in live cells.**
Localization of msfGFP-CcrZ (green) and FtsZ-mCherry (red) overlaid with phase contrast (gray) at 30°C shows that CcrZ and FtsZ co-localize overtime. Time interval: 10 min.

**Supplementary Video 4: 3D-SIM of GFP-CcrZ and FtsZ-mCherry in wild type cells.**
Volume projection of 240 reconstructed 3D-SIM images of a chain of four *S. pneumoniae* cells shows that msfGFP-CcrZ (green) and FtsZ-mCherry (red) form a similar ring structure and co-localize.

**Supplementary Video 5: Time lapse microscopy of CcrZ-mKate2 in FtsZ depleted cells.**
Depletion of FtsZ overtime shows a rapid spread of CcrZ-mKate2 (red) signal in the cytoplasm. Cells are visualized by phase contrast (grey). Time interval: 10 min.

**Supplementary Video 6: Time lapse analysis of HlpA-mKate2 in Δ*ccrZ* cells.**
Localization overtime of HlpA-mKate2 (red) used as a chromosomal marker in live *ccrZ*-deleted cells shows absence of nucleus and "guillotined" chromosome in several cells visualized by phase contrast (grey). Time interval: 5 min.

**Supplementary Video 7: Time lapse microscopy of FtsZ-mTurquoise2 in DnaA$^{TS}$ following temperature shift.**
Localization overtime of FtsZ-mTurquoise2 (cyan) in DnaA$^{TS}$ when cells are grown at 40°C following a pre-incubation at 30°C shows a delocalization of FtsZ after roughly four to five generations. Cells are visualized by phase contrast (grey). Time interval: 15 min.

**Supplementary Video 8: Time lapse analysis of CcrZ-mKate2, DnaN-sfTQ$^{OX}$ and ParB$_p$-sfmYFP in live cells.**
Localization overtime of CcrZ-mKate2 (red, marked by red arrows), DnaN-sfTQ$^{OX}$ (cyan, marked by cyan arrows) and ParB$_p$-sfmYFP (yellow, marked by yellow arrows) overlaid with phase contrast (grey) shows that the origin of replication is first brought to the future division site, while the replication machinery localizes at mid-cell with CcrZ, until CcrZ re-localizes to the new division site to start a new round of replication. Time interval: 2 min 30 sec.

### Supplementary Methods

#### Strains constructions

All strains and plasmids used for constructions are listed in Supplementary Table 1 and all primers are listed in Supplementary Table 2.

All *S. pneumoniae* strains were constructed by integrating into the chromosome by double homologous recombination either circular plasmids or linear DNA fragments possessing two ~1 kb region homology assembled using standard restriction-ligation, PCR assembly, Gibson assembly or Golden Gate assembly. In all cases parental strains were transformed directly with the assembled products. Constructs were confirmed by PCR and the resulting fragment sequenced.

$P_{Zn}$-$ccrZ^{+/+}$ and $ccrZ^{+/+}$. For insertion of a second copy of *spv_0476* (*ccrZ*) under control of either $P_{Zn}$ or *Plac*, *spv_0476* with its native RBS was amplified by PCR from D39V genome with primers 1/2 or 1/9 respectively and cloned into pMK11 vector[1] between EcoRI-SpeI restriction sites allowing for the genetic fusion $P_{Zn}$-*spv_0476* to be inserted in place of *bgaA* (coding for β-galactosidase) locus, or cloned into pPEPY[2] between EcoRI-BamHI sites to obtain the genetic fusion *Plac-spv_0476* for insertion in place of the *cil* locus (chromosomal integration locus – disrupting the non-coding *spv_2165* gene). pMK11-*spv_0476* was then transformed into D39V wild type and pPEPY-*spv_0476* into D39V *lacI*[3] as this strains expresses constitutively the repressor LacI.

$P_{Zn}$-$ccrZ^{-/+}$ and $ccrZ^{-/+.}$ For deletions of *spv_0476,* both flanking regions of the gene were amplified by PCR on D39V genome with primers 3/4 and 5/6 and assembled using Gibson assembly[4] to an erythromycin resistant marker amplified from a strain *ftsZ-mKate2*-ery[1] with primers 7/8. The final product was transformed into $P_{Zn}$-$ccrZ^{+/+}$ strain in presence of $ZnCl_2$ in order to express the extra copy of *ccrZ*. The resulting *ccrZ*::*ery* product together with the flanking regions was amplified from the $P_{Zn}$-$ccrZ^{-/+}$ genome with primers 10/11 and transformed into $ccrZ^{+/+}$ in presence of IPTG.

*ftsZ-spc* and *ftsZ-mTurquoise2*. The spectinomycin resistance marker s*pc* was amplified by PCR from was plasmid pPEP23[5] with primers 16/17, *ftsZ* and flanking regions were amplified on D39V genome with primers 12/13 and 14/15. All three fragments were assembled using Golden Gate assembly with BsmBI[6] and the resulting product transformed into D39V wild type. *mTurquoise2* was amplified from vector mTurquoise2-pBAD (addgene) with primers 19/20, *ftsZ-linker* was amplified from *ftsZ-mKate2* strain[1] with primers 12/18 and downstream flanking region of *ftsZ* with *spc* amplified on *ftsZ-spc* genome with primers 15/21. All three fragments were assembled using Golgen Gate assembly (BsmBI) and transformed into D39V wild type in order to replace the native *ftsZ* with *ftsZ-mTurquoise2-ery* fusion.

$ccrZ^{-/+}$ *ftsZ-mTurquoise2* and $dnaA^{TS}$ *mTurquoise2*. *ftsZ-mTurquoise2* genetic fusion and flanking regions were amplified by PCR with primers 12/15 from *ftsZ-mTurquoise2* strain and inserted either into $ccrZ^{-/+}$ strain in presence of IPTG to keep high levels of CcrZ ($ccrZ^{-/+}$ *ftsZ-mTurquoise2*) or into the thermosensitive *dnaA* mutant $dnaA^{TS}$[7] ($dnaA^{TS}$ *mTurquoise2*).

$ccrZ^{-/+}$ Δ*trmB*. For co-deletion of *ccrZ* and *trmB* (same operon and overlapping), upstream region of *ccrZ* was amplified by PCR on D39V genome with primers 22/23, erythromycin resistance marker was amplified with primers 24/25 on *ccrZ*$^{-/+}$ genome and downstream region of *trmB* was amplified on D39V genome with primers 26/27. All three fragments were assembled using Golden Gate (BsaI) and inserted into strain *ccrZ*$^{-/+}$ in presence of IPTG.

*P*$_{Zn}$*-ccrZ-gfp*, *P*$_{Zn}$*-ftsZ-gfp,* *P*$_{Zn}$*-gfp-ccrZ*, R6 *P*$_{Zn}$*-gfp-ccrZ* and TIGR4 *P*$_{Zn}$*-gfp-ccrZ*. For translational fusion of monomeric superfolder green fluorescent protein (msfGFP) at the C-terminal extremities of CcrZ and FtsZ, *ccrZ* and *ftsZ* together with their respective RBS were amplified by PCR on D39V genome with primers 28/29 and 30/31, respectively, and cloned into plasmid pMK17[1] between restriction sites NotI-SpeI, allowing for insertion of fusion *ccrZ-gfp* or *ftsZ-gfp* under control of $P_{Zn}$ at the *bgaA* locus. The final product was then transformed into D39V wild type, R6[8] and TIGR4[9]. For translational fusion of msfGFP at the N-terminal extremity of CcrZ (as *ccrZ* 3' region overlaps with *trmB* 5' region), *ccrZ* excluding the START codon and including STOP codon was first amplified by PCR from D39V genome with primers 32/33 and cloned into pCG6[3] between restriction sites SpeI-NotI, allowing for genetic fusion of *msfgfp* to the 5' of *ccrZ* under control of $P_{Zn}$. pCG6 derivates from pMK17 and can therefore integrates into the *bgaA* locus. D39V wild type was then transformed with the final product.

*gfp-ccrZ* and *gfp-ccrZ ftsZ-mCherry*. *gfp-ccrZ* fragment was amplified by PCR from *P*$_{Zn}$*-gfp-ccrZ* strain with primers 37/38, upstream flanking region was amplified with primers 34/35 from D39V genome and a kanamycin marker was amplified with primers 36/39 from pPEP2K1[10]. All three fragments were assembled by Gibson assembly and transformed into D39V wild type resulting in strain *gfp-ccrZ*, with *gfp-ccrZ* fusion in place of native *ccrZ* and under control of the native $P_{ccrZ}$ promoter. For co-localization of CcrZ with FtsZ, *ftsZ-mCherry* was amplified by PCR with primers 12/15 from *ftsZ-mCherry* strain [1] and transformed into *gfp-ccrZ* strain.

*ftsZ*$^{+/+}$ and *ftsZ*$^{-/+}$. For insertion of an inducible ectopic version of *ftsZ*, *ftsZ* gene was amplified from D39V genome by PCR with primers 40/41 and cloned into vector pPEPY between EcoRI and BamHI restriction sites. D39V *lacI* was then transformed with the ligated product, leading to insertion of *P*$_{lac}$*-ftsZ* in the *cil* locus. Spectinomycin marker together with *ftsZ* downstream region were amplified with primers 15/42 from strain *ftsZ-spc* and the region upstream of *ftsZ* START codon was amplified by PCR from D39V genome with primers 43/44. Both fragments were assembled using Golden Gate assembly (BsmBI) and the strain *ftsZ*$^{+/+}$ was transformed with the resulting product in presence of IPTG, to keep a high level of FtsZ, leading to the depletion strain *ftsZ*$^{-/+}$.

*ccrZ-mKate2* and *ftsZ*$^{-/+}$ *ccrZ-mKate2*. For genetic fusion of *ccrZ* with the fluorescent protein mKate2, *ccrZ* upstream and downstream regions where amplified by PCR with primers 45/46 and 49/50, respectively, from D39V genome and *mKate2-ery* (with linker) was amplified with primers 47/48 from *ftsZ-mKate2* strain. All three fragments were assembled by Golden Gate assembly (BsmBI) and the product transformed into D39V wild type. *ccrZ-mKate2* was then amplified by PCR from the previously created strain with primers 50/51 and the product transformed into *ftsZ*$^{-/+}$ strain in presence of IPTG to keep a high level of FtsZ.

*ccrZ-LgBit*. In order to fuse *ccrZ* to *LgBit* sequence in place of the native *ccrZ*, upstream region of *ccrZ* was amplified by PCR with primers 52/53 and the downstream region together with erythromycin marker was amplified with primers 54/55 on strain *ccrZ-mKate2*. A gBlocks fragment (Integrated DNA

Technologies) containing the *LgBit* sequence flanked with two BsaI sites was synthesised. Both PCR fragments and gBlocks were assembled using Golden Gate assembly (BsaI) and inserted into D39V wild type by transformation.

*ftsZ-SmBit* and *ccrZ-LgBit ftsZ-SmBit*. For fusion of *ftsZ* with *SmBit* sequence, *SmBit* sequence was synthesized as a gBlocks fragment (Integrated DNA Technologies) and amplified by PCR with primers 56/57. *ftsZ* with its upstream region sequence were amplified with primers 58/59 on D39V genome and the downstream region with spectinomycin marker were amplified with primers 60/61 on strain *ftsZ-spc*. All three fragments were then assembled with Golden Gate assembly (BsmBI) and transformed into D39V wild type and *ccrZ-LgBit* strains.

*ccrZ-SmBit* and *ccrZ-LgBit ccrZ-SmBit*. For *ccrZ-SmBit* construction, *ccrZ* upstream and downstream regions were amplified by PCR with primers 62/63 and 66/67, respectively, on D39V gDNA and SmBit was amplified with primers 64/65 from strain *ftsZ-SmBit*. All three fragments were assembled with Golden Gate assembly (BsmBI) and the product was transformed into strain D39V wild type. *ccrZ-SmBit* was then amplified by PCR with primers 70/67 from the resulting strain and *ccrZ-LgBit*, with upstream region was amplified with primers 68/69 from strain *ccrZ-LgBit*. The two fragments obtained were assembled by Golden Gate assembly (BsmBI) and transformed into D39V wild type.

*ccrZ-LgBit cps2E-SmBit, ccrZ-LgBit hlpA-SmBit, ccrZ-LgBit dnaA-SmBit, ccrZ-LgBit ezrA-SmBit, ccrZ-LgBit ftsA-SmBit, ccrZ-LgBit ftsH-SmBit, ccrZ-LgBit ftsW-SmBit, ccrZ-LgBit pepN-SmBit, ccrZ-LgBit pbp2x-SmBit, ccrZ-LgBit zapA-SmBit, ccrZ-LgBit fruA-SmBit, ccrZ-LgBit plsC-SmBit, ccrZ-LgBit scrA-SmBit* and *ccrZ-LgBit spv_1621-SmBit*. All fourteen double LgBit-SmBit labeled strains were constructed as followed. *SmBit* fragment was amplified by PCR from *ftsZ-SmBit* genome with primers 71/72 when using BsmbI and 73/74 when using BsaI (for *dnaA-SmBit and ezrA-SmBit*). *cps2E, hlpA*, *dnaA, ezrA, ftsA, ftsH, ftsW, pepN, pbp2x, zapA, fruA, plsC, scrA* and *spv_1621* upstream (of Start codon) regions were amplified from D39V genome with primers 75/76, 79/80, 83/84, 87/88, 91/92, 95/96, 99/100, 103/104, 107/108, 111/112, 115/116, 119/120, 123/124 and 127/128, respectively, and downstream regions were amplified from D39V genome with primers 77/78, 81/82, 85/86, 89/90, 93/94, 97/98, 101/102, 105/106, 109/110, 113/114, 117/118, 121/122, 125/126 and 129/130, respectively. Each downstream and upstream fragment was assembled to *SmBit* fragment using Golden Gate assembly (BsmBI for all except *dnaA* and *ezrA* using BsaI) and transformed into strain *ccrZ-LgBit*.

*hlpA-LgBit* and *hlpA-LgBit hlpA-SmBit*. For construction of the final strain *hlpA-LgBit hlpA-SmBit* used as positive control for SplitLuc assay as HlpA proteins interact together [11], a first strain *hlpA-LgBit* was made as followed. *Prs1* upstream region was amplified by PCR with primers 131/132 on strain VL2429 (Veening lab collection) and fragment *LgBit* fuse to chloramphenicol resistant marker, together with *prs1* downstream region was amplified with primers 135/136 from strain VL2429. *hlpA* was amplified with primers 133/134 from D39V genome. All three fragments were assembled by overlapping PCR using primers 131/136 and transformed into D39V wild type. *hlpA-SmBit* was amplified from strain *ccrZ-LgBit hlpA-SmBit* with primers 79/82 and transformed into the previously created strain *hlpA-LgBit*, leading to strain *hlpA-LgBit hlpA-SmBit*.

*$P_{lac}$-dnaA-SmBit*. For insertion of *dnaA* fused to *SmBit* under control of $P_{lac}$ promoter, plasmid pPEPZ[2] was amplified by PCR with primers 222/223 (upstream) and 224/225 (downstream) and *dnaA-SmBit* was amplified by PCR with primers 226/227 from strain *ccrZ-LgBit dnaA-SmBit*. All three fragments were assembled using Golden Gate (BsmBI) and inserted into D39V *lacI* strain, allowing for insertion of *$P_{lac}$-dnaA-SmBit* into the *zip* locus (pPEPZ integration platform - causing disruption of the non-coding gene *spv_2417*).

*$P_{lac}$-dnaA-SmBit yabA-LgBit* and *yabA-LgBit*. For fusion of *yabA* with *LgBit*, *LgBit* fragment was amplified by PCR with primers 228/229 from strain *ccrZ-LgBit* and *yabA* together with upstream and downstream regions were amplified with primers 230/231 and 232/233 respectively. All three fragments were assembled using Golden Gate (BsmBI) and inserted into strain *$P_{lac}$-dnaA-SmBit* and D39V, leading to strains *$P_{lac}$-dnaA-SmBit yabA-LgBit* and *yabA-LgBit*.

*yabA-LgBit ccrZ-SmBit*. *ccrZ-SmBit* fragment, containing upstream and downstream regions, was amplified from *ccrZ-SmBit* strain using primers 62/67 and transformed into strain *yabA-LgBit*, resulting in strain *yabA-LgBit ccrZ-SmBit*.

*hlpA-mKate2 ΔccrZ*. *ccrZ ::ery* fragment was amplified by PCR with primers 10/11 on strain *$ccrZ^{-/+}$* and transformed into *hlpA-mKate2* strain[1].

*$P_{Zn}$-$ccrZ^{+/+}$ ftsZ-cfp, $P_{Zn}$-$ccrZ^{+/+}$ ftsZ-cfp hlpA-mKate2* and *$P_{Zn}$-$ccrZ^{-/+}$ ftsZ-cfp hlpA-mKate2*. *ftsZ-cfp* was amplified from *ftsZ-cfp* strain[1] by PCR with primers 12/15 and transformed into strain *$P_{Zn}$-$ccrZ^{+/+}$* leading to strain *$P_{Zn}$-$ccrZ^{+/+}$ ftsZ-cfp*. *hlpA-mKate2* was amplified by PCR from strain *hlpA-mKate2*[1] with primers 137/138 and the resulting fragment was used for transformation of strain *$P_{Zn}$-$ccrZ^{+/+}$ ftsZ-cfp*. The newly created strain *$P_{Zn}$-$ccrZ^{+/+}$ ftsZ-cfp hlpA-mKate2* was then transformed in presence of IPTG with *ccrZ::ery* fragment amplified by PCR with primers 10/11 on strain *$ccrZ^{-/+}$*, resulting in strain *$P_{Zn}$-$ccrZ^{-/+}$ ftsZ-cfp hlpA-mKate2*.

*$ccrZ^{supp1}$, $ccrZ^{supp2}$* and *$ccrZ^{supp3}$*. The deletion fragment *ccrZ::ery* was amplified by PCR with primers 10/11 on strain *$ccrZ^{-/+}$* and inserted in place of *ccrZ* in strain D39V wild type. Among the 10,000 colonies appearing on erythromycin-agar plates, 200 colonies were as large as wild type colony, while nearly 9,800 were very small colonies. 3 large colonies were cultivated, and their growth was assessed by plate-reader assay, before to be stored at -80°C. Mutation determination is described in *Genome resequencing of ccrZ suppressors by NGS* section. *$ccrZ^{supp1}$* has a 741G>T substitution leading to DnaA-Q247H and an additional insertion 13_14insA into *licD2*; *$ccrZ^{supp2}$* has a 874G>T substitution leading to DnaA-S292G; and *$ccrZ^{supp3}$* had a 277G>T substitution leading to YabA E93* (insertion of STOP codon).

*ΔccrZ dnaA-Q247H* and *ΔccrZ dnaA-S292G*. For re-insertion of the two *dnaA* point mutations in wild type background, fragments corresponding to *dnaA-Q247H* and *dnaA-S292G* with flanking region (marker-less) were first constructed by PCR assembly. The fragment corresponding to *dnaA-Q247H*_up was amplified on D39V genome with primers 143/140 and *dnaA-Q247H*_down was amplified with primers 139/144; a final PCR with primers 143/144 using both fragments as template led to the final *dnaA-Q247H* product. *dnaA-S292G* fragment was constructed in the same way using primers 143/142 and 141/144. Either assemblies were then inserted into D39V wild type, together with the deletion fragment *ccrZ::ery* (amplified by PCR with primers 10/11 on strain *ccrZ*$^{-/+}$), allowing for selection. In both cases, small colonies with few large colonies (2%) were present and only the large colonies were selected for confirmation by PCR and sequencing.

*kan-ccrZ, dnaA-Q247H* and *dnaA-S292G*. In order to re-insert *ccrZ* into strain Δ*ccrZ dnaA-Q247H* and Δ*ccrZ dnaA-S292G*, a strain *kan-ccrZ* was constructed as intermediate. The upstream region of *ccrZ*, together with *kan* (kanamycin resistant marker), was amplified by PCR on strain *gfp-ccrZ* with primers 45/145 and *ccrZ* gene was amplified with primers 146/11 on D39V genome. Both fragments were then assembled using Golden Gate assembly (BsmBI) and inserted into D39V wild type. *kan-ccrZ* fragment was amplified by PCR from the resulting strain *kan-ccrZ* with primers 22/55. Strains Δ*dnaA Q247H* and Δ*dnaA S292G* were then transformed with the obtained DNA fragments and selected for kanamycin resistance, allowing for replacement of *ccrZ*::*ery* with *kan-ccrZ*.

Δ*yabA* and Δ*yabA* Δ*ccrZ*. For *yabA* deletion, sequences upstream and downstream of *yabA* were amplified on D39V genome by PCR with primers 147/148 and 151/152, respectively, and spectinomycin resistance marker was amplified on pPEP23 with primers 149/1509. All fragments were then assembled using Golden Gate assembly (BsmBI) and inserted into D39V wild type. Δ*yabA* strain was then transformed with a fragment *ccrZ*::*ery* amplified by PCR with primers 10/11 on strain *ccrZ*$^{-/+}$ resulting in strain Δ*yabA* Δ*ccrZ*.

*ccrZ-N164A*, *ccrZ-H157A* and *ccrZ-D177A*. Insertion of *ccrZ* point mutations into $P_{Zn}$-*ccrZ*$^{+/+}$ strain was performed as followed. For *ccrZ-N164A*, *kan-ccrZ_up* was amplified on *kan-ccrZ* genome with primers 45/154 and *ccrZ_down* was amplified with primers 153/11 on D39V genome. Both fragments were assembled by PCR assembly using primers 45/11. *ccrZ-H157A* and *ccrZ-D177A* were constructed in a similar manner, with primers 45/156, 155/11, 45/158 and 157/11 respectively. All three assembled products were then inserted into strain $P_{Zn}$-*ccrZ*$^{+/+}$ in presence of $ZnCl_2$ in order to keep high CcrZ levels.

$P_{lac}$-*ccrZ-gfp*, $P_{lac}$-*ccrZ-H157A-gfp*, $P_{lac}$-*ccrZ-N164A-gfp* and $P_{lac}$-*ccrZ-D177A-gfp*. For insertion of *ccrZ* fused to *gfp* under control of $P_{lac}$ promoter, *ccrZ-gfp* was amplified from $P_{Zn}$-*ccrZ-gfp* strain genome with primers 161/162 and plasmid pPEPZ[2] was amplified by PCR with primers 159/160. Assembly of both fragments using Golden Gate (BsmBI) and insertion into D39V *lacI* strain allowed insertion of $P_{lac}$-*ccrZ-GFP* into the *zip* locus (pPEPZ integration platform - causing disruption of the non-coding gene *spv_2417*). To mutate *ccrZ-gfp*, PCR assembly was used for the three mutants. For $P_{lac}$-*ccrZ-H157A-gfp*, upstream of the *zip* locus together with $P_{lac}$-*ccrZ_up* were amplified by PCR with primers 163/156 and *ccrZ-gfp_down* with downstream region of *zip* locus was amplified with primers 155/164, from $P_{lac}$-*ccrZ-gfp* genome. Both fragments were assembled by PCR assembly using primers 163/164. $P_{lac}$-*ccrZ-N164A-gfp* and $P_{lac}$-*ccrZ-D177A-gfp* fragments were constructed in a similar manner, using primers 163/154, 153/164, 163/158 and 157/164 respectively. All three products were then inserted into strain D39V *lacI*.

*comCDE-parS*$_p$ *ccrZ-mKate2*, *parB*$_p$*-YFP comCDE-parS*$_p$ *ccrZ-mKate2* and *dnaN-mTQ*$^{ox}$ *parB*$_p$*-YFP comCDE-parS*$_p$ *ccrZ-mKate2*. In order to visualize the replication machinery together with the origin of replication and CcrZ, a first strain was made, expressing CcrZ-mKate2 and containing $parS_p$ sites from *Lactococcus lactis* (able to bind $parB_p$ proteins from *L. lactis*) in the genome close to the origin of replication *oriC*. *comCDE-parS*$_p$ was amplified by PCR from strain D39V, *comCDE-parS*$_p$ *bgaA*::*parB*$_p$*-sfmGFP*[1] using primers 165/166 and the resulting fragment was transformed into strain *ccrZ-mKate2*. Plasmid pMK19-02 (carrying *bgaA*::$P_{Zn}$-*parB*$_p$-*msfYFP*, Veening Lab collection ) was transformed into the resulting strain *comCDE-parS*$_p$ *ccrZ-mKate2*. To prevent any alteration in replication process, the genetic fusion of *dnaN* with *sfmTurquoise2*$^{ox}$ was introduced as a second copy downstream of the original *dnaN* gene. *dnaN* and upstream region were amplified by PCR with primers 167/168 from D39V genome, the second *dnaN* copy was amplified with primers 169/170 from D39V genome, *sfmTurquoise2*$^{ox}$ with linker were amplified with primers 173/174 from strain D39V, *CEP*::*P3-spv_1159-sfmTurquoise2*$^{ox}$*-opt* (Veening lab collection ), chloramphenicol resistance marker (*cam*) was amplified with primers 175/176 from *hlpA-mKate2* strain and downstream of *dnaN* was amplified with primers 171/172 from D39V genome. All five fragments were assembled using Golden Gate assembly (BsaI for the three first fragments and BsaI / SapI for the two others) and transformed into strain *parB*$_p$*-YFP comCDE-parS*$_p$ *ccrZ-mKate2*.

*ccrZ-mKate2* $P_{Zn}$*-dnaA-GFP*. *dnaA* was amplified from D39V genome by PCR with primers 177/178 and the resulting fragment was digested with NotI-SpeI restriction enzymes and ligated into plasmid pMK17, allowing genetic fusion with *msfgfp* under control of $ZnCl_2$-inducible promoter. The resulting product was transformed into strain *ccrZ-mKate2*.

*ccrZ*$_{Sa}$$^{sgRNA}$ and *ccrZ*$_{Sa}$$^{sgRNA}$ $P_{ccrZSa}$*-ccrZ*$_{Bs}$. Inverse PCR was used to construct sgRNA-plasmid, as described previously[3,12]; in which the phosphorylated primer 179 was combined with gene specific forward primers containing the 20 bp targeting region as overhangs. Primer 180 was used as specific primer to construct the plasmid pCG248-sgRNA(*ccrZ*$_{Sa}$). The sgRNA was designed to target the 5' end *ccrZ*$_{Sa}$. For construction of the complementation plasmid pCG248-sgRNA(*ccrZ*$_{Sa}$)-$P_{ccrZ}$*-ccrZ*$_{Bs}$, *B. subtilis ccrZ*-homolog, *ytmP* (*ccrZ*$_{Bs}$), was fused to the *ccrZ*$_{Sa}$ promoter and integrated into plasmid pCG248-sgRNA(*ccrZ*$_{Sa}$). *ccrZ*$_{Bs}$ was amplified from plasmid pSG3174 (pUC19-*ccrZ*$_{Bs}$, see below) using primers 183/184. *ccrZ*$_{Sa}$ -promoter ($P_{ccrZSa}$) was amplified from *S. aureus* SH1000 genome using primers 181/182. Both fragments were fused in a second PCR step, using primers 181/184. The resulting fragment was digested with restriction enzymes BamHI and KpnI and ligated into plasmid pCG248-sgRNA(*ccrZ*$_{Sa}$).

$P_{lac}$*-gfp-ccrZ*$_{Sa}$. For expression of CcrZ-GFP under control of an IPTG-inducible promoter in *S. aureus* SH1000, *ccrZ*$_{Sa}$ gene was amplified from genomic DNA of *S. aureus* SH1000 using primers 185/186. The fragment was digested with NcoI and BamHI and ligated into the corresponding sites of plasmid pLOW-*parB-msfgfp*. The resulting plasmid expressing the translational fusion CcrZ$_{Sa}$-msfGFP fusion under control of an IPTG-inducible promoter was then transformed into SH1000 strain.

$P_{tet}$*-dCas9,* $P_{lac}$*-ccrZ,* Δ*ccrZ*. A *S. pneumoniae* strain expressing lacI and tetR under control of the constitutive promoter PF6 (*prs1*::PF6-*lacI-tetR*) was transformed with a DNA fragment $P_{tet}$*-dCas9,* amplified by PCR from strain VL2212[13] (D39V, *prs1*::*PF6-tetR*, *bgaA::*$P_{tet}$*-dcas9*) with primers 219/220, that integrated in the *bgaA* region. The resulting $P_{tet}$*-dCas9* strain was then transformed with a PCR fragment $P_{lac}$*-ccrZ* amplified with primers 217/218 from strain *ccrZ*$^{+/+}$. This fragment integrated in place of the *CIL* locus. The resulting strain $P_{tet}$*-dCas9,* $P_{lac}$*-ccrZ* was then transformed in presence of 0.1 mM IPTG with a PCR fragment *ccrZ::ery* amplified with primers 10/11 from strain *ccrZ*$^{-/+}$, resulting in strain $P_{tet}$*-dCas9,* $P_{lac}$*-ccrZ,* Δ*ccrZ.*

All *B. subtilis* strains were constructed by integrating into the chromosome by double homologous recombination using either genomic DNA or linear DNA fragments possessing two ~1 kb region homology. Assembled products were introduced into parent strains using natural transformation. Constructs were confirmed by PCR and the resulting fragment sequenced.

*1A700 ΔccrZ$_{Bs}$*. In frame deletion of *ccrZ* homolog in *B. subtilis*, *ytmP* (*ccrZ$_{Bs}$*), was performed using Golden Gate allelic replacement strategy as described before[14]. Upstream homology region of *ccrZ$_{Bs}$* was amplified by PCR with primers 187/188 and ligated into plasmid pUC19 (leading to pSG3174), 5' region and 3' region (including downstream homology of *ccrZ$_{Bs}$*) were amplified with primers pairs 189/190 and 191/192 and ligated into plasmid pUC19 (leading to pSG3178 and pSG3177, respectively). The three resulting plasmids and plasmid pSG0682[14] (carrying an erythromycin resistance cassette) where assembled together with Golden Gate backbone plasmid pSG1525[14] and transformed into 1A700 wild type.

*ccrZ$_{Bs}$-gfp*. For translational fusion of *ccrZ$_{Bs}$* with msfGFP, *ccrZ$_{Bs}$* (*ytmP*) was amplified by PCR with primers 193/194 and ligated into plasmid pUC19 (leading to pSG3175), *msfgfp* was amplified with primers 195/196 and ligated into plasmid pUC19 (giving pSG3179) and homology region downstream of *ccrZ$_{Bs}$* was amplified with primers pair 189/190 and ligated into pUC19 (leading to pSG3176). The three plasmids obtained and plasmid pSG0682 were assembled using Golden Gate and transformed into 1A700 wild type strain.

*ΔccrZ$_{Bs}$*. Deletion of *ccrZ$_{Bs}$* was constructed by replacing the open reading frame with a chloramphenicol resistance cassette (cam) by using linear Gibson assembly fragments containing ~1KB of flanking homology for *ccrZ$_{Bs}$*. Upstream of *ytmP* (*ccrZ$_{Bs}$*) was amplified using primers 197/198; the chloramphenicol resistance cassette was amplified from pGEM*::cat*[15] using primers 199/200; downstream of *ccrZ$_{Bs}$* was amplified using primers 201/202.

*oriN* and *oriN ΔccrZ$_{Bs}$*. Strain *oriN* was constructed by introducing the heterologous origin and initiator (*oriN/repN*)[16] in place of *oriC* and introducing a constitutive promoter to drive expression of *dnaN*. The constitutive promoter used was Ppen[17] with the following sequence replacing the -10 to -35 box (5'-GTTGCATTTATTCTTAGATAGTGTAATACT-3'). The various fragments were amplified by PCR and assembled using Gibson assembly. The fragments were amplified using the following primers and templates: upstream *dnaA* was amplified using primer 203/204, kanamycin cassette was amplified from pGK67[18] using primers 205/206, *oriN*/*repN* was amplified from pDL110[16] using primers 207/208, Ppen2027 was amplified from CAL2072 using 209/210 and *dnaN* was amplified with primers 211/212. The assembled fragments were then transformed into strain JH642 (*oriN*) or Δ*ccrZ$_{Bs}$* (*oriN* Δ*ccrZ$_{Bs}$*).

**Capsule immunofluorescence**

For fluorescence analysis of *S. pneumoniae* polysaccharide capsule, cells were grown in C+Y medium at 37°C until $OD_{595nm}$ = 0.1 and 1:1000 diluted serum anti-serotype 4 from rabbit (Neufeld antisera, Statens Serum Institut) was added for 5 min at 4°C. Cells were washed three times with fresh C+Y medium and 1 $mg.mL^{-1}$ of secondary antibody anti-rabbit coupled to Alexa Fluor 555 (Invitrogen) was added for 5 min at 4°C. Cells were then spotted onto a PBS-agarose slide. Acquisition of the fluorescent signal was performed on DV Elite microscope with mCherry filter set (Ex: 575/25 nm, BS: 605/50, Em: 632/60 nm).

**Conservation and gene neighborhood**

CcrZ protein sequence was aligned against all non-redundant protein sequences from NIH (ncbi.nlm.nih.gov) using PSI-BLAST for different firmicutes families. Sequences with highest identity were then aligned using Clustal Omega (ebi.ac.uk/Tools/msa/clustalo) and a phylogenetic three was generated by iTOL (Interactive Tree Of Life; itol.embl.de). Gene neighborhood data were obtained from the STRING database[19]. For residues conservation data, 1000 sequences of *ccrZ* homologs were retrieved with PSI-BLAST (ebi.ac.uk) from the UniRef50 database. Conservation visualization was obtained using WebLogo 3 (weblogo.threeplusone.com). Sequences were then aligned using Clustal Omega and CcrZ sequence with conservation scores was mapped using UCSF Chimera (cgl.ucsf.edu/chimera) onto the crystal structure of *S. pneumoniae* LicA (PDB 4R78), the closest homolog protein using HMM-HMM comparison with HHpred[20].

### Supplementary Table 1: Strains and plasmids used in this study

| ***S. pneumoniae* strains** | **Relevant genotype** | **Reference** |
|---|---|---|
| D39V wild type | *S. pneumoniae* serotype 2 | 21 |
| R6 wild type | *S. pneumoniae* D39 derivative | gift from C. Grangeasse laboratory, Lyon France |
| TIGR4 wild type | *S. pneumoniae* serotype 4 | 22 |
| $ccrZ^{sgRNA}$ | D39V, *prs1::lacI*, *bgaA::*$P_{lac}$*-dCas, CEP::P3-ccrZ-sgRNA*; $Gen^R$, $Tet^R$, $Spc^R$ | 3 |
| $P_{Zn}$*-ccrZ*$^{+/+}$ | D39V, *bgaA::*$P_{Zn}$*-spv_0476*; $Tet^R$ | This study |
| $P_{Zn}$*-ccrZ*$^{-/+}$ | D39V, *bgaA::*$P_{Zn}$*-spv_0476*, Δ*spd_0476*; $Tet^R$, $Ery^R$ | This study |
| D39V *lacI* | D39V, *prs1::lacI*; $Gen^R$ | 3 |
| *ccrZ*$^{+/+}$ | D39V, *prs1::lacI*, *cil::*$P_{lac}$*-spv_0476*; $Gen^R$, $Kan^R$ | This study |
| *ccrZ*$^{-/+}$ | D39V, *prs1::lacI*, *cil::*$P_{lac}$*-spv_0476*, Δ*spd_0476*; $Gen^R$, $Kan^R$, $Ery^R$ | This study |
| *ftsZ-spc* | D39V, *ftsZ-spc*; $Spc^R$ | This study |
| *ftsZ-mTurquoise2* | D39V, *ftsZ::ftsZ-mTurquoise2*; $Spc^R$ | This study |
| *ccrZ*$^{-/+}$ *ftsZ-mTurquoise2* | D39V, *prs1::lacI*, *cil::*$P_{lac}$*-spv_0476*, Δ*spd_0476*, *ftsZ::ftsZ-mTurquoise2*; $Gen^R$, $Kan^R$, $Ery^R$, $Spc^R$ | This study |
| *dnaA*$^{TS}$ | D39V, *dnaA::dnaA-M398T* | 7 |
| *dnaA*$^{TS}$ *ftsZ-mTurquoise2* | D39V, *dnaA* thermosensitive; *ftsZ::ftsZ-mTurquoise2; Spc*$^R$ | This study |
| *ccrZ*$^{-/+}$ Δ*trmB* | D39V, *prs1::lacI*, *cil::*$P_{lac}$*-spv_0476*, *spv_0476-trmB::ery*; $Gen^R$, $Kan^R$, $Ery^R$ | This study |
| $P_{Zn}$*-ccrZ-gfp* | D39V, *bgaA::*$P_{Zn}$*-spv_0476-msfgfp*; $Tet^R$ | This study |
| $P_{Zn}$*-gfp-ccrZ* | D39V, *bgaA::*$P_{Zn}$*-msfgfp-spv_0476*; $Tet^R$ | This study |
| $P_{Zn}$*-ftsZ-gfp* | D39V, *bgaA::*$P_{Zn}$*-ftsZ-msfgfp*; $Tet^R$ | This study |
| R6 $P_{Zn}$*-gfp-ccrZ* | R6, *bgaA::*$P_{Zn}$*-msfGFP-spv_0476;* Tet$^R$ | This study |
| TIGR4 $P_{Zn}$*-gfp-ccrZ* | TIGR4, *bgaA::*$P_{Zn}$*-msfGFP-spv_0476;* $Tet^R$ | This study |
| *gfp-ccrZ* | D39V, *spv_0476::msfgfp-spv_0476*; $Kan^R$ | This study |
| *ftsZ-mCherry* | D39V, *ftsZ::ftsZ-mCherry*; $Kan^R$ | 1 |
| *gfp-ccrZ ftsZ-mCherry* | D39V, *spv_0476::msfgfp-spv_0476*, *ftsZ::ftsZ-mCherry*; $Kan^R$, $Ery^R$ | This study |
| *ftsZ*$^{+/+}$ | D39V, *prs1::lacI*, *cil::*$P_{lac}$*-ftsZ*; $Gen^R$, $Kan^R$ | This study |
| *ftsZ*$^{-/+}$ | D39V, *prs1::lacI*, *cil::*$P_{lac}$*-ftsZ*, Δ*ftsZ*; $Gen^R$, $Kan^R$, $Spc^R$ | This study |
| *ftsZ-mKate2* | D39V, *ftsZ::ftsZ-mKate2*; $Ery^R$ | 1 |
| *ccrZ-mKate2* | D39V, *spv_0476::spv_0476-mKate2*; $Ery^R$ | This study |
| *ftsZ*$^{-/+}$ *ccrZ-mKate2* | D39V, *prs1::lacI*, *cil::*$P_{lac}$*-ftsZ*, Δ*ftsZ*, *spv_0476::spv_0476-mKate2*; $Gen^R$, $Spc^R$, $Kan^R$, $Ery^R$ | This study |
| *ccrZ-LgBit* | D39V, *spv_0476::spv_0476-LgBit*; $Ery^R$ | This study |
| *ftsZ-SmBit* | D39V, *ftsZ::ftsZ-SmBit*; $Spc^R$ | This study |
| *ccrZ-LgBit ftsZ-SmBit* | D39V, *spv_0476::spv_0476-LgBit*, *ftsZ::ftsZ-SmBit*; $Ery^R$, $Spc^R$ | This study |
| *ccrZ-LgBit ccrZ-SmBit* | D39V, *spv_0476::spv_0476-LgBit-spv_0476-SmBit*; $Ery^R$, $Spc^R$ | This study |
| *ccrZ-LgBit cps2E-SmBit* | D39V, *spv_0476::spv_0476-LgBit*, *cps2E::cps2E-SmBit*; $Ery^R$, $Spc^R$ | This study |
| *ccrZ-LgBit hlpA-SmBit* | D39V, *spv_0476::spv_0476-LgBit*, *hlpA::hlpA-SmBit*; $Ery^R$, $Spc^R$ | This study |
| *ccrZ-LgBit dnaA-SmBit* | D39V, *spv_0476::spv_0476-LgBit*, *dnaA::dnaA-SmBit*; $Ery^R$, $Spc^R$ | This study |

| | | |
|---|---|---|
| *ccrZ-LgBit ezrA-SmBit* | D39V, *spv_0476*::*spv_0476-LgBit*, *ezrA*::*ezrA-SmBit*; Ery$^{R}$, Spc$^{R}$ | This study |
| *ccrZ-LgBit ftsA-SmBit* | D39V, *spv_0476*::*spv_0476-LgBit*, *ftsA*::*ftsA-SmBit*; Ery$^{R}$, Spc$^{R}$ | This study |
| *ccrZ-LgBit ftsH-SmBit* | D39V, *spv_0476*::*spv_0476-LgBit*, *ftsH*::*ftsH-SmBit*; Ery$^{R}$, Spc$^{R}$ | This study |
| *ccrZ-LgBit ftsW-SmBit* | D39V, *spv_0476*::*spv_0476-LgBit*, *ftsW*::*ftsW-SmBit*; Ery$^{R}$, Spc$^{R}$ | This study |
| *ccrZ-LgBit pepN-SmBit* | D39V, *spv_0476*::*spv_0476-LgBit*, *pepN*::*pepN-SmBit*; Ery$^{R}$, Spc$^{R}$ | This study |
| *ccrZ-LgBit pbp2x-SmBit* | D39V, *spv_0476*::*spv_0476-LgBit*, *pbp2x*::*pbp2x-SmBit*; Ery$^{R}$, Spc$^{R}$ | This study |
| *ccrZ-LgBit zapA-SmBit* | D39V, *spv_0476*::*spv_0476-LgBit*, *zapA*::*zapA-SmBit*; Ery$^{R}$, Spc$^{R}$ | This study |
| *ccrZ-LgBit fruA-SmBit* | D39V, *spv_0476*::*spv_0476-LgBit*, *fruA*::*fruA-SmBit*; Ery$^{R}$, Spc$^{R}$ | This study |
| *ccrZ-LgBit plsC-SmBit* | D39V, *spv_0476*::*spv_0476-LgBit*, *plsC*::*plsC-SmBit*; Ery$^{R}$, Spc$^{R}$ | This study |
| *ccrZ-LgBit scrA-SmBit* | D39V, *spv_0476*::*spv_0476-LgBit*, *scrA*::*scrA-SmBit*; Ery$^{R}$, Spc$^{R}$ | This study |
| *ccrZ-LgBit spv_1621-SmBit* | D39V, *spv_0476*::*spv_0476-LgBit*, *spv*_1621::*spv*_1621-*SmBit*; Ery$^{R}$, Spc$^{R}$ | This study |
| *comCDE-LgBit* | D39V, *prs1*::$P_{comC}$-*comC-comD-comE-LgBit*; Cam$^{R}$ | Veening Lab collection |
| *hlpA-LgBit* | D39V, *prs1*::$P_{hlpA}$-*hlpA-LgBit*; Cam$^{R}$ | This study |
| *hlpA-LgBit hlpA-SmBit* | D39V, *prs1*::$P_{hlpA}$-*hlpA-LgBit*, *hlpA*::*hlpA-SmBit;* Cam$^{R}$, Spc$^{R}$ | This study |
| D39V *lacI ccrZ-LgBit* | D39V, *prs1*::*lacI, spv_0476*::*spv_0476-LgBit;* Ery$^{R}$, Gen$^{R}$ | This study |
| *ccrZ-LgBit* $P_{lac}$-*ftsZ* | D39V, *prs1*::*lacI, spv_0476*::*spv_0476-LgBit, cil*::$P_{lac}$-*ftsZ*; Gen$^{R}$, Ery$^{R}$, Kan$^{R}$ | This study |
| $P_{lac}$-*dnaA-SmBit* | D39V, *prs1*::*lacI*, *zip*::$P_{lac}$-*dnaA-SmBit*; Gen$^{R}$, Spc$^{R}$ | This study |
| $P_{lac}$-*dnaA-SmBit yabA-LgBit* | D39V, *prs1*::*lacI*, *zip*::$P_{lac}$-*dnaA-SmBit, yabA*::*yabA-LgBit*; Gen$^{R}$, Spc$^{R}$, Ery$^{R}$ | This study |
| *yabA-LgBit* | D39V, *yabA*::*yabA-LgBit*; Ery$^{R}$ | This study |
| *yabA-LgBit ccrZ-SmBit* | D39V, *yabA*::*yabA-LgBit, spv_0476*::*spv_0476-SmBit*; Spc$^{R}$, Ery$^{R}$ | This study |
| *P3-gfp* | D39V, *cep*::*P3-sfgfp*; Spc$^{R}$ | 5 |
| *hlpA-mKate2* | D39V, *hlpA*::*hlpA-hlpA-mKate2*; Cam$^{R}$ | 23 |
| *hlpA-mKate2* Δ*ccrZ* | D39V, *hlpA*::*hlpA-hlpA-mKate2*; Δ*spd_0476*; Cam$^{R}$, Ery$^{R}$ | 1 |
| *ftsZ-cfp* | D39V, *ftsZ*::*ftsZ-cfp*; Kan$^{R}$ | 1 |
| $P_{Zn}$-*ccrZ*$^{+/+}$ *ftsZ-cfp* | D39V, *bgaA*::$P_{Zn}$-*spv_0476*, *ftsZ*::*ftsZ-cfp*; Tet$^{R}$, Kan$^{R}$ | This study |
| $P_{Zn}$-*ccrZ*$^{+/+}$ *ftsZ-cfp hlpA-mKate2* | D39V, *bgaA*::$P_{Zn}$-*spv_0476*, *ftsZ*::*ftsZ-cfp, hlpA*::*hlpA-hlpA-mKate2*; Tet$^{R}$, Kan$^{R}$, Cam$^{R}$ | This study |
| $P_{Zn}$-*ccrZ*$^{-/+}$ *ftsZ-cfp hlpA-mKate2* | D39V, *bgaA*::$P_{Zn}$-*spv_0476*, *ftsZ*::*ftsZ-cfp, hlpA*::*hlpA-hlpA-mKate2*, Δ*spd_0476*; Tet$^{R}$, Kan$^{R}$, Cam$^{R}$, Ery$^{R}$ | This study |
| *ccrZ*$^{supp1}$ | D39V, Δ*spd_0476*, *dnaA*::*dnaA-Q247H,* 13_14insA *licD2*; Ery$^{R}$ | This study |
| *ccrZ*$^{supp2}$ | D39V, Δ*spd_0476*, *dnaA*::*dnaA-S292G*; Ery$^{R}$ | This study |
| *ccrZ*$^{supp3}$ | D39V, Δ*spd_0476*, *yabA*::*yabA-E93**; Ery$^{R}$ | This study |
| Δ*ccrZ dnaA-Q247H* | D39V, Δ*spd_0476*, *dnaA*::*dnaA-Q247H*; Ery$^{R}$ | This study |
| Δ*ccrZ dnaA-S292G* | D39V, Δ*spd_0476*, *dnaA*::*dnaA-S292G*; Ery$^{R}$ | This study |
| *kan-ccrZ* | D39V, *spv_0476*::*kan-spv_0476*; Kan$^{R}$ | This study |
| *dnaA-Q247H* | D39V, *spv_0476*::*kan-spv_0476*, *dnaA*::*dnaA-Q247H*; Kan$^{R}$ | This study |
| *dnaA-S292G* | D39V, *spv_0476*::*kan-spv_0476*, *dnaA*::*dnaA-S292G*; Kan$^{R}$ | This study |
| Δ*yabA* | D39V, Δ*yabA*; Spc$^{R}$ | This study |
| Δ*yabA* Δ*ccrZ* | D39V, Δ*yabA*, Δ*spd_0476*; Spc$^{R}$, Ery$^{R}$ | This study |
| *ccrZ-N164A* | D39V, *bgaA*::$P_{Zn}$-*spv_0476*, *spv_0476*::*kan-spv_0476-N164A*; Tet$^{R}$, Kan$^{R}$ | This study |

| | | |
|---|---|---|
| *ccrZ-H157A* | D39V, *bgaA*::$P_{Zn}$-*spv_0476*, *spv_0476*::*kan-spv_0476-H157A*; $Tet^R$, $Kan^R$ | This study |
| *ccrZ-D177A* | D39V, *bgaA*::$P_{Zn}$-*spv_0476*, *spv_0476*::*kan-spv_0476-D177A*; $Tet^R$, $Kan^R$ | This study |
| $P_{lac}$-*ccrZ-gfp* | D39V, *prs1*::*lacI*, *zip*::$P_{lac}$-*spv_0476-msfgfp*; $Gen^R$, $Spc^R$ | This study |
| $P_{lac}$-*ccrZ-H157A-gfp* | D39V, *prs1*::*lacI*, *zip*:: $P_{lac}$-*spv_0476-H157A-msfgfp*; $Gen^R$, $Spc^R$ | This study |
| $P_{lac}$-*ccrZ-N164A-gfp* | D39V, *prs1*::*lacI*, *zip*:: $P_{lac}$-*spv_0476-N164A-msfgfp*; $Gen^R$, $Spc^R$ | This study |
| $P_{lac}$-*ccrZ-D177A-gfp* | D39V, *prs1*::*lacI*, *zip*:: $P_{lac}$-*spv_0476-D177A-msfgfp*; $Gen^R$, $Spc^R$ | This study |
| *comCDE-parS*$_p$ *ccrZ-mKate2* | D39V, *spv_0476::spv_0476-mKate2, comCDE::comCDE-parS*$_p$; $Ery^R$ | This study |
| *parB*$_p$*-YFP comCDE-parS*$_p$ *ccrZ-mKate2* | D39V, *spv_0476::spv_0476-mKate2, comCDE::comCDE-parS*$_p$, *bgaA::*$P_{Zn}$-*parB*$^p$*-msfYFP*; $Ery^R$, $Tet^R$ | This study |
| *dnaN-mTQ*$^{ox}$ *parB*$_p$*-YFP comCDE-parS*$_p$ *ccrZ-mKate2* | D39V, *spv_0476::spv_0476-mKate2, comCDE::comCDE-parS*$_p$, *bgaA::*$P_{Zn}$-*parB*$^p$*-msfYFP,* ; $Ery^R$, $Tet^R$, $Cam^R$ | This study |
| *comCDE-parS*$_p$ *parB*$_p$*-GFP* | D39V, *comCDE-parS*$_p$ *bgaA::parB*$_p$*-sfmGFP*; $Kan^R$ | 1 |
| *spv_1159-mTurquoise2*$^{ox}$ | D39V, *CEP*::*P3-spv_1159-sfmTurquoise2*$^{ox}$*-opt;* $Spc^R$ | Veening Lab collection |
| *ccrZ-mKate2* $P_{Zn}$*-dnaA-GFP* | D39V, *ccrZ*::*ccrZ-mKate2, bgaA*::$P_{Zn}$*-dnaA-msfGFP;* $Ery^R$, $Tet^R$ | This study |
| $P_{tet}$-*dCas9* | *D39V, prs1::PF6-lacI-tetR, bgaA::Ptet-dcas9; Gen*$^R$*, Tet*$^R$ | This study |
| $P_{tet}$-*dCas9,* $P_{lac}$-*ccrZ* | *D39V, prs1::PF6-lacI-tetR, bgaA::Ptet-dcas9, cil::Plac-spv_0476; Gen*$^R$*, Tet*$^R$*, Kan*$^R$ | This study |
| $P_{tet}$-*dCas9,* $P_{lac}$-*ccrZ,* Δ*ccrZ* | *D39V, prs1::PF6-lacI-tetR, bgaA::Ptet-dcas9, cil::Plac-spv_0476, Δspd_0476; Gen*$^R$*, Tet*$^R$*, Kan*$^R$*, Ery*$^R$ | This study |
| ***S. aureus*** **strains** | | |
| SH1000 | *S. aureus* SH1000 | 24 |
| SH1000 *dcas9* | SH1000, *pLOW-dcas9;* $Ery^R$ | 12 |
| *ccrZ*$_{Sa}^{sgRNA}$ | SH1000, *pLOW-dcas9*, *pCG248-sgRNA(ccrZ*$_{Sa}$*);* $Ery^R$, $Cam^R$ | This study |
| *ccrZ*$_{Sa}^{sgRNA}$ $P_{ccrZSa}$-*ccrZ*$_{Bs}$ | SH1000, *pLOW-dcas9*, *pCG248-sgRNA(ccrZ*$_{Sa}$*)-*$P_{ccrZSa}$*-ccrZ*$_{Bs}$*;* $Ery^R$, $Cam^R$ | This study |
| $P_{lac}$-*gfp-ccrZ*$_{Sa}$ | SH1000, *pLOW-m(sf)gfp-ccrZ*$_{Sa}$*;* $Ery^R$ | This study |
| ***B. subtilis*** **strains** | | |
| 1A700 | *B. subtilis* 1A700 | Gruber lab collection |
| 1A700 Δ*ccrZ*$_{Bs}$ | 1A700, Δ*ytmP*; $Ery^R$ | This study |
| *ccrZ*$_{Bs}$*-gfp* | 1A700, *ytmP-msfgfp*; $Ery^R$ | This study |
| JH642 | *B. subtilis* JH642 | 25 |
| JH642 Δ*ccrZ*$_{Bs}$ | JH642, Δ*ytmP*; $Cam^R$ | This study |
| *oriN* | JH642, *oriC::oriN-repN, Ppen-dnaN*; $Kan^R$ | This study |
| *oriN,* Δ*ccrZ*$_{Bs}$ | JH642, *oriC::oriN-repN, Ppen-dnaN, ΔytmP*; $Cam^R$; $Can^R$ | This study |
| *dnaB134* | *dnaB* temperature sensitive mutant | Grossman lab collection |
| **Plasmids** | | |
| pPEPY | Vector with *Plac,* allowing integration into *S. pneumoniae* CIL locus (*spv_2166-spv_2165*); $Kan^R$ | 26 |
| pPEPZ | Vector with *Plac,* allowing integration into *S. pneumoniae* ZIP locus (spv_2416-*spv_2419*); $Spc^R$ | 26 |
| pCG6 | pMK17 derivative, encoding *ccrZ* fused to 3'of *msfyfp;* $Tet^R$ | 3 |

| | | |
|---|---|---|
| pMK11 | Vector with $P_{Zn}$ ($P_{czcD}$) allowing integration into *S. pneumoniae bgaA* locus; $Tet^R$ | 1 |
| pMK17 | pMK11 derivative, encoding *msfgfp;* $Tet^R$ | 1 |
| pMK19-02 | pMK17 derivative, encoding *parB* fused to *msfyfp;* $Tet^R$ | Veening Lab collection |
| pLOW-dcas9 | *dcas9* downstream of $P_{lac}$ promoter; $Amp^R$, $Ery^R$ | 12 |
| *pCG248-sgRNA(ccrZ$_{Sa}$)* | For constitutive expression of sgRNA(*ccrZ$_{Sa}$*); $Amp^R$, $Cam^R$ | This study |
| *pCG248-sgRNA(ccrZ$_{Sa}$)-P$_{ccrZSa}$-ccrZ$_{Bs}$* | For constitutive expression of sgRNA(*ccrZ$_{Sa}$*), *ccrZ$_{Bs}$* (*ytmP*) expressed from $P_{ccrZSa}$; $Amp^R$, $Cam^R$ | This study |
| pLOW-*msggfp-parB* | *msfgfp-parB* fusion downstream of $P_{lac}$ promoter; $Amp^R$, $Ery^R$ | Kjos Lab collection |
| pGK67 | pGEM derivative, carrying a kanamycin resistance cassette; $Cam^R$ | 18 |
| pDL110 | pJH101 derivative, carrying *oriN-repN* from pBPA23; $Amp^R$ | 27 |
| pUT18 | Encoding CyaA T18 domain under control of $P_{lac}$; $Amp^R$ | 28 |
| pST25 | Encoding CyaA T25 domain under control of $P_{lac}$; $Spc^R$ | 28 |
| pUC19 | pUC18 derivative, inverted MCS, $Amp^R$ | Addgene |
| pJET1.2 | pJET1.2 | Invitrogen |
| pSG1525 | pET-Gate2 derivative with MazEF toxin-antitoxin system for Golden gate assembly selection; $Kan^R$ | 14 |
| pSG0682 | pJET1.2 derivative with erythromycin (*ermB*) cassette; $Ery^R$ | 14 |
| pSG3174 | pUC19 derivative, carrying *ccrZ$_{Bs}$* (*ytmP*); $Amp^R$ | This study |
| pSG3177 | pUC19 derivative, carrying sequence of upstream region of *ccrZ$_{Bs}$* (*ytmP*); $Amp^R$ | This study |
| pSG3178 | pUC19 derivative, carrying sequence of downstream region of *ccrZ$_{Bs}$* (*ytmP*); $Amp^R$ | This study |
| pSG3175 | pUC19 derivative, carrying *ccrZ$_{Bs}$* (*ytmP*) with no STOP codon for fusion with *msfgfp*; $Amp^R$ | This study |
| pSG3176 | pUC19 derivative, carrying sequence of downstream region of *ccrZ$_{Bs}$* (*ytmP*); $Amp^R$ | This study |
| pSG3179 | pUC19 derivative, carrying *msfgfp*; $Amp^R$ | This study |
| pSG2949 | pET-Gate2 derivative, encoding *ccrZ* with C-terminal cysteine protease domain (CPD) with a 10his tag ; $Kan^R$ | This study |
| pSG2950 | pET-Gate2 derivative, encoding *ccrZ* fused to *sfgfp* with a 10his tag ; $Kan^R$ | This study |
| pSG4227 | pJET1.2 derivative, carrying *ftsZ* from *S. pneumoniae*; $Amp^R$ | This study |
| pSG1694 | pET-Gold1 derivative encoding a *ccdB* cassette, used for Golden gate assembly ; $Amp^R$ | Gruber Lab collection |
| pSG4268 | pETGold1 derivative encoding *ftsZ* from *S. pneumoniae ;* $Amp^R$ | This study |
| pSG436 | pET-Gate2 derivative encoding *ccdB* cassette ; $Kan^R$ | Gruber Lab collection |
| pSG366 | pJET1.2 derivative carrying short sequence2 for Golden gate assembly ; $Amp^R$ | Gruber Lab collection |
| pSG367 | pJET1.2 derivative carrying short sequence3 for Golden gate assembly ; $Amp^R$ | Gruber Lab collection |
| pSG2559 | pET-Gold1 derivative carrying C-terminal cysteine protease domain (CPD) with a 10his tag; $Amp^R$ | Gruber Lab collection |
| pSG2562 | pET-Gold1 derivative carrying *sfGFP* with a 10his ; $Amp^R$ | Gruber Lab collection |
| ***E. coli* strains** | | |

| | | |
|---|---|---|
| HM1784 | BTH101 derivative, $F^-$, *cya-99*, *araD139*, *galE15*, *galK16*, *rpsL1*, *hsdR2*, *mcrA1*, *mcrB1,* Δ*rnh*::*kan*; $Str^R$ | 28 |
| IM08B | K12 DH10B derivative, *mcrA,* Δ(*mrr-hsdRMS-mcrBC*), φ80*lacZ*Δ*M15,* Δ*lacX74, recA1, araD139,* Δ(*ara-leu*)7697, *galU, galK, rpsL, endA1, nupG,* Δ*dcm*, Ω*Phelp-hsdMS* (CC8-2), Ω*PN25-hsdS* (CC8-1); $Str^R$ | 29 |
| BL21 DE3 Gold | B $F^-$, *ompT, hsdS*($r_B^-$ $m_B^-$), *dcm*$^+$, *gal*λ(DE3), *endA,* Hte; $Tet^R$ | Agilent Technologies |

Abbreviations: $Tet^R$: tetracycline resistance; $Kan^R$: kanamycin resistance; $Cam^R$: chloramphenicol resistance; $Ery^R$: erythromycin resistance; $Spc^R$: spectinomycin resistance; $Gen^R$: gentamycin resistance; $Amp^R$: ampicillin resistance; SmBit: Small Bit; LgBit: Large Bit

**Supplementary Table 2: Oligonucleotides used in this study**

| Primer name | forward / reverse | Sequence (5' -> 3') |
|---|---|---|
| 1 | F | CGATGGAATTCGAAACTTTATACGGAGGAAAGAAATG |
| 2 | R | GCGCACTAGTTCATCTTCTCTTTCCATACTTGTCT |
| 3 | F | CACTTATTTTTCCTAGATTCCA |
| 4 | R | TGTTCATATGAAAATTCCTCCGGGCGCGACTTCTTTCCTCCGTATAAAGTTTCCT |
| 5 | F | AGTTATCTATTATTTAACGGGAGGAAATAAGACAAGTATGGAAAGAGAAGATGAG |
| 6 | R | CATATATTCCATAGTCAACTC |
| 7 | F | GTCGCGCCCGGAGGAATTTTCATATG |
| 8 | R | TTATTTCCTCCCGTTAAATAATAGA |
| 9 | R | GCGCGGATCCTCATCTTCTCTTTCCATACTTGTCT |
| 10 | F | CATTTCTTCGCTGTTTCTTC |
| 11 | R | GGAAATATTGTTCTGGGAAG |
| 12 | F | CGGATTTCCAACAAGCTTCA |
| 13 | R | GGAACGTCGTCTCCGTTTTAACGATTTTTGAAAAATGGAGG |
| 14 | F | GGAACGTCGTCTCGTAGTTTTCAAAAATCGTTAAGTAAATGAATG |
| 15 | R | CCAAATAGTCCAAACAACGAC |
| 16 | F | GGAACGTCGTCTCGAAACGAGGTGAAATCATGAGC |
| 17 | R | GGAACGTCGTCTCGACTAATTGAGAGAAGTTTCTATAGAATTTT |
| 18 | R | GGATTCCCGTCTCCACTCCTTTAGCTGCAGCTTC |
| 19 | F | GGATTCCCGTCTCCGAGTGAGCAAGGGCGAGGAG |
| 20 | R | GGATTCCCGTCTCCGTTTTACTTGTACAGCTCGTCC |
| 21 | F | GGATTCCCGTCTCGAAACGAGGTGAAATCATGAGC |
| 22 | F | CAGCCATAGAGGAGATCATCATGTA |
| 23 | R | TGCGCTAGGTCTCCTTTCTTTCCTCCGTATAAAGTTTC |
| 24 | F | TGCGCTAGGTCTCGGAAATGAACAAAAATATAAAATATTCTCAAAACT |
| 25 | R | TGCGCTAGGTCTCGTCTTATTTCCTCCCGTTAAATAATAGAT |
| 26 | F | TGCGCTAGGTCTCGAAGAGATAGCCTAAAATTAGGCTG |
| 27 | R | GGCTGCTACACGACCAAACTCAG |
| 28 | F | GCATGCGGCCGCAAACTTTATACGGAGGAAAGAAATG |
| 29 | R | CGCCACTAGTTCTTCTCTTTCCATACTTGTCTCGG |
| 30 | F | GCATGCGGCCGCATAAAGAGGAAAAATAAATTATGAC |
| 31 | R | GCGCACTAGTACGATTTTTGAAAAATGGAGGTGTA |
| 32 | F | GCGCACTAGTGATTTGGGTGATAATGAGC |
| 33 | R | GCATGCGGCCGCTCATCATCTTCTCTTTCCATACTTGT |
| 34 | F | GTACTGCCTGCGTGGTTATC |
| 35 | R | ACGGATCCCCAGCTTGCGCGTCCTCTTCTCAAAGAAAAGCCTCTGGATTG |
| 36 | R | CTATTCGTCAACCACTCTAACCCGAAGATATAAAACATCAGAGTATGGAC |
| 37 | F | TCTGATGTTTTATATCTTCGGGTTAGAGTGGTTGACGAATAGGCCAAAAACTAGTAGAATAGTAAG GAAACTTTATACGGAGGAAAGAAATGAAACATCTTAGCTCAAAAGGAGAAGAGC |
| 38 | R | TCATCTTCTCTTTCCATACTTGTCTC |
| 39 | F | GAGGACGCGCAAGCTGGGGATCCGT |
| 40 | F | CGATGGAATTCAATAAAGAGGAAAAATAAATTATG |
| 41 | R | GCGCGGATCCTTAACGATTTTTGAAAAATGGAGGTG |
| 42 | F | TGTCCTCCGTCTCCTAACGAGGTGAAATCATGAGC |
| 43 | R | GGAAATTACGGATCAAGATG |
| 44 | R | TGTCCTCCGTCTCCGTTAATTTATTTTTCCTCTTTATTCGTCA |
| 45 | F | GCTATGTGGTCGGATTTTGGTT |
| 46 | R | CAGGACACGTCTCGATCTTCTCTTTCCATACTTGTCT |
| 47 | F | CAGGACACGTCTCGAGATCCGGATCTGGTGGAGAA |
| 48 | R | CAGGACACGTCTCGACTTATTTCCTCCCGTTAAATAATAGAT |
| 49 | F | CAGGACACGTCTCCAAGTATGGAAAGAGAAGATGAGAGT |
| 50 | R | CCAAACGGCTATCAAATACTTCA |
| 51 | F | GCTATGTGGTCGGATTTTGGTT |
| 52 | F | CTTTGAGAATCGGGTTAGAG |

| | | |
|---|---|---|
| 53 | R | GCTGTCAGGTCTCGCCTCTTCTCTTTCCATACTTGTCT |
| 54 | F | GCTGTCAGGTCTCCTAACCCGGAGGAATTTTCA |
| 55 | R | GGCCATGGATCTGAAAAGTTC |
| 56 | F | GGAAGGTCGTCTCCTCGTACAACAGATTCAGTCGTTT |
| 57 | R | TGCCGATCGTCTCCTACATCGCTG |
| 58 | F | GATCGGAAGCATGTTTGACG |
| 59 | R | GGAAGGTCGTCTCCACGAACAATCGATTCACGGCG |
| 60 | F | TGCCGATCGTCTCGTGTAGATGTTCTAGTCGTCGTGAATC |
| 61 | R | CCAGCTAAAACATGACAAGC |
| 62 | F | GACAATTCTTGGTTAGGGCT |
| 63 | R | TGAGGGTCGTCTCCCTCTTCTCTTTCCATACTTGTCT |
| 64 | F | TGAGGGTCGTCTCCAGAGGCGGCTCATCAGGCGGCG |
| 65 | R | TGAGGGTCGTCTCCACCTAATTGAGAGAAGTTTCTATAG |
| 66 | F | TGAGGGTCGTCTCCAGGTATGGAAAGAGAAGATGAGAGT |
| 67 | R | ATTCGGACGGTTTTCTTACGCG |
| 68 | F | GATACCCAAGCCAATCCAGAGGC |
| 69 | R | TGAGGGTCGTCTCGGTTATTTCCTCCCGTTAAATAATAGAT |
| 70 | F | TGAGGGTCGTCTCCTAACTTTATACGGAGGAAAGAAATGG |
| 71 | F | CCCGTATCGTCTCTGGCGGCTCATCAGGCGGCGGC |
| 72 | R | CCCGTATCGTCTCCCTAATTGAGAGAAGTTTCTATAG |
| 73 | F | CCCGTATGGTCTCTGGCGGCTCATCAGGCGGCGGC |
| 74 | R | CCCGTATGGTCTCCCTAATTGAGAGAAGTTTCTATAG |
| 75 | F | CTACTTAGTGCTGTGAGAGAAGC |
| 76 | R | CCCGTATCGTCTCCCGCCCTTCGCTCCATCTCTCA |
| 77 | F | CCCGTATCGTCTCGTTAGTTTACTTTTGTTTTAGACTACTAG |
| 78 | R | GCTTCTGCAATTCGTTGGCTAG |
| 79 | F | AGCATAAATAGCAGCACCTA |
| 80 | R | CCCGTATCGTCTCCCGCCTTTAACAGCGTCTTTAAGAGCTTTACC |
| 81 | F | CCCGTATCGTCTCGTTAGTCAGTCTTTAAAAAGCCTATTGTATC |
| 82 | R | GCGATGGTTTCAATATCCAAGTG |
| 83 | F | GGCTGTATCAGCCGCTTTAGCTGTC |
| 84 | R | CCCGTATGGTCTCCCGCCTTTGATTTTCTTTTTGATTGATTCA |
| 85 | F | CCCGTATGGTCTCGTTAGTTTGTGGATAACTTTTAGTTTTTTATC |
| 86 | R | CACCAGTAACCTGATCAGTATCGA |
| 87 | F | GGCCTTAAGTCATATTGTGGATCGT |
| 88 | R | CCCGTATGGTCTCCCGCCAAAACGAATCGTTTCACGTGTTTTC |
| 89 | F | CCCGTATGGTCTCGTTAGTAAAAGAAAAAGATTTTATTGTGTGAG |
| 90 | R | TGCCTCTGCAGCCTTCAAATGGCGG |
| 91 | F | GATGCAGCAGCAACTGCTATC |
| 92 | R | CCCGTATCGTCTCCCGCCTTCGTCAAACATGCTTCCGATC |
| 93 | F | CCCGTATCGTCTCGTTAGAGAGGAAAAATAAATTATGAC |
| 94 | R | CTCACGGTAGTTAGTAGCAGATCTA |
| 95 | F | GAGGTAATGACGAACGTGAACAAAC |
| 96 | R | CCCGTATCGTCTCCCGCCTTTTTCGTCATTCATTTTTGACTTTAC |
| 97 | F | CCCGTATCGTCTCGTTAGCCCTGAGAGAGGCTGGAGCCTC |
| 98 | R | TTAGGCACGCCGCCAGCGTTCGTCC |
| 99 | F | GCGCCTTGCAGTTGGTTCGAAACC |
| 100 | R | CCCGTATCGTCTCCCGCCCTTCAACAGAAGGTTCATTGGTTG |
| 101 | F | CCCGTATCGTCTCGTTAGGATAAAGAAAGGATAGTTTATGTCTC |
| 102 | R | GAATATCAAACAAGGCACGACGG |
| 103 | F | CACTGAAAATACTGCCCACTATATT |
| 104 | R | CCCGTATCGTCTCCCGCCTGCATTTCCGTATTGAAGAACAACTGC |
| 105 | F | CCCGTATCGTCTCGTTAGATAAGCCTAAAATAAAAAGAAAACTCAGC |
| 106 | R | CGATAGATTTCTTCCTGCCCATAGC |
| 107 | F | GAAGGATTGACTGGTGGCAGAATG |
| 108 | R | CCCGTATCGTCTCCCGCCGTCTCCTAAAGTTAATGTAATTTTT |
| 109 | F | CCCGTATCGTCTCGTTAGTATGTTTATTTCCATCAGTGCTG |

| | | |
|---|---|---|
| 110 | R | CATCAGCTTCGCCTTATCTACAACC |
| 111 | F | GAGATCACGCGCAATGACAGAAGAA |
| 112 | R | CCCGTATCGTCTCCCGCCTAAGGAATCCTCAATCTTGCTCTGT |
| 113 | F | CCCGTATCGTCTCGTTAGCAGAGCAAGATTGAGGATTCCTTATG |
| 114 | R | CGACGGATTCGCGCCAATTCTTCAC |
| 115 | F | GATGAGTGGTGTATCTCAAATG |
| 116 | R | CCCGTATCGTCTCCCGCCTGCTTGTGGTTTGCGTAGGTAACCA |
| 117 | F | CCCGTATCGTCTCGTTAGAAAAATAGAAAAATGAAAAGATTGG |
| 118 | R | CAAAGCTAGCAAAGGTTGCTTCTAA |
| 119 | F | GGTCATGAGATTGAGCTTGTCC |
| 120 | R | CCCGTATCGTCTCCCGCCTGCAAGTTCTTCTCTTTTCTTATCT |
| 121 | F | CCCGTATCGTCTCGTTAGAAGAAATGAACCTTGGCCAAACAGC |
| 122 | R | TTCCTACGTTCGACATTACCCACTA |
| 123 | F | CATGTCTATCCTTGGACTCTTTGTC |
| 124 | R | CCCGTATCGTCTCCCGCCGATTTTCACTTCGATCACAGCATCCC |
| 125 | F | CCCGTATCGTCTCGTTAGTCAATCCTCTCTAATGTGAAAACG |
| 126 | R | GTCGCTACCGTCCGTAATCACTTAA |
| 127 | F | GACCTGCCATGAGTTTGTTGAAACT |
| 128 | R | CCCGTATCGTCTCCCGCCAAGAGTATAGGCCATGGCCCCTGC |
| 129 | F | CCCGTATCGTCTCGTTAGAAATCTCTTTAAACCATGTCAGC |
| 130 | R | GCGGCTGTCGCAGAGTTGAGACAAA |
| 131 | F | ATGTCTTTTTCTGATTTAAAGCTGTTTGCCC |
| 132 | R | CTATTTCTTCCAATTTTATTTATTTTGAAGGCGAATGCTCTATCCAGC |
| 133 | F | GCTGGATAGAGCATTCGCCTTCAAAATAAATAAAATTGGAAGAAATAG |
| 134 | R | GCCGCCGCCGCCTGATGAGCCGCCTTTAACAGCGTCTTTAAGAGCTTTAC |
| 135 | F | GTAAAGCTCTTAAAGACGCTGTTAAAGGCGGCTCATCAGGCGGCGGCGGC |
| 136 | R | CTTGTCCCCACTGACAATGGTAATATC |
| 137 | F | GATTGTAACCGATTCATCTG |
| 138 | R | GGAATGCTTGGTCAAATCTA |
| 139 | F | CAAGCAAAAACATATTGTCCTAACG |
| 140 | R | CGTTAGGACAATATGTTTTTGCTTG |
| 141 | F | GCCATTTTACAACGTAAAACGGAAC |
| 142 | R | GTTCCGTTTTACGTTGTAAAATGGC |
| 143 | F | TCTTCTTTTATCCCCAACCTG |
| 144 | R | GAGTCTGTGAAATCTGTGGA |
| 145 | R | ACCCTGACGTCTCGTTTCTTTCCTCCGTATAAAGTTTC |
| 146 | F | ACCCTGACGTCTCCGAAATGGATTTGGGTGATAATGAGC |
| 147 | F | GCTACTTCTCTATATTGCCAGTC |
| 148 | R | CACGAACCGTCTCGCATAAAACAGCCCTTTCCTTTCTTTATCATTC |
| 149 | F | CACGAACCGTCTCGTATGAGCAATTTGATTAACGGAAAAATAC |
| 150 | R | CACGAACCGTCTCGGCCTAATTGAGAGAAGTTTCTATAGAATTTTTC |
| 151 | F | CACGAACCGTCTCCAGGCATGCAGATTCAAAAAAGTTTTAAGG |
| 152 | R | CCCTTTCTTTCATCTTCTTCCC |
| 153 | F | GTACGACATAGTGCTTGGATTGAGAC |
| 154 | R | GTCTCAATCCAAGCACTATGTCGTAC |
| 155 | F | GCGACCATTGTCGCTGGAGATGTACG |
| 156 | R | CGTACATCTCCAGCGACAATGGTCGC |
| 157 | F | GATTTATTTAGTAGCTTGGGATTCGG |
| 158 | R | CCGAATCCCAAGCTACTAAATAAATC |
| 159 | R | GCGTCACGTCTCAGCATTATTTTTCCTCCTTATTTAT |
| 160 | F | GCGTCACGTCTCACGGATCCCTCCAGTAACTCGAGAA |
| 161 | F | GCGTCACGTCTCAATGCAAACTTTATACGGAGGAAAGAAATGG |
| 162 | R | GCGTCACGTCTCATCCGGCTTACTTATAAAGCTCATCCATGCC |
| 163 | F | GCCAATAAATTGCTTCCTTGTTTTG |
| 164 | R | AATAATTCTTTCTTTGACACCGTATATG |
| 165 | F | GATGTTATTGAACATCAAGTGAGACTAGAG |
| 166 | R | AATCGCCATCTTCCAATCCC |

| | | |
|---|---|---|
| 167 | F | AATTGCGGCCGCAGGAGGTTATCATGTCAATTTATCAAGAATTTG |
| 168 | R | AGTGGGTCTCCCTTTCACTTAATTTGTACGAACTGG |
| 169 | F | AGTGGGTCTCGAAAGGAGAATCCATGATTCATTTTT |
| 170 | R | AGTGGGTCTCGATCCATTTGTACGAACTGGTGTAATG |
| 171 | F | GGTTGCTCTTCGGAAAGAGGTTGAGCCTGGCTC |
| 172 | R | CAAAGGCGTCTTCACGTCCTTG |
| 173 | F | TCACGGTCTCGGGATCCGGATCTGGTGGA |
| 174 | R | TCACGGTCTCGTAGCTTATTATTATTTGTATAGTTCGTCCATGC |
| 175 | F | TCACGGTCTCCGCTAGTAGGAGGCATATCAAATGA |
| 176 | R | TGAGGCTCTTCGTTCTGCCGCTTATAAAAGCCAG |
| 177 | F | GCATGCGGCCGCAATAGTAAAGGAGGAGAAAGGATTG |
| 178 | R | CGCCACTAGTTTTGATTTTCTTTTTGATTGATTCA |
| 179 | R | TATAGTTATTATACCAGGGGGACAGTGC |
| 180 | F | GCTGATAATGCCGCAATAAAGTTTAAGAGCTATGCTGGAAACAG |
| 181 | F | TCGAGGATCCTAGAGTTTATCGCCTACAGAG |
| 182 | R | GTAATTGTCCCAACCAGTTCAT CTCGTCCACCTCACTTTCAA |
| 183 | F | TTGAAAGTGAGGTGGACGAG ATGAACTGGTTGGGACAATTAC |
| 184 | R | ACTGGGTACCTTAATCAACAATTCGTTTCATGAGGA |
| 185 | F | GATCCCATGGAGCAGTTTTATCAATTAGG |
| 186 | R | TCCGGGGATCCAAATAAACATGTTACTATTCATAACT |
| 187 | F | GTTACAGAATTCGGTCTCGCGAGTAGCACCGATTGTTCCGGAACC |
| 188 | R | GTTACATCTAGAGGTCTCGTATTTCAAGCAAAACTCTTAAGATCTGATGC |
| 189 | F | GTTACAGAATTCGGTCTCGAGGTCATATCCTCATGAAACGAATTGTTG |
| 190 | R | GTTACATCTAGAGGTCTCTATGGCAATATCTGCGTTTTCAGCC |
| 191 | R | GTTACAGAATTCGGTCTCGCCTGTAATCAGGAATACGAATCCATTTGC |
| 192 | R | GTTACATCTAGAGGTCTCGACCTAGTAATTGTCCCAACCAGTTCAT |
| 193 | F | GTTACAGAATTCGGTCTCGTCTACGAAGTTTGTGCTATAGCGCTAAAATTTAAAG |
| 194 | R | GTTACATCTAGAGGTCTCGCTCCATCAACAATTCGTTTCATGAGG |
| 195 | F | GTTACAGAATTCGGTCTCAGGAGCAGGCGGTAGCGGCAGCGGTGGCTCA |
| 196 | R | GTTACATCTAGAGGTCTCTCAGGTCATTTGTATAGTTCATCCATGC |
| 197 | F | CAAATCGAGACTGACGTATGTCAG |
| 198 | R | TCAGGAATACGAATCCATTTGCTC |
| 199 | F | GGGTAGAGCAAATGGATTCGTATTCCTGAGGCTTAACTATGCGGCATCAG |
| 200 | R | GCTGGATAGACTAGAAACGATTAGCCCGGTATTTTCTCCTTACGCATCTG |
| 201 | F | GGCTAATCGTTTCTAGTCTATCCAG |
| 202 | R | CAAGCTGTGATGCATGGAATTG |
| 203 | F | TACGCCAACCATACTTAATAGCA |
| 204 | R | CGCAACTGTCCATACTCTGATGTCTATTATGGTTGCAAGAAATAAAAG |
| 205 | F | CTTTTATTTCTTGCAACCATAATAGACATCAGAGTATGGACAGTTGCG |
| 206 | R | GGAAAAGATTTTAGGAGGAAGCTGAACCATTTGAGGTGATAGGTAAG |
| 207 | F | GTTGAACTAATGGGTGCAGCTTCCTCCTAAAATCTTTTCCCAT |
| 208 | R | CGTTTTTTCGGAAAGAAGAATATGTAGAAGAAGTTATTGATG |
| 209 | F | CTTCTACATATTCTTCTTTCCGAAAAAACGGTTGCATTTA |
| 210 | R | TCCTCCTAACGGATAATGTATGCAGCCGACTCAAACATCAAATC |
| 211 | F | GCTGCATACATTATCCGTTAGGAGGATAAAAATGAAATTCACGAT TCAAAAAGATCGTC |
| 212 | R | CTGTATCAATTGAAATCGGATTTGC |
| 213 | F | CTCGAGGGTCTCACGAGAATAATTTTGTTTAACTTTAAGAAGGAGATATACATATGGATTTGGGTGA TAATGAGCTAACACTG |
| 214 | R | GCTGCAGGTCTCAGCCAGCTCTTCTCTTTCCATACTTGTCTCGGAAATG |
| 215 | F | GCAGCCCGTCTCATATGATGACATTTTCATTTGATACAGCTGCTGC |
| 216 | R | CGTCTCACAGGTTAACGATTTTTGAAAAATGGAGGTGTATCC |
| 217 | F | GCATCAACATCTTGTGTGGCACG |
| 218 | R | GCTTGGTTAATCAAACACGCCC |
| 219 | F | GCACTCAAACCTAGAAGAGC |
| 220 | R | CAACTCACATGAACTACATGATG |
| 222 | F | GCCAATAAATTGCTTCCTTGTTTTG |
| 223 | R | GCGTCACGTCTCAGCATTATTTTTCCTCCTTATTTAT |

| | | |
|---|---|---|
| 224 | F | GCGTCACGTCTCACGGATCCCTCCAGTAACTCGAGAA |
| 225 | R | AATAATTCTTTCTTTGACACCGTATATG |
| 226 | F | GCGTCACGTCTCAATGCAATAGTAAAGGAGGAGAAAGGATTG |
| 227 | R | GCGTCACGTCTCATCCGTTATTATAGGATTTCTTCGA |
| 228 | F | CCCTGTACGTCTCCagggctcatcaggcggcggcg |
| 229 | R | CCCTGTACGTCTCCgcTTATTTCCTCCCGTTAAATAATAGATAAC |
| 230 | F | gatgtcaccttgattaagccag |
| 231 | R | CCCTGTACGTCTCCccctccctgtatagcaactcg |
| 232 | F | CCCTGTACGTCTCGAAgcatgcagattcaaaaaagtt |
| 233 | R | gtcaacttcatcagctctcttatct |
| OT1 | F | ACGGTCTATCCCAGCTGTTG |
| OT2 | R | ATAGGCGCGTGCTTCTTCTA |
| OT3 | F | GAAAAGTACCATCCCCAGCA |
| OT4 | R | AGCCTTGGTGCCTATCATTG |
| OT5 | F | TGCATATCCGCGTCAAATAG |
| OT6 | R | GCATGAATGACGGTCGTATG |
| OT7 | F | CGCGTGTAGGTATTGCGTTA |
| OT8 | R | CTTCGCGCACTTGAATAACA |
| OT9 | F | TTGCCGCAGATTGAAGAG |
| OT10 | R | AGGTGGACACTGCAAATAC |
| OT11 | F | CGCGCTGACTCTGATATTATG |
| OT12 | R | CAAAGAGGAGCTGCTGTAAC |

**Supplementary Table 3: Ratio of spectral counts between GFP-CcrZ and GFP from LC-MS/MS**

| Identified Proteins | Fold change (GFP-CcrZ / GFP) |
|---|---|
| SPV_0476 (CcrZ) | 27 |
| ScrA | 12 |
| PepN | 8.5 |
| Pbp2X | 8.3 |
| FruA | 7.4 |
| EzrA | 7.4 |
| SPV_1621 | 6.8 |
| FtsZ | 6.6 |
| PlsC | 6 |
| FtsH | 5.1 |
| DnaA | 2.8 |
