## Supplementary Data S1 for "Spatio-temporal control of DNA replication by the pneumococcal cell cycle regulator CcrZ"

**Data S1. CRISPRi-seq results for ccrZ-complementation vs ccrZ-depletion**

| sgRNA Locus Tag | sgRNA Genes Name | ccrZ-complementation_FC | ccrZ-depletion_FC | ΔFC | adjusted p-value |
|---|---|---|---|---|---|
| SPV_0826 | holB | -2.179296063 | 0.687518638 | 2.880213051 | 1.8104E-23 |
| SPV_0827 | yabA | -0.592367822 | 1.906847812 | 2.506607828 | 1.10492E-21 |
| SPV_0878 | orfX | -0.555680867 | -2.932040114 | -2.354094105 | 1.22832E-07 |
| SPV_1694,SPV_1683,SPV_1686,SPV_1691,SPV_1689,SPV_1682,SPV_1684,SPV_1692,SPV_1688,SPV_1695,SPV_1685,SPV_1690,SPV_1693,SPV_1687 | tRNA-Thr-2,tRNA-Ile-1,tRNA-fMet-1,tRNA-Arg-3,tRNA-Met-1,tRNA-Ser-2,tRNA-Gly-1,tRNA-Leu-2,tRNA-Ile2-1,tRNA-Leu-3,tRNA-Phe-1,tRNA-Pro-1,tRNA-Gly-2,tRNA-Ser-3 | -2.893051294 | -0.90915804 | 1.991945873 | 6.42896E-06 |
| SPV_1428 | cmk | -1.974005288 | -0.275828784 | 1.714634334 | 2.03072E-05 |
| SPV_1885 | tRNA-fMet-2 | -3.439860375 | -1.494969379 | 1.954702654 | 5.46984E-05 |
| SPV_0506 | pheT | -2.482124978 | -0.7833601 | 1.708627067 | 8.04168E-05 |
| SPV_1416,SPV_1417 | murT,gatD | -1.926208965 | -0.192682941 | 1.750092894 | 8.04168E-05 |
| SPV_1477 | sepF | -3.55870129 | -1.588879631 | 1.982772842 | 0.000239202 |
| SPV_0721 | folD | -1.652401696 | -0.182804326 | 1.483308348 | 0.000467842 |
| SPV_0825 | tmk | -3.869862405 | -1.894128077 | 1.983234865 | 0.000711927 |
| SPV_2447 | srf-27 | -2.316797638 | -0.479761391 | 1.847934107 | 0.000821475 |
| SPV_0098 | unknown | -2.122984816 | -0.490913628 | 1.645051352 | 0.002533742 |
| SPV_0774 | ftsK | -0.170887755 | -1.706872257 | -1.519039798 | 0.002533742 |
| SPV_1039,SPV_1040 | ptsI,ptsH | -3.170785881 | -1.284467275 | 1.905958004 | 0.003405068 |
| SPV_1216,SPV_1217 | alaS,unknown | -2.972622339 | -1.31147978 | 1.673170925 | 0.003405068 |
| SPV_1548,SPV_1547 | gmk,rpoZ | -2.899192625 | -1.120402318 | 1.78807634 | 0.003405068 |
| SPV_1123,SPV_1127,SPV_1125,SPV_1124,SPV_1126 | licC,tarI,licA,licB,tarJ | -1.687173449 | -0.147999186 | 1.552593606 | 0.013692144 |
| SPV_1128 | tacF | -1.640755496 | -0.166329692 | 1.489081922 | 0.017904583 |
| SPV_1305,SPV_2339,SPV_1303,SPV_1304 | glyQ,unknown,unknown,glyS | -2.684730722 | -1.062662614 | 1.636808836 | 0.023216907 |
| SPV_0346,SPV_0349,SPV_0347,SPV_0348 | mvk,fni,mvaD,mvaK2 | -0.731036709 | 0.664878635 | 1.411396592 | 0.034598568 |
| SPV_1129 | licD1 | -1.252242045 | 0.179891007 | 1.446329412 | 0.034598568 |
| SPV_0967 | murA-1 | -0.746073175 | 0.703826401 | 1.465921163 | 0.039305333 |
| SPV_1475 | ylmH | -3.102326636 | -1.265744988 | 1.854601247 | 0.039305333 |
| SPV_1476 | ylmG | -3.379762828 | -1.697526932 | 1.694544623 | 0.041789451 |
| SPV_0099 | aatA | -2.239841026 | -0.618360054 | 1.633004491 | 0.051860657 |
| SPV_1484 | ddl | -5.037916775 | -3.130353866 | 1.92304724 | 0.051860657 |
| SPV_0968 | unknown | -0.408395853 | 0.946237791 | 1.365935496 | 0.066453427 |
| SPV_1373 | aspC | -2.660155248 | -1.170328881 | 1.50241862 | 0.0682772 |

| | | | | | |
|---|---|---|---|---|---|
| SPV_0248 | glmS | -2.363863897 | -0.853404628 | 1.520229101 | 0.098922549 |
| SPV_0477,SPV_0476 | trmB,unknown | -1.59224514 | -0.132900208 | 1.481771086 | 0.103831727 |
| SPV_1790 | rpmH | -2.326022076 | -0.829525691 | 1.511425831 | 0.128604337 |
| SPV_1639 | sczA | -2.113679665 | -0.536456527 | 1.595221668 | 0.132612406 |
| SPV_0526 | fba | -1.902391 | -0.537939073 | 1.376553388 | 0.147456354 |
| SPV_1791 | unknown | -3.657827241 | -1.9484279 | 1.712583712 | 0.147456354 |
| SPV_1358 | ytgP | -1.920149182 | -0.532999078 | 1.403065047 | 0.161530087 |
| SPV_1940,SPV_1941 | unknown,aspS | -2.899745251 | -1.334550345 | 1.57644954 | 0.235658723 |
| SPV_2421 | srf-23 | -2.988511895 | -1.476919037 | 1.527792322 | 0.273312395 |
| SPV_2330 | srf-20 | -4.222499881 | -2.432572316 | 1.801518882 | 0.32660583 |
| SPV_1374 | unknown | -2.78366114 | -1.383485018 | 1.415037288 | 0.401892134 |
| SPV_1886 | tRNA-Ser-4 | -3.43399682 | -1.964075773 | 1.482603185 | 0.419574733 |
| SPV_2449 | srf-28 | -1.218145989 | 0.11773098 | 1.352475915 | 0.518354988 |
| SPV_0664 | metK | -2.554868958 | -1.149335882 | 1.418056863 | 0.543429203 |
| SPV_0244,SPV_0243,SPV_0245,SPV_0246 | cdsA,uppS,eep,proS | -2.780458064 | -1.380603749 | 1.413605936 | 0.611487797 |
| SPV_2066 | tRNA-Arg-5 | -2.876064544 | -1.580095713 | 1.309153574 | 0.611487797 |
| SPV_1200,SPV_1197,SPV_1198,SPV_1199,SPV_1201 | unknown,tarQ,tarP,cps23FU,licD3 | -0.808829997 | 0.567457266 | 1.393507712 | 0.621557512 |
| SPV_0834 | pyrH | -1.844182125 | -0.478691438 | 1.379970932 | 0.65139486 |
| SPV_1041,SPV_1042,SPV_1043 | nrdH,nrdE,nrdF | -5.316688458 | -3.613743828 | 1.719680749 | 0.790311758 |
| SPV_1472 | ileS | -3.233257036 | -1.831650532 | 1.41614798 | 0.901420256 |
| SPV_0019 | rrfA | -2.829990552 | -1.442102366 | 1.402193807 | 0.951787356 |
| SPV_1628,SPV_1629 | NA,pbuX | -0.041374146 | 0.057849351 | 0.113389588 | 1 |
| SPV_0001,SPV_0002 | dnaA,dnaN | -2.878280575 | -3.498099771 | -0.605657277 | 1 |
| SPV_0003 | unknown | 0.132873911 | -0.212548713 | -0.329101517 | 1 |
| SPV_0004 | ychF | -0.215240326 | -0.0585306 | 0.17299331 | 1 |
| SPV_0005 | pth | 0.043141758 | 0.256570504 | 0.228201034 | 1 |
| SPV_0006 | mfd | 0.038646947 | 0.166016077 | 0.144240862 | 1 |
| SPV_0007,SPV_0008 | unknown,divIC | -0.46091029 | 0.151231801 | 0.625134635 | 1 |
| SPV_0010 | unknown | -0.420577558 | -0.087357953 | 0.351882237 | 1 |
| SPV_0011 | tilS | -0.226775137 | -0.086842441 | 0.150994092 | 1 |
| SPV_0012 | hpt | -0.618792135 | -0.681177474 | -0.050813567 | 1 |
| SPV_0013 | ftsH | -0.25060478 | -0.372785814 | -0.110470599 | 1 |
| SPV_0014 | comX1 | 0.121944224 | 0.131117849 | 0.023590897 | 1 |
| SPV_0016 | rrsA | -0.266224197 | -0.551918797 | -0.271751346 | 1 |
| SPV_0017 | tRNA-Ala-1 | 0.085454843 | -0.082505907 | -0.155815844 | 1 |
| SPV_0018 | rrlA | -3.048829927 | -2.044679377 | 1.015086264 | 1 |
| SPV_0025,SPV_0026 | unknown,scRNA | -0.027230796 | 0.125265836 | 0.161242246 | 1 |
| SPV_0026 | scRNA | -0.259140977 | -0.050518096 | 0.221943583 | 1 |
| SPV_0029,SPV_0027,SPV_0028 | radA,dut,unknown | -0.071742014 | 0.062546287 | 0.149106459 | 1 |
| SPV_0030,SPV_2079 | unknown,prsW | -0.254316812 | 0.764061577 | 1.03112222 | 1 |
| SPV_0033 | prs1 | -2.00358388 | -1.065804606 | 0.948639636 | 1 |

| | | | | | |
|---|---|---|---|---|---|
| SPV_0034 | unknown | 0.056589574 | 0.100219937 | 0.06248154 | 1 |
| SPV_0038 | polA | -0.059916121 | -0.220170104 | -0.150869893 | 1 |
| SPV_0039 | unknown | 0.216951857 | -0.04187237 | -0.252310374 | 1 |
| SPV_0040 | yeiH | 0.017642671 | 0.02924105 | 0.02318848 | 1 |
| SPV_0041,SPV_0045,SPV_0043,SPV_0042,SPV_0044 | araT,unknown,plsX,recO,unknown | -0.187679352 | -0.174966674 | 0.029373606 | 1 |
| SPV_0046 | blpU | 0.211960961 | -0.036174493 | -0.229289628 | 1 |
| SPV_0047 | unknown | 0.05435227 | -0.030591736 | -0.067514876 | 1 |
| SPV_0060,SPV_0056,SPV_0057,SPV_0055,SPV_0054,SPV_0062,SPV_0052,SPV_0061,SPV_0051,SPV_0059,SPV_0053,SPV_0058 | purK,vanZ,purH,purN,purM,purB,purL,unknown,purC,purE,purF,purD | -0.009727087 | 0.107498002 | 0.127736396 | 1 |
| SPV_0060,SPV_0062,SPV_0061,SPV_0059,SPV_0058 | purK,purB,unknown,purE,purD | 0.242520799 | 0.070074191 | -0.152202121 | 1 |
| SPV_0062,SPV_0061 | purB,unknown | -0.634024974 | -0.092027698 | 0.559722209 | 1 |
| SPV_0063 | strH | 0.130334965 | 0.067440373 | -0.045436706 | 1 |
| SPV_0064 | cpsR | -0.204878145 | -0.165717345 | 0.054073533 | 1 |
| SPV_0066,SPV_0065,SPV_0068,SPV_0067,SPV_0071,SPV_0069,SPV_0070 | gadV,bgaC,gadE,gadW,galM,gadF,agaS | 0.004306141 | -0.012459094 | -0.002623791 | 1 |
| SPV_0073 | unknown | -0.008876242 | -0.00312718 | 0.021764475 | 1 |
| SPV_0073,SPV_0072 | unknown,unknown | 0.072418974 | -0.074786903 | -0.133080504 | 1 |
| SPV_0074 | unknown | 0.030787653 | 0.07027947 | 0.053521531 | 1 |
| SPV_0076 | unknown | 0.012454752 | 0.087518724 | 0.085850226 | 1 |
| SPV_0077 | cabP | 0.021763215 | 0.080277447 | 0.073281186 | 1 |
| SPV_0080 | pavB | 0.026526898 | 0.0069893 | -0.000759951 | 1 |
| SPV_0082,SPV_0081 | unknown,unknown | 0.061979263 | 0.01506472 | -0.028237184 | 1 |
| SPV_0083 | rpsD | -3.364410599 | -2.394861976 | 0.985291236 | 1 |
| SPV_0086 | unknown | 0.048210386 | 0.026323123 | -0.006725104 | 1 |
| SPV_0087 | unknown | -0.004913589 | 0.114187449 | 0.130406198 | 1 |
| SPV_0088 | unknown | 0.218330896 | -0.050299275 | -0.249396652 | 1 |
| SPV_0089 | unknown | 0.211337287 | -0.017960691 | -0.212516135 | 1 |
| SPV_0090 | unknown | -0.012054384 | 0.042893871 | 0.060876852 | 1 |
| SPV_0091 | unknown | -0.095716471 | 0.050696131 | 0.159947351 | 1 |
| SPV_0092 | unknown | -0.085411157 | 0.065338316 | 0.166914523 | 1 |
| SPV_0093,SPV_0095,SPV_0094,SPV_0096 | ptvC,ptvA,ptvB,ptvR | 0.083846873 | 0.05393513 | -0.017153439 | 1 |
| SPV_0097 | unknown | 0.247115348 | 0.006758177 | -0.225388645 | 1 |
| SPV_0101,SPV_0102,SPV_0103,SPV_2095 | gph,sdhA,sdhB,unknown | 0.089143714 | 0.031497843 | -0.038048484 | 1 |
| SPV_0104 | unknown | -0.001909232 | 0.052352937 | 0.065308909 | 1 |
| SPV_0106 | unknown | -0.073504609 | 0.015740836 | 0.098037178 | 1 |
| SPV_0107,SPV_0108 | unknown,unknown | -0.107409053 | 0.104489687 | 0.219939904 | 1 |

| | | | | | |
|---|---|---|---|---|---|
| SPV_0109 | unknown | 0.111592439 | 0.197377099 | 0.097661617 | 1 |
| SPV_0110 | argG | 0.035581643 | -0.029954655 | -0.046904272 | 1 |
| SPV_0111 | argH | 0.033707184 | 0.118192589 | 0.1013849 | 1 |
| SPV_0112 | unknown | 0.36198505 | -0.019846151 | -0.367087784 | 1 |
| SPV_0113 | unknown | 0.016633063 | -0.048998881 | -0.051711722 | 1 |
| SPV_0114 | unknown | 0.016487582 | 0.066433278 | 0.066208656 | 1 |
| SPV_0115 | unknown | 0.028384346 | 0.067190128 | 0.052297117 | 1 |
| SPV_0116 | unknown | 0.101088981 | 0.036593026 | -0.050380014 | 1 |
| SPV_0117 | unknown | 0.17254511 | 0.096013266 | -0.064420512 | 1 |
| SPV_0118 | unknown | 0.17229943 | 0.046004717 | -0.110724221 | 1 |
| SPV_0123,SPV_2103,SPV_0124 | unknown,unknown,unknown | -0.007516928 | 0.030210743 | 0.04781355 | 1 |
| SPV_0126 | pspA | 0.308458854 | 0.058815189 | -0.231299876 | 1 |
| SPV_0129,SPV_0128,SPV_0127,SPV_2107 | mnmG,unknown,mnmA,srf-04 | -2.478992576 | -1.622383604 | 0.869468436 | 1 |
| SPV_0129,SPV_0128,SPV_2107 | mnmG,unknown,srf-04 | 0.05887864 | -0.072670337 | -0.1187243 | 1 |
| SPV_0130 | rnjA | -1.886997229 | -0.705232638 | 1.196230532 | 1 |
| SPV_0131 | unknown | -1.465837339 | -0.444491046 | 1.037420182 | 1 |
| SPV_0134 | unknown | -0.894469724 | -0.784371112 | 0.125158287 | 1 |
| SPV_0135 | rimI | -0.673549846 | -0.726732998 | -0.04120775 | 1 |
| SPV_0136 | tsaD | -0.804539262 | -0.763656359 | 0.055953971 | 1 |
| SPV_0138 | epsF | -0.132052559 | 0.086220802 | 0.231624039 | 1 |
| SPV_0139 | unknown | 0.002797331 | 0.068564229 | 0.077320004 | 1 |
| SPV_0140 | unknown | 0.124805646 | 0.078017957 | -0.032508998 | 1 |
| SPV_0141 | unknown | -0.152973564 | 0.100751134 | 0.267500773 | 1 |
| SPV_0142 | unknown | -0.085990182 | 0.009813859 | 0.106491479 | 1 |
| SPV_0143 | ugd | 0.098863538 | 0.107606442 | 0.023369184 | 1 |
| SPV_0144 | mutR1 | 0.368240657 | 0.008160516 | -0.345710818 | 1 |
| SPV_0147,SPV_0148 | unknown,unknown | -0.182707133 | 0.081379657 | 0.275787685 | 1 |
| SPV_0147,SPV_0148,SPV_0146 | unknown,unknown,unknown | -0.041347707 | 0.073383622 | 0.129601345 | 1 |
| SPV_0147,SPV_0148,SPV_0146,SPV_0145 | unknown,unknown,unknown,unknown | 0.510052746 | -0.075435127 | -0.565391163 | 1 |
| SPV_0149 | azlC | -1.008605082 | 0.045761183 | 1.071904471 | 1 |
| SPV_0150 | gshT | 0.238093359 | 0.011045937 | -0.213192833 | 1 |
| SPV_0151 | metQ | 0.201293303 | 0.011778383 | -0.174962443 | 1 |
| SPV_0152 | dapE | 0.209913331 | 0.142564623 | -0.054205292 | 1 |
| SPV_0154,SPV_0153 | metP,metN | 0.268255455 | -0.015079036 | -0.267295869 | 1 |
| SPV_0155 | unknown | 0.415691449 | -0.011517339 | -0.41200113 | 1 |
| SPV_0157,SPV_0156,SPV_0158 | unknown,unknown,unknown | -0.133597417 | 0.080314011 | 0.226444061 | 1 |
| SPV_0159 | unknown | 0.134951349 | 0.160796137 | 0.037515675 | 1 |
| SPV_0160 | nrdI | 0.146615464 | 0.023827142 | -0.101106642 | 1 |
| SPV_0161 | unknown | -0.089493647 | 0.016443971 | 0.122683826 | 1 |
| SPV_0163,SPV_0162 | unknown,unknown | 0.264704193 | 0.098105565 | -0.149365064 | 1 |
| SPV_0165 | hexB | -0.049354282 | 0.022016662 | 0.082223034 | 1 |

| | | | | | |
|---|---|---|---|---|---|
| SPV_0167,SPV_0168,SPV_0166,SPV_0169 | ribB,ribE,ribH,ribD | 0.114493092 | 0.018519352 | -0.081969219 | 1 |
| SPV_0171,SPV_0170 | tag,ruvA | -0.095154516 | -0.05244422 | 0.053621594 | 1 |
| SPV_0172 | unknown | -0.053884996 | -0.011980167 | 0.054056391 | 1 |
| SPV_0173 | unknown | 0.135360237 | 0.098544474 | -0.019569022 | 1 |
| SPV_0175,SPV_0174 | corA1,unknown | -1.050379528 | -0.669094135 | 0.385386445 | 1 |
| SPV_0177,SPV_0176 | ypdF,uvrA | 0.198552392 | 0.063157414 | -0.122738624 | 1 |
| SPV_0178 | spxA2 | -0.824475485 | -0.486130019 | 0.351993328 | 1 |
| SPV_0179 | unknown | 0.425920498 | -0.005568137 | -0.412484714 | 1 |
| SPV_0181,SPV_0182,SPV_0180 | yqgF,unknown,unknown | 0.185553295 | -0.536098872 | -0.706141436 | 1 |
| SPV_0183 | folC | -0.181399086 | 0.029478485 | 0.221477411 | 1 |
| SPV_0184 | unknown | -0.001904372 | 0.061524675 | 0.078729747 | 1 |
| SPV_0185 | cls | 0.175563149 | -0.237443558 | -0.39939427 | 1 |
| SPV_0186 | unknown | 0.140886032 | -0.08215101 | -0.206201397 | 1 |
| SPV_0189,SPV_0187,SPV_0190,SPV_0191,SPV_0188 | unknown,nrdD,nrdG,unknown,unknown | -0.110682909 | -0.025521892 | 0.099659846 | 1 |
| SPV_0191 | unknown | 0.24007408 | 0.105129775 | -0.120040492 | 1 |
| SPV_0210,SPV_0204,SPV_0213,SPV_0206,SPV_0203,SPV_0209,SPV_0196,SPV_0192,SPV_0211,SPV_0194,SPV_0208,SPV_0202,SPV_0200,SPV_0212,SPV_0195,SPV_0199,SPV_0193,SPV_0201,SPV_0198,SPV_0205,SPV_0207,SPV_0197 | rpsE,rplX,secY,rpsN,rplN,rplR,rplB,rpsJ,rpmD,rplD,rplF,rpsQ,rplP,rplO,rplW,rpsC,rplC,rpmC,rplV,rplE,rpsH,rpsS | -3.636222928 | -2.977788775 | 0.67463783 | 1 |
| SPV_0210,SPV_0204,SPV_0213,SPV_0206,SPV_0203,SPV_0209,SPV_0196,SPV_0211,SPV_0194,SPV_0208,SPV_0202,SPV_0200,SPV_0212,SPV_0195,SPV_0199,SPV_0193,SPV_0201,SPV_0198,SPV_0205,SPV_0207,SPV_0197 | rpsE,rplX,secY,rpsN,rplN,rplR,rplB,rpmD,rplD,rplF,rpsQ,rplP,rplO,rplW,rpsC,rplC,rpmC,rplV,rplE,rpsH,rpsS | -0.435764138 | -0.076777233 | 0.368860835 | 1 |
| SPV_0210,SPV_0204,SPV_0213,SPV_0206,SPV_0203,SPV_0209,SPV_0211,SPV_0208,SPV_0202,SPV_0212,SPV_0201,SPV_0205,SPV_0207 | rpsE,rplX,secY,rpsN,rplN,rplR,rpmD,rplF,rpsQ,rplO,rpmC,rplE,rpsH | -3.879777178 | -3.1222472 | 0.770137638 | 1 |
| SPV_0214 | adk | -2.911565835 | -1.769488408 | 1.163528879 | 1 |
| SPV_0218,SPV_0217,SPV_0216,SPV_0219,SPV_0215,SPV_2122 | rpoA,rpsK,rpsM,rplQ,infA,rpmJ | -3.206072971 | -2.557198398 | 0.667821784 | 1 |
| SPV_0220,SPV_0221 | unknown,unknown | -0.074738324 | 0.092674832 | 0.184420825 | 1 |
| SPV_0222 | gpmB1 | -0.009257085 | 0.084740781 | 0.100609496 | 1 |
| SPV_0225 | unknown | 0.207374709 | 0.019884442 | -0.180049722 | 1 |
| SPV_0229,SPV_0228 | unknown,unknown | 0.123847827 | -0.050109261 | -0.154438725 | 1 |
| SPV_0230 | deoR | 0.061949399 | 0.046630421 | 0.001673957 | 1 |

| | | | | | |
|---|---|---|---|---|---|
| SPV_0231 | unknown | -0.026402807 | 0.057357873 | 0.095009501 | 1 |
| SPV_0232 | unknown | 0.185628354 | 0.013989267 | -0.152632117 | 1 |
| SPV_0233 | unknown | -0.051752525 | 0.18960221 | 0.250431458 | 1 |
| SPV_0234 | unknown | -0.196355923 | 0.060004709 | 0.268823555 | 1 |
| SPV_0235 | unknown | 0.032730778 | 0.052658436 | 0.033992646 | 1 |
| SPV_0236 | talC | 0.126638356 | 0.003350314 | -0.109873091 | 1 |
| SPV_0237 | gldA | 0.032693575 | 0.000156336 | -0.023834833 | 1 |
| SPV_0238 | leuS | -2.813203571 | -1.88731115 | 0.939296199 | 1 |
| SPV_0239 | unknown | 0.037605749 | 0.07136472 | 0.047969847 | 1 |
| SPV_0240 | unknown | 0.253631558 | 0.030377848 | -0.205777273 | 1 |
| SPV_0241 | ruvB | -0.046207649 | -0.323020129 | -0.25514593 | 1 |
| SPV_0242 | unknown | -2.576895942 | -1.345957108 | 1.242104632 | 1 |
| SPV_0247 | bglA | 0.286798294 | 0.037021966 | -0.227479154 | 1 |
| SPV_0249 | unknown | 0.26918216 | 0.090055884 | -0.160758287 | 1 |
| SPV_0250 | spuA | 0.289147259 | 0.151023475 | -0.117460879 | 1 |
| SPV_0251,SPV_0252 | rpsL,rpsG | -3.96069808 | -2.854782942 | 1.119889227 | 1 |
| SPV_0253 | fusA | -2.733317564 | -2.443124593 | 0.304052493 | 1 |
| SPV_0254 | polC | -1.899465134 | -1.118467018 | 0.785929627 | 1 |
| SPV_0255 | relB1 | 0.162829837 | 0.026360496 | -0.123962646 | 1 |
| SPV_0256 | relE1 | -0.230478023 | -0.058233632 | 0.185475975 | 1 |
| SPV_0257 | unknown | -0.005512892 | 0.13954463 | 0.15883714 | 1 |
| SPV_0258 | pepS | 0.203817389 | 0.139671294 | -0.050378829 | 1 |
| SPV_0259 | unknown | 0.090621948 | 0.041299906 | -0.038319113 | 1 |
| SPV_0260 | rsuA-1 | 0.227431195 | 0.075101908 | -0.136653823 | 1 |
| SPV_0261 | pepC | 0.079481405 | -0.022263948 | -0.086593802 | 1 |
| SPV_0262,SPV_0263,SPV_0264 | manN,manM,manL | -0.068207616 | 0.116615669 | 0.198128779 | 1 |
| SPV_0265 | adhA | -0.038926464 | 0.034496756 | 0.08308434 | 1 |
| SPV_0266 | unknown | -0.033187986 | 0.041807492 | 0.089336677 | 1 |
| SPV_0267 | unknown | 0.224126196 | 0.022046709 | -0.188461176 | 1 |
| SPV_0268 | unknown | 0.091557986 | -0.089374314 | -0.162011434 | 1 |
| SPV_0269 | sulA | -1.285870938 | -0.335112323 | 0.956838382 | 1 |
| SPV_0270 | sulB | -1.102725429 | -0.223104099 | 0.893889524 | 1 |
| SPV_0271 | sulC | -1.210052448 | -0.11633491 | 1.105624595 | 1 |
| SPV_0272 | sulD | -0.162952073 | 0.074557984 | 0.259383726 | 1 |
| SPV_0273 | unknown | 0.104867635 | 0.047623927 | -0.04868996 | 1 |
| SPV_0274,SPV_0275 | rplM,rpsI | -3.386182267 | -2.590386935 | 0.811575596 | 1 |
| SPV_0276 | unknown | 0.137391415 | 0.185520292 | 0.061963368 | 1 |
| SPV_0283,SPV_0280,SPV_0278,SPV_0277,SPV_0281,SPV_0279,SPV_0282 | celD,celR,unknown,celA,celC,celB,unknown | 0.037714502 | 0.186293938 | 0.163859601 | 1 |
| SPV_0283,SPV_0280,SPV_0281,SPV_0279,SPV_0282 | celD,celR,celC,celB,unknown | 0.076657097 | 0.03226849 | -0.025974618 | 1 |

| | | | | | |
|---|---|---|---|---|---|
| SPV_0283,SPV_0280,SPV_0281,SPV_0282 | celD,celR,celC,unknown | 0.034691479 | 0.116114331 | 0.092133639 | 1 |
| SPV_0284 | unknown | 0.149220111 | 0.06032251 | -0.082003528 | 1 |
| SPV_0285 | unknown | 0.087206243 | 0.118209973 | 0.043262037 | 1 |
| SPV_0286 | basA | 0.059819077 | 0.103076538 | 0.060135044 | 1 |
| SPV_0287 | spnHL | 0.138703187 | 0.036165368 | -0.086993604 | 1 |
| SPV_0289,SPV_0290,SPV_0291,SPV_0292 | eda,unknown,unknown,unknown | -0.049226031 | 0.053542516 | 0.115027109 | 1 |
| SPV_0293 | unknown | 0.208019844 | -0.090084745 | -0.281119839 | 1 |
| SPV_0294 | ugl | 0.002776453 | 0.140410185 | 0.150151154 | 1 |
| SPV_0295 | unknown | 0.201583626 | 0.008469125 | -0.172831873 | 1 |
| SPV_0296 | unknown | 0.146137242 | 0.112065908 | -0.023001753 | 1 |
| SPV_0297 | unknown | -0.104463856 | 0.111261041 | 0.226529584 | 1 |
| SPV_0298 | yajC-1 | 0.123482095 | 0.053158251 | -0.050291429 | 1 |
| SPV_0301 | regR | 0.026298122 | 0.014875677 | 0.004859472 | 1 |
| SPV_0302 | unknown | 0.185538142 | -0.04205335 | -0.214111824 | 1 |
| SPV_0303 | unknown | 0.225145168 | 0.01381164 | -0.197212804 | 1 |
| SPV_0304 | mraW | -6.582518444 | -4.965905379 | 1.598609433 | 1 |
| SPV_0305 | ftsL | -6.377251211 | -4.986828619 | 1.403338857 | 1 |
| SPV_0307,SPV_0306 | mraY,pbp2x | -5.550050933 | -4.603868331 | 0.968890279 | 1 |
| SPV_0308 | clpL | 0.143291343 | 0.023804903 | -0.102311835 | 1 |
| SPV_0309 | luxS | 0.004394956 | -0.060025999 | -0.047812367 | 1 |
| SPV_0310 | unknown | -0.035502779 | 0.068922512 | 0.120063981 | 1 |
| SPV_0311,SPV_0313 | dexB,unknown | -0.09098779 | 0.066232161 | 0.170483808 | 1 |
| SPV_0326,SPV_0329,SPV_0330,SPV_0323,SPV_0320,SPV_0317,SPV_0315,SPV_0319,SPV_0324,SPV_0328,SPV_0321,SPV_0322,SPV_0316,SPV_0327,SPV_0318,SPV_0325 | cps2K,cps2M,cps2N,cps2H,cps2T,cps2C,cps2A,cps2E,cps2I,cps2L,cps2F,cps2G,cps2B,cps2P,cps2D,cps2J | 0.346439847 | 0.576109055 | 0.236663181 | 1 |
| SPV_0326,SPV_0329,SPV_0330,SPV_0323,SPV_0324,SPV_0328,SPV_0321,SPV_0322,SPV_0327,SPV_0325 | cps2K,cps2M,cps2N,cps2H,cps2I,cps2L,cps2F,cps2G,cps2P,cps2J | 0.373099686 | 0.364272803 | 0.003617385 | 1 |
| SPV_0331 | cps2O | -0.532618107 | -0.625175864 | -0.076756416 | 1 |
| SPV_0334 | aliA | 0.071274589 | 0.073561916 | 0.016492441 | 1 |
| SPV_0335 | eng | 0.120401081 | -0.051580023 | -0.153391362 | 1 |
| SPV_0337,SPV_0336 | recU,pbp1a | -0.496868373 | -0.611200952 | -0.103420387 | 1 |
| SPV_0338 | unknown | -0.009208175 | 0.034620663 | 0.059143001 | 1 |
| SPV_0339 | gpsB | -1.731578747 | -0.502025886 | 1.255082341 | 1 |
| SPV_0340 | rnpB | -2.059095939 | -0.94412287 | 1.127231272 | 1 |
| SPV_0341 | rlmL | -0.005870506 | -0.206447011 | -0.186679034 | 1 |
| SPV_0342 | mapZ | -0.086476781 | -0.259951927 | -0.159976621 | 1 |
| SPV_0344,SPV_0343 | ritR,gnd | -4.074283404 | -2.532802677 | 1.558384402 | 1 |
| SPV_0345 | cbpF | 0.275663763 | 0.050641994 | -0.209437124 | 1 |

| | | | | | |
|---|---|---|---|---|---|
| SPV_0352,SPV_2148,SPV_0350,SPV_0351 | vraR,alkD,vraT,vraS | 0.096890224 | -0.121112538 | -0.205892616 | 1 |
| SPV_0355,SPV_0357,SPV_0356 | unknown,cbpK,cbpG | 0.016643871 | 0.097157781 | 0.09650284 | 1 |
| SPV_0358 | unknown | 0.296875926 | -0.087508681 | -0.371338185 | 1 |
| SPV_0360 | mtlA | -0.027629668 | -0.004234204 | 0.037832689 | 1 |
| SPV_0361 | mtlR | -0.106009436 | -0.023586578 | 0.094555105 | 1 |
| SPV_0362 | mtlA2 | 0.306389833 | 0.010254632 | -0.28032221 | 1 |
| SPV_0363 | mtlD | 0.207082425 | 0.146819136 | -0.044846248 | 1 |
| SPV_0364 | unknown | 0.214332294 | 0.108659667 | -0.091758947 | 1 |
| SPV_0365 | tig | -0.077101697 | -0.54364709 | -0.45266935 | 1 |
| SPV_0366 | yrrC | 0.111794667 | -0.165866329 | -0.26078253 | 1 |
| SPV_0367,SPV_0368 | lepB,rnhC | -1.434119389 | -0.786291864 | 0.661729492 | 1 |
| SPV_0370,SPV_0369 | zapB,zapA | 0.066973321 | -0.093114779 | -0.144471592 | 1 |
| SPV_0371 | mutS2 | 0.076822419 | 0.009280933 | -0.055015712 | 1 |
| SPV_0372 | glyP | -0.179124368 | -0.222177247 | -0.025801053 | 1 |
| SPV_0373 | mip | 0.215786337 | -0.046159941 | -0.242550793 | 1 |
| SPV_0374 | shetA | 0.289842276 | 0.138051364 | -0.140357668 | 1 |
| SPV_0375 | serS | -2.357551145 | -1.121593427 | 1.254765228 | 1 |
| SPV_0376,SPV_0375,SPV_0377 | manO,serS,lysC | -0.169511908 | 0.183613398 | 0.365605939 | 1 |
| SPV_0380,SPV_0378,SPV_0379,SPV_0381 | fabH,fabM,fabT,acpP | 0.136695567 | 0.070096467 | -0.051591431 | 1 |
| SPV_0380,SPV_0379,SPV_0381 | fabH,fabT,acpP | -1.132980415 | 0.070424581 | 1.214332814 | 1 |
| SPV_0385,SPV_0389,SPV_0388,SPV_0386,SPV_0383,SPV_0390,SPV_0384,SPV_0387,SPV_0382 | fabF,accD,accC,accB,fabD,accA,fabG1,fabZ,fabK | -0.773972811 | -0.015694392 | 0.770056135 | 1 |
| SPV_0393,SPV_0395,SPV_0394 | nusB,efp,unknown | -0.59633339 | -0.641542508 | -0.031923665 | 1 |
| SPV_0397,SPV_0398,SPV_0396 | gatA,gatC,gatB | -2.825092559 | -1.634061529 | 1.208989526 | 1 |
| SPV_0399 | prfC | 0.144447148 | -0.105176485 | -0.239230008 | 1 |
| SPV_0400 | unknown | -0.018604455 | 0.020020224 | 0.049374625 | 1 |
| SPV_0401 | rpmB | -0.067757174 | -0.124000361 | -0.04492795 | 1 |
| SPV_0402 | asp23 | -0.169882533 | 0.051829151 | 0.239424294 | 1 |
| SPV_0403 | unknown | -0.724191284 | 0.02992853 | 0.767771373 | 1 |
| SPV_0404,SPV_0406,SPV_2159,SPV_0405,SPV_0408 | ilvB,ilvC,unknown,ilvH,unknown | 0.257476931 | 0.052571788 | -0.193429803 | 1 |
| SPV_0409 | ilvA | -0.04538977 | 0.079360582 | 0.136850952 | 1 |
| SPV_0410 | unknown | 0.06594488 | -0.520940221 | -0.571235038 | 1 |
| SPV_0412,SPV_0411 | glnHP1,glnQ1 | 0.282423114 | 0.171929864 | -0.095298942 | 1 |
| SPV_0413 | unknown | 0.091096509 | 0.136021309 | 0.059525247 | 1 |
| SPV_0414 | unknown | -0.055485579 | 0.009000953 | 0.078485683 | 1 |
| SPV_0417 | uppP | -0.081277931 | 0.431738628 | 0.527864413 | 1 |
| SPV_0418 | unknown | -0.129966935 | 0.157736295 | 0.297741883 | 1 |
| SPV_0419 | dinP | 0.057883393 | -0.046456776 | -0.089516371 | 1 |
| SPV_0420 | pfl | 0.457744766 | 0.079444681 | -0.373246957 | 1 |

| | | | | | |
|---|---|---|---|---|---|
| SPV_0422 | unknown | 0.26254001 | -0.102238732 | -0.348897274 | 1 |
| SPV_0423 | xylR | 0.23092213 | 0.154768571 | -0.058999198 | 1 |
| SPV_0424 | unknown | 0.103106981 | -0.052179805 | -0.142789619 | 1 |
| SPV_0425 | unknown | 0.111612836 | 0.098510137 | 0.010083896 | 1 |
| SPV_0426 | lacF-1 | -0.154601256 | 0.15098528 | 0.318429209 | 1 |
| SPV_0427 | lacG-1 | 0.022269013 | 0.064706877 | 0.054444602 | 1 |
| SPV_0428 | lacE-1 | 0.027280508 | 0.1150082 | 0.098651459 | 1 |
| SPV_0429,SPV_0430 | trkH,trkA | 0.586635605 | -0.138244854 | -0.711857758 | 1 |
| SPV_0431 | unknown | 0.150540785 | 0.057680225 | -0.081204616 | 1 |
| SPV_0432 | mtsA | 0.113312168 | 0.014800088 | -0.086122512 | 1 |
| SPV_0433 | unknown | -0.059472588 | 0.296414692 | 0.369733265 | 1 |
| SPV_0434 | mtsB | 0.149507092 | 0.072747357 | -0.058233569 | 1 |
| SPV_0435 | mtsC | 0.142550367 | 0.036910792 | -0.088556376 | 1 |
| SPV_0436 | cspR | 0.154457053 | 0.067800865 | -0.076217698 | 1 |
| SPV_0438 | unknown | -0.10838623 | -0.010145783 | 0.116065297 | 1 |
| SPV_0439 | unknown | -0.02983907 | -0.131671501 | -0.092790304 | 1 |
| SPV_0440 | unknown | -0.029329385 | -0.011774788 | 0.034337031 | 1 |
| SPV_0441 | rpoE | -1.8329069 | -0.997760684 | 0.850369039 | 1 |
| SPV_0442 | pyrG | 0.340062415 | -0.013869884 | -0.338411065 | 1 |
| SPV_0443 | nptA | 0.022835407 | -0.408336636 | -0.420172225 | 1 |
| SPV_0444 | endoD | 0.153200963 | 0.13504003 | -0.005675806 | 1 |
| SPV_0445 | pgk | -0.812636952 | -0.383984432 | 0.444987062 | 1 |
| SPV_0446 | unknown | 0.054720442 | 0.242963901 | 0.205100281 | 1 |
| SPV_0447,SPV_0448 | glnR,glnA | 0.007039529 | 0.362231456 | 0.36883811 | 1 |
| SPV_0449 | unknown | 0.141226502 | 0.073225014 | -0.054001789 | 1 |
| SPV_0452 | creX | -0.027676159 | -0.07399907 | -0.034076152 | 1 |
| SPV_0453 | unknown | 0.065027224 | 0.083648132 | 0.033693005 | 1 |
| SPV_0455,SPV_0454,SPV_2170 | hsdR,hsdM,hsdS-F | -0.088734105 | 0.059435406 | 0.164937985 | 1 |
| SPV_0456 | unknown | 0.063064106 | -0.07846598 | -0.12930862 | 1 |
| SPV_0457 | unknown | 0.046323205 | 0.07273979 | 0.043002346 | 1 |
| SPV_0458,SPV_0460,SPV_0459,SPV_0461,SPV_2171 | hrcA,dnaK,grpE,dnaJ,unknown | -0.115395134 | -0.21045909 | -0.078907399 | 1 |
| SPV_0463,SPV_0462 | unknown,unknown | 0.139355519 | 0.036536905 | -0.092532724 | 1 |
| SPV_0464,SPV_0465 | ecsA,ecsB | 0.100407917 | 0.052413734 | -0.031432304 | 1 |
| SPV_0466 | blpT | -0.02232523 | 0.116176498 | 0.159403678 | 1 |
| SPV_0467,SPV_0468,SPV_0469 | blpS,blpR,blpH | -0.048458519 | 0.011956748 | 0.068028467 | 1 |
| SPV_0470,SPV_0471,SPV_0472 | blpC,blpB,blpA | 0.216829476 | 0.011378397 | -0.192022416 | 1 |
| SPV_0475,SPV_2176,SPV_0474,SPV_0473 | pncP,pncW,blpZ,blpY | 0.014098972 | 0.113270718 | 0.115241716 | 1 |
| SPV_0482,SPV_0480,SPV_0478,SPV_0483,SPV_0481,SPV_0479 | infB,unknown,rimP,rbfA,unknown,nusA | -1.272750296 | -0.163809821 | 1.123715519 | 1 |
| SPV_0484 | unknown | 0.104917285 | 0.028400282 | -0.057738101 | 1 |
| SPV_0485 | unknown | -0.075015174 | 0.08954703 | 0.179285503 | 1 |

| | | | | | |
|---|---|---|---|---|---|
| SPV_0486 | unknown | -0.021674694 | -0.02499013 | 0.01411212 | 1 |
| SPV_0487,SPV_0488 | unknown,unknown | 0.004282334 | -0.032281373 | -0.020335368 | 1 |
| SPV_0489 | unknown | -0.149774724 | 0.126189633 | 0.290765904 | 1 |
| SPV_0490 | unknown | -0.035217935 | -0.091349108 | -0.044694937 | 1 |
| SPV_0493 | unknown | -2.59524607 | -1.144653895 | 1.467525113 | 1 |
| SPV_0494 | valS | -2.363018164 | -1.107724882 | 1.266570861 | 1 |
| SPV_0496,SPV_0495 | unknown,unknown | -2.558888227 | -2.124303823 | 0.451656882 | 1 |
| SPV_0501 | bglG | 0.005491468 | 0.106821243 | 0.11507378 | 1 |
| SPV_0502 | bglF | 0.180661284 | 0.049002386 | -0.119328628 | 1 |
| SPV_0503 | bglA-2 | 0.04342642 | 0.093641568 | 0.058056059 | 1 |
| SPV_0504,SPV_0505 | pheS,paiA | -1.545812699 | -0.722428619 | 0.834761639 | 1 |
| SPV_0507 | unknown | 0.028838096 | 0.105508373 | 0.087133519 | 1 |
| SPV_0508 | unknown | -0.189128108 | 0.088868391 | 0.289551528 | 1 |
| SPV_0509 | higA | 0.051666061 | 0.042350247 | 0.009229306 | 1 |
| SPV_0510,SPV_0511 | metE,metF | 0.114073901 | 0.005438456 | -0.096847494 | 1 |
| SPV_0512,SPV_0516,SPV_0513,SPV_0517,SPV_0514,SPV_0515 | pnp,mrnC,cysE,unknown,unknown,cysS | -1.755661309 | -0.775009105 | 0.991553229 | 1 |
| SPV_0523,SPV_0522,SPV_0521 | vex3,vex2,vex1 | 0.04215823 | 0.08388123 | 0.057045787 | 1 |
| SPV_0525,SPV_0524 | vncS,vncR | 0.049192436 | -0.05010129 | -0.078489908 | 1 |
| SPV_0527 | unknown | 0.229164921 | 0.03901667 | -0.170905215 | 1 |
| SPV_0531,SPV_0529,SPV_0528,SPV_0530 | glnQ2,glnP2a,glnP2b,glnH2 | 0.072565048 | 0.087524414 | 0.026768349 | 1 |
| SPV_0532 | recJ | 0.08371482 | -0.013614818 | -0.085646708 | 1 |
| SPV_0533 | rnjB | -1.587509128 | -0.370818682 | 1.229987353 | 1 |
| SPV_0534 | estA | 0.005608188 | 0.140642805 | 0.146727709 | 1 |
| SPV_0535,SPV_0536 | murM,murN | 0.114697694 | 0.37267894 | 0.27267531 | 1 |
| SPV_0537 | unknown | 0.149496807 | 0.087796342 | -0.046063262 | 1 |
| SPV_0539,SPV_0538 | unknown,uvrC | 0.035022695 | 0.001323434 | -0.019959581 | 1 |
| SPV_0540 | unknown | 0.211668927 | 0.015557428 | -0.185510593 | 1 |
| SPV_0542,SPV_0541 | pepV,nrd | -0.211494399 | -0.211465755 | 0.01532245 | 1 |
| SPV_0543 | unknown | -0.078200678 | -0.017258848 | 0.074316693 | 1 |
| SPV_0546 | brnQ | 0.054771267 | 0.107098811 | 0.066516006 | 1 |
| SPV_0548,SPV_0547,SPV_0549 | unknown,unknown,ldcB | 0.034692768 | 0.144250626 | 0.122209539 | 1 |
| SPV_0550,SPV_0551 | rplK,rplA | -2.343471796 | -1.189260785 | 1.171600999 | 1 |
| SPV_0552 | unknown | 0.034745925 | 0.0745809 | 0.052524703 | 1 |
| SPV_0553 | unknown | -0.020511888 | 0.034858298 | 0.06677444 | 1 |
| SPV_0554 | unknown | 0.136173679 | 0.045121275 | -0.077179173 | 1 |
| SPV_0555 | unknown | 0.283485519 | 0.104078492 | -0.164793322 | 1 |
| SPV_0556 | unknown | 0.096841037 | 0.084249577 | 0.001475728 | 1 |
| SPV_0557 | unknown | 0.054334411 | 0.1496823 | 0.10862122 | 1 |
| SPV_0558 | prtA | 0.465554781 | 0.475225687 | 0.027811689 | 1 |
| SPV_0561,SPV_0560,SPV_0562,SPV_0559 | unknown,unknown,bgaA,unknown | 0.055371045 | 0.07374473 | 0.035300062 | 1 |

| | | | | | |
|---|---|---|---|---|---|
| SPV_0563 | unknown | 0.026003178 | 0.108785788 | 0.093993654 | 1 |
| SPV_0564 | unknown | -0.022622645 | 0.281664197 | 0.317751818 | 1 |
| SPV_0567 | ytqB | -0.003342845 | 0.109748951 | 0.127581764 | 1 |
| SPV_0568 | unknown | 0.069386252 | 0.155783942 | 0.100557741 | 1 |
| SPV_0569 | nha2 | 0.165646258 | 0.057542365 | -0.09108974 | 1 |
| SPV_0570 | yihY | -0.004149693 | 0.059085879 | 0.072395409 | 1 |
| SPV_0573,SPV_0572,SPV_0571 | msrAB2,unknown,ccdA-1 | -0.004271706 | -0.00570734 | 0.015965548 | 1 |
| SPV_0575,SPV_0574 | yesM,yesN | 0.056531505 | 0.042473723 | -0.001800901 | 1 |
| SPV_0576 | ywlG | 0.131927353 | 0.031141294 | -0.086672501 | 1 |
| SPV_0577 | zmpB | 0.087000331 | 0.013741955 | -0.053937313 | 1 |
| SPV_0578 | pabB | 0.006723289 | 0.043907906 | 0.051567089 | 1 |
| SPV_0579 | cbpL | 0.081136549 | 0.05153526 | -0.015046307 | 1 |
| SPV_0580 | gki | -0.163472053 | 0.047784457 | 0.225246739 | 1 |
| SPV_0581 | thyA | -1.105967671 | -0.335620807 | 0.787186738 | 1 |
| SPV_0582 | unknown | 0.081225428 | 0.016578272 | -0.062436423 | 1 |
| SPV_0585,SPV_0584,SPV_0586,SPV_0583,SPV_0587 | unknown,hflX,rnz,miaA,unknown | -0.308673303 | -0.776790416 | -0.45338729 | 1 |
| SPV_0588 | mtaR | 0.04280643 | 0.119216803 | 0.093430705 | 1 |
| SPV_0590,SPV_0589 | unknown,unknown | -0.093074733 | 0.125284346 | 0.230737787 | 1 |
| SPV_0591 | unknown | -0.077858627 | 0.137950032 | 0.228388811 | 1 |
| SPV_0592 | rsuA-2 | 0.344009448 | 0.038187876 | -0.289530406 | 1 |
| SPV_0594,SPV_0593 | unknown,typA | 0.092212578 | -0.284329247 | -0.361546062 | 1 |
| SPV_0597,SPV_0596,SPV_0595 | unknown,unknown,unknown | 0.031278064 | 0.019682024 | 0.003043425 | 1 |
| SPV_0599,SPV_0598 | murG,murD | -5.395518522 | -4.346845621 | 1.067009218 | 1 |
| SPV_0600 | divIB | -0.213894914 | 0.558126864 | 0.788858947 | 1 |
| SPV_0601 | unknown | -0.096527067 | 0.292958571 | 0.39997608 | 1 |
| SPV_0602 | unknown | 0.074681905 | 0.361312504 | 0.303595939 | 1 |
| SPV_0605 | unknown | 0.11141904 | -0.014861169 | -0.117968036 | 1 |
| SPV_0606 | unknown | -0.067221768 | 0.146649704 | 0.22569863 | 1 |
| SPV_0607 | unknown | 0.180931846 | 0.107682416 | -0.058175159 | 1 |
| SPV_0608,SPV_0610,SPV_0609 | pyrF,unknown,pyrE | -0.087802384 | -0.02166813 | 0.080922248 | 1 |
| SPV_0610 | unknown | 0.335987319 | 0.06150168 | -0.255329746 | 1 |
| SPV_0611 | unknown | 0.075791309 | 0.185665158 | 0.125366122 | 1 |
| SPV_0612 | unknown | 0.105792683 | 0.125331975 | 0.031505008 | 1 |
| SPV_0613 | unknown | -0.118267357 | -0.12781198 | -0.001507715 | 1 |
| SPV_0614 | unknown | 0.098222181 | -0.025760772 | -0.105346173 | 1 |
| SPV_0616,SPV_0618,SPV_0615,SPV_0617 | glnQ3,glnP3a,glnH3,glnP3b | 0.086838994 | -0.021911133 | -0.092638206 | 1 |
| SPV_0619 | unknown | -0.096854824 | 0.093625432 | 0.199878838 | 1 |
| SPV_0620 | lysS | -2.052865288 | -1.06378095 | 1.004368831 | 1 |
| SPV_0621 | lctO | 0.04106681 | -0.073974839 | -0.10283887 | 1 |
| SPV_0623 | thiM-1 | -0.039166986 | 0.112658546 | 0.168895821 | 1 |
| SPV_0624 | thiE-1 | -0.152522011 | 0.115096988 | 0.280188828 | 1 |

| | | | | | |
|---|---|---|---|---|---|
| SPV_0632 | thiD | 0.099869933 | 0.116219213 | 0.029331507 | 1 |
| SPV_0634,SPV_0635,SPV_0633 | cupA,ctpA,copY | 0.177044405 | -0.007587528 | -0.170899971 | 1 |
| SPV_0636 | spxB | 0.137250519 | -0.107832179 | -0.2308168 | 1 |
| SPV_0637 | unknown | -0.068134559 | 0.114248129 | 0.193709016 | 1 |
| SPV_0640 | unknown | 0.142220304 | -0.015851308 | -0.142033279 | 1 |
| SPV_0641 | manA | -0.12257343 | 0.041156151 | 0.180196973 | 1 |
| SPV_0642 | unknown | 0.109116166 | -0.015921983 | -0.112466687 | 1 |
| SPV_0643 | unknown | -0.039281051 | 0.089811624 | 0.141587432 | 1 |
| SPV_0644,SPV_0645 | unknown,unknown | 0.205316797 | 0.123720912 | -0.059818481 | 1 |
| SPV_0646 | unknown | 0.154496789 | 0.102227345 | -0.036116969 | 1 |
| SPV_0647,SPV_0648 | unknown,comEB | 0.063136566 | 0.085243228 | 0.033798631 | 1 |
| SPV_0649 | upp | -0.740024236 | -0.658724888 | 0.095434397 | 1 |
| SPV_0650 | clpP | -0.182927635 | -0.902309506 | -0.703817382 | 1 |
| SPV_0651 | unknown | 0.354509417 | 0.141631141 | -0.192380545 | 1 |
| SPV_0652 | livJ | 0.292251499 | 0.065818378 | -0.209320492 | 1 |
| SPV_0653 | livH | 0.114166928 | 0.05342475 | -0.041171846 | 1 |
| SPV_0654 | livM | -0.074895908 | 0.060785587 | 0.157924261 | 1 |
| SPV_0655 | livG | 0.063566949 | 0.061757027 | 0.014134033 | 1 |
| SPV_0656 | livF | 0.287333741 | 0.06701357 | -0.199880713 | 1 |
| SPV_0657 | unknown | -0.112021093 | -0.030237021 | 0.094113894 | 1 |
| SPV_0659,SPV_0660,SPV_2217 | ftsE,ftsX,NA | -1.76081467 | -1.887193255 | -0.10769489 | 1 |
| SPV_0661 | malT | 0.111752437 | 0.031327378 | -0.064204202 | 1 |
| SPV_0662 | unknown | 0.089290019 | 0.100758906 | 0.02405966 | 1 |
| SPV_0663 | yqfR | -0.91709097 | -0.457392612 | 0.471402465 | 1 |
| SPV_0666,SPV_0665 | holA,pyrDa | 0.124479163 | -0.004898628 | -0.114748024 | 1 |
| SPV_0667 | sodA | 0.062614959 | -0.110729861 | -0.150965693 | 1 |
| SPV_0668 | unknown | -0.051243603 | 0.139162673 | 0.201431986 | 1 |
| SPV_0669 | rlmN | 0.205919558 | 0.026294554 | -0.165655095 | 1 |
| SPV_0670 | unknown | 0.190434334 | 0.012564734 | -0.158482004 | 1 |
| SPV_0671 | unknown | 0.032628868 | 0.079409852 | 0.060322139 | 1 |
| SPV_0672 | ppiA | 0.103826635 | 0.134047802 | 0.041163258 | 1 |
| SPV_0674,SPV_0675 | rpsP,khpA | -3.244414842 | -2.82587381 | 0.433152669 | 1 |
| SPV_0678 | rimM | -0.130563783 | 0.073385722 | 0.219740754 | 1 |
| SPV_0679 | trmD | -0.115603457 | 0.06344484 | 0.192645949 | 1 |
| SPV_0680 | unknown | 0.34877409 | 0.242872913 | -0.090370197 | 1 |
| SPV_0681 | unknown | -0.089393486 | -0.096875416 | 0.005106698 | 1 |
| SPV_0682 | unknown | 0.208068618 | 0.06664984 | -0.125055687 | 1 |
| SPV_0683 | unknown | 0.265182503 | 0.070578592 | -0.182804869 | 1 |
| SPV_0684 | bioY | 0.189983802 | -0.06564269 | -0.239071365 | 1 |
| SPV_0685 | gor | 0.047735636 | -0.007190105 | -0.04165384 | 1 |
| SPV_0686,SPV_0687,SPV_0688 | unknown,unknown,unknown | 0.285817604 | 0.08643326 | -0.187098387 | 1 |
| SPV_0689 | metG | -2.664479743 | -1.433624453 | 1.247920599 | 1 |
| SPV_0690 | unknown | 0.233039624 | 0.068506743 | -0.148626177 | 1 |

| | | | | | |
|---|---|---|---|---|---|
| SPV_0691 | unknown | -0.019456721 | -0.067025852 | -0.032373849 | 1 |
| SPV_0692 | unknown | 0.364761836 | 0.036327856 | -0.312200659 | 1 |
| SPV_0693 | unknown | -0.131411341 | 0.111585024 | 0.252419323 | 1 |
| SPV_0695,SPV_0694 | fabG2,unknown | -0.029127514 | 0.030301121 | 0.073929791 | 1 |
| SPV_0696 | unknown | 0.151329027 | -0.004767887 | -0.139145018 | 1 |
| SPV_0697 | unknown | 0.058869997 | -0.118538912 | -0.163046715 | 1 |
| SPV_0698 | unknown | 0.032540826 | 0.028575771 | 0.007235529 | 1 |
| SPV_0699 | unknown | -0.085437226 | -0.058883568 | 0.039139531 | 1 |
| SPV_0700 | pepN | -0.204058024 | -0.058332968 | 0.163359443 | 1 |
| SPV_0701,SPV_0702 | ciaR,ciaH | 0.02323254 | 0.138848494 | 0.125480769 | 1 |
| SPV_0703 | unknown | -0.101936143 | 0.383518248 | 0.495208895 | 1 |
| SPV_0704 | unknown | 0.099222185 | 0.043766661 | -0.040501596 | 1 |
| SPV_0705 | yoaA | 0.157210979 | 0.111923057 | -0.030188657 | 1 |
| SPV_0706 | rodA | 0.120785446 | 0.033665577 | -0.074804704 | 1 |
| SPV_0707 | thiJ | 0.189517337 | 0.071918743 | -0.09984804 | 1 |
| SPV_0708 | unknown | -2.879995805 | -1.501628247 | 1.387378429 | 1 |
| SPV_0709 | gyrB | -0.314221339 | -0.109794024 | 0.219043538 | 1 |
| SPV_0710 | ezrA | -1.835249007 | -1.841762167 | 0.006865766 | 1 |
| SPV_0713 | unknown | -0.426020503 | 0.625945529 | 1.065877355 | 1 |
| SPV_0714 | unknown | 0.161132504 | 0.153510416 | 0.006293807 | 1 |
| SPV_0715 | unknown | 0.122565301 | 0.083119912 | -0.024852716 | 1 |
| SPV_0717 | clpE | 0.172114347 | 0.071947232 | -0.085445496 | 1 |
| SPV_0718 | unknown | 0.046262886 | -0.333126661 | -0.367633944 | 1 |
| SPV_0721,SPV_0720,SPV_0719 | folD,glnQ4,glnP4 | -0.328766554 | 0.084868963 | 0.427601352 | 1 |
| SPV_0722 | nnrD | -0.058896246 | 0.066941107 | 0.139100793 | 1 |
| SPV_0723,SPV_0725,SPV_0726,SPV_072<br>4 | rpiA,unknown,unknown,deoB | 0.011751863 | -0.102041293 | -0.100586823 | 1 |
| SPV_0729 | unknown | 0.108966339 | 0.079687426 | -0.015555327 | 1 |
| SPV_0730 | deoD | -0.041787909 | 0.066938935 | 0.124514057 | 1 |
| SPV_0731 | flaR | 0.225050099 | 0.045079143 | -0.169387137 | 1 |
| SPV_0732 | rpsT | 0.072201416 | 0.02381866 | -0.029812302 | 1 |
| SPV_0733 | coaA | -0.175345463 | 0.132943646 | 0.321196931 | 1 |
| SPV_0734 | unknown | 0.161152132 | 0.033556708 | -0.115859097 | 1 |
| SPV_0735 | unknown | -0.104486086 | -0.015105418 | 0.101387643 | 1 |
| SPV_0736 | pdp | 0.236694302 | 0.058247857 | -0.166611774 | 1 |
| SPV_0738,SPV_0739,SPV_0737 | cdd-1,unknown,deoC | 0.15019781 | 0.202981634 | 0.070886743 | 1 |
| SPV_0740 | unknown | 0.102123444 | -0.060457263 | -0.149467296 | 1 |
| SPV_0741 | unknown | 0.185153123 | 0.198927466 | 0.029004966 | 1 |
| SPV_0742 | unknown | 0.071657414 | 0.008879899 | -0.049883315 | 1 |
| SPV_0745 | plsY | -0.966485644 | -0.626273008 | 0.357839861 | 1 |
| SPV_0746 | parE | -2.046609519 | -1.535838837 | 0.525866655 | 1 |
| SPV_0747 | unknown | -1.226864771 | -1.093376202 | 0.148919478 | 1 |
| SPV_0748 | parC | -1.501635011 | -1.542065765 | -0.024877263 | 1 |

| | | | | | |
|---|---|---|---|---|---|
| SPV_0750,SPV_0752,SPV_0751,SPV_0753,SPV_0749 | unknown,unknown,unknown,pcp1,ilvE | -0.153240763 | -0.574918408 | -0.406141344 | 1 |
| SPV_0754,SPV_0755 | unknown,tRNA-Tyr-1 | 0.124090003 | -0.352322831 | -0.460472027 | 1 |
| SPV_0755 | tRNA-Tyr-1 | -1.952501394 | -0.832434032 | 1.138987555 | 1 |
| SPV_0756 | tRNA-Gln-1 | -3.64922602 | -2.645201065 | 1.022418785 | 1 |
| SPV_0757 | rpsA | -0.931287872 | -0.566200835 | 0.378631919 | 1 |
| SPV_0758 | unknown | -0.060991623 | 0.115370777 | 0.186120804 | 1 |
| SPV_0759 | unknown | -2.922867201 | -1.713547952 | 1.22284467 | 1 |
| SPV_0760 | dnaX | -2.691161661 | -1.973020511 | 0.732402007 | 1 |
| SPV_0761 | unknown | -0.685129918 | -0.472822738 | 0.22292396 | 1 |
| SPV_0764,SPV_0766,SPV_0763,SPV_0762,SPV_0765 | sufS,sufB,sufD,sufC,sufE2 | -0.026104802 | 0.000581037 | 0.043177936 | 1 |
| SPV_0767 | pbp3 | 0.050628229 | 0.277083272 | 0.242828094 | 1 |
| SPV_0768 | cozE | -0.749825407 | -0.465352316 | 0.297602465 | 1 |
| SPV_0769 | ssrA | -0.03975881 | -0.204039213 | -0.153862827 | 1 |
| SPV_0770 | tRNA-Ser-1 | -0.008475388 | 0.114957357 | 0.133786557 | 1 |
| SPV_0771,SPV_0772 | fruR,fruB | -0.060298481 | 0.008856843 | 0.078577078 | 1 |
| SPV_0773 | fruA | 0.083288444 | 0.121703715 | 0.047214636 | 1 |
| SPV_0775 | unknown | 0.193187578 | 0.144138577 | -0.035363804 | 1 |
| SPV_0776,SPV_0777 | unknown,thiI | 0.07740133 | 0.115358432 | 0.053980346 | 1 |
| SPV_0783,SPV_0782,SPV_0785,SPV_0784 | unknown,unknown,unknown,unknown | 0.18163558 | -0.044120822 | -0.207267 | 1 |
| SPV_0785,SPV_0784 | unknown,unknown | 0.031582892 | 0.047646117 | 0.029942531 | 1 |
| SPV_0786,SPV_0787 | argR2,pepX | 0.035885962 | -0.010773036 | -0.031565674 | 1 |
| SPV_0788 | dnaE | -0.163846955 | -0.033104131 | 0.144218677 | 1 |
| SPV_0789,SPV_0790 | pfkA,pyk | -3.732428187 | -3.122910889 | 0.625037397 | 1 |
| SPV_0790 | pyk | -2.672110845 | -2.716003816 | -0.033079924 | 1 |
| SPV_0792,SPV_0793 | unknown,unknown | 0.075222049 | 0.055871525 | -0.002836532 | 1 |
| SPV_0794 | unknown | 0.099558094 | 0.006782579 | -0.075037153 | 1 |
| SPV_0795 | unknown | -0.10212864 | 0.016316237 | 0.133019475 | 1 |
| SPV_0796 | unknown | 0.151287353 | -0.066771576 | -0.204713994 | 1 |
| SPV_0797 | unknown | -0.060677465 | 0.033286466 | 0.102374519 | 1 |
| SPV_0798 | unknown | 0.035513583 | 0.032490787 | 0.013207444 | 1 |
| SPV_0799 | unknown | 0.014894824 | -0.0010922 | -0.005681596 | 1 |
| SPV_0802 | tex | 0.139655305 | -0.057808193 | -0.184124397 | 1 |
| SPV_0803 | unknown | -0.002881924 | 0.005776229 | 0.025905844 | 1 |
| SPV_0804 | unknown | 0.080929006 | 0.028001623 | -0.040292707 | 1 |
| SPV_0805 | unknown | 0.182936709 | 0.03274107 | -0.136720713 | 1 |
| SPV_0806 | unknown | 0.121693772 | 0.132832411 | 0.027277151 | 1 |
| SPV_0807 | unknown | 0.08519983 | 0.071033909 | 0.000926653 | 1 |
| SPV_0809 | cad | 0.036717991 | 0.036295785 | 0.013616385 | 1 |
| SPV_0810 | unknown | 0.015428805 | 0.007055199 | 0.006162554 | 1 |
| SPV_0811 | speE | -0.037595944 | -0.113782315 | -0.059322984 | 1 |

| | | | | | |
|---|---|---|---|---|---|
| SPV_0812 | lys1 | 0.263650857 | 0.058323036 | -0.188980923 | 1 |
| SPV_0813 | nspC | 0.045899867 | -0.017548176 | -0.049306232 | 1 |
| SPV_0814 | aguA | 0.160347159 | -0.043749261 | -0.189024678 | 1 |
| SPV_0815 | unknown | 0.115605623 | 0.065753765 | -0.034686908 | 1 |
| SPV_0816 | unknown | 0.058275647 | 0.113094107 | 0.070477358 | 1 |
| SPV_0819,SPV_0818 | lspA,cmbR | 0.037528735 | -0.051214825 | -0.073153156 | 1 |
| SPV_0820 | rluD1 | 0.22671289 | -0.20004876 | -0.410413368 | 1 |
| SPV_0821 | cbpE | 0.274563909 | -0.0043604 | -0.266608604 | 1 |
| SPV_0822 | proB | 0.042721717 | 0.22924933 | 0.202405348 | 1 |
| SPV_0823 | proA | -0.15774524 | 0.157516592 | 0.327561036 | 1 |
| SPV_0824 | proC | 0.367823578 | 0.274986195 | -0.071839266 | 1 |
| SPV_0828 | unknown | 0.143989704 | 1.146579697 | 1.018690222 | 1 |
| SPV_0829 | unknown | 0.038840221 | 0.259406088 | 0.2355578 | 1 |
| SPV_0831 | unknown | 0.157785778 | 0.06647837 | -0.077546006 | 1 |
| SPV_0833 | trmFO | 0.08250507 | -0.066279521 | -0.135816402 | 1 |
| SPV_0835 | frr | -0.27656583 | -0.516699878 | -0.224394508 | 1 |
| SPV_0836 | unknown | -0.092080882 | -0.643752889 | -0.538543127 | 1 |
| SPV_0837 | unknown | 0.050237616 | -0.041182488 | -0.080046854 | 1 |
| SPV_0838,SPV_2252,SPV_0839 | phoH,unknown,unknown | -0.167702414 | 0.17422726 | 0.354235429 | 1 |
| SPV_0841 | ald | -0.190059856 | 0.078076073 | 0.278641013 | 1 |
| SPV_0842 | unknown | 0.107060945 | 0.070021371 | -0.025312702 | 1 |
| SPV_0847 | infC | -3.115997533 | -2.06730544 | 1.063713849 | 1 |
| SPV_0848 | rpmI | -3.500989783 | -2.594483772 | 0.919600569 | 1 |
| SPV_0849 | rplT | -2.567715306 | -1.372393313 | 1.20976806 | 1 |
| SPV_0850 | gloA | 0.070413404 | -0.034590823 | -0.0903527 | 1 |
| SPV_0852,SPV_0853,SPV_0851 | pyrDb,lytB,pyrK | 0.13468418 | -0.025419671 | -0.14673705 | 1 |
| SPV_0853 | lytB | 0.097814596 | 0.041612521 | -0.041724013 | 1 |
| SPV_0854 | pavA | -0.215178869 | -0.021261582 | 0.206945174 | 1 |
| SPV_0855,SPV_0856 | ybeY,dgkA | -1.12346343 | -0.838014049 | 0.299956409 | 1 |
| SPV_0857 | era | -0.618298983 | -0.769916706 | -0.14022064 | 1 |
| SPV_0858 | mutM | -0.533674297 | -0.104428177 | 0.441328655 | 1 |
| SPV_0859 | coaE | -0.252284093 | -0.212250302 | 0.051150137 | 1 |
| SPV_0860 | pmrA | -0.048663457 | -0.085341078 | -0.020690767 | 1 |
| SPV_0861 | secG | -1.118405499 | -0.673134969 | 0.455133966 | 1 |
| SPV_0862 | rnr | -0.160056961 | -0.497840882 | -0.324566642 | 1 |
| SPV_0863 | smpB | -0.29193759 | -0.739275043 | -0.437271528 | 1 |
| SPV_0864 | tehB | -0.119885229 | -0.14025017 | -0.006323537 | 1 |
| SPV_0865 | coiA | -0.034195846 | 0.135409268 | 0.182203324 | 1 |
| SPV_0866,SPV_0867 | pepF1,unknown | 0.027845624 | 0.090261052 | 0.078955519 | 1 |
| SPV_0868 | prsA | 0.236498908 | 0.021608194 | -0.202747022 | 1 |
| SPV_0870 | gpmB2 | 0.044193365 | 0.045437587 | 0.014268241 | 1 |
| SPV_0871,SPV_0872 | ebsC,unknown | -0.10225661 | 0.328710603 | 0.445642423 | 1 |
| SPV_0873 | unknown | 0.259172096 | -0.022221852 | -0.266046502 | 1 |

| | | | | | |
|---|---|---|---|---|---|
| SPV_0875,SPV_0874 | unknown,glmU | -1.523645428 | -0.429152496 | 1.103730461 | 1 |
| SPV_0876 | unknown | -0.125608478 | -0.54356386 | -0.399463513 | 1 |
| SPV_0877 | mtnN | -0.036806093 | -0.601506785 | -0.549417326 | 1 |
| SPV_0879 | dnaQ | -0.068808187 | -0.032915084 | 0.048345991 | 1 |
| SPV_0880 | unknown | -0.688293235 | -0.192468198 | 0.51327091 | 1 |
| SPV_0883 | unknown | 0.012633091 | -0.005204458 | -0.00419332 | 1 |
| SPV_0884 | unknown | 0.275869018 | -0.009965871 | -0.265893368 | 1 |
| SPV_0886 | unknown | 0.105568099 | 0.068109057 | -0.026881831 | 1 |
| SPV_0886,SPV_0885 | unknown,ccdA-2 | -0.019973849 | 0.065175583 | 0.099642829 | 1 |
| SPV_0887 | yfnA | 0.185860039 | 0.010095937 | -0.157782531 | 1 |
| SPV_0890 | phtE | 0.007047671 | 0.06187212 | 0.066404229 | 1 |
| SPV_0894 | pepT | 0.196132394 | -0.118381257 | -0.304320455 | 1 |
| SPV_0895 | hemH | 0.190572089 | 0.08421996 | -0.095433297 | 1 |
| SPV_0896 | mscL | 0.016625209 | -0.026716051 | -0.028567718 | 1 |
| SPV_0897 | mesH | 0.01117301 | 0.131673985 | 0.134557165 | 1 |
| SPV_0898 | queT | 0.063319792 | 0.021667332 | -0.031623709 | 1 |
| SPV_0899 | tRNA-Thr-1 | 0.061086222 | 0.037103147 | -0.014198967 | 1 |
| SPV_0901,SPV_0900 | dapA,asd | -0.06390988 | 0.117356454 | 0.194233685 | 1 |
| SPV_0902 | mnmE | -0.074924861 | -0.3122835 | -0.223927865 | 1 |
| SPV_0903 | xylH | -0.125929483 | 0.003800907 | 0.14497975 | 1 |
| SPV_0904 | tdk | -0.890259848 | -0.806967043 | 0.098823604 | 1 |
| SPV_0905 | bltD | -1.098153933 | -0.773527278 | 0.338699742 | 1 |
| SPV_0906 | prfA | -0.708452043 | -0.291801594 | 0.431934064 | 1 |
| SPV_0907 | hemK | -0.757014248 | -0.410706541 | 0.358356635 | 1 |
| SPV_0908 | tsaC | -1.143811312 | -0.416789611 | 0.738000501 | 1 |
| SPV_0909 | unknown | -0.739769992 | -0.281462677 | 0.470441003 | 1 |
| SPV_0910 | glyA | -0.203790857 | 0.058803355 | 0.276645335 | 1 |
| SPV_0911 | unknown | -0.067334478 | 0.349858298 | 0.428263599 | 1 |
| SPV_0912 | pvaA | 0.010449612 | 0.328566062 | 0.337578352 | 1 |
| SPV_0913 | unknown | -0.116704485 | 0.1360458 | 0.266919467 | 1 |
| SPV_0914 | rlmCD | 0.226632734 | -0.247147091 | -0.460380474 | 1 |
| SPV_0920 | unknown | -0.106577044 | -0.016321173 | 0.097535611 | 1 |
| SPV_0922,SPV_0921 | unknown,ccrB | -0.014131329 | 0.10989437 | 0.139820571 | 1 |
| SPV_0923 | unknown | 0.013820906 | 0.10834437 | 0.105777352 | 1 |
| SPV_0924 | unknown | -0.05460796 | 0.031402 | 0.09778559 | 1 |
| SPV_0925 | unknown | 0.09479987 | 0.009513156 | -0.074378824 | 1 |
| SPV_0926 | unknown | 0.155689765 | 0.026384172 | -0.110561716 | 1 |
| SPV_0934 | unknown | 0.133672909 | 0.019015266 | -0.101201075 | 1 |
| SPV_0935 | unknown | 0.104684826 | 0.13899639 | 0.04141621 | 1 |
| SPV_0939 | mutR2 | 0.018028775 | 0.041134502 | 0.035222708 | 1 |
| SPV_0940 | rffD | 0.314415965 | -0.076267966 | -0.376741363 | 1 |
| SPV_0941 | unknown | -0.051687702 | 0.039852545 | 0.103202056 | 1 |
| SPV_0943 | unknown | 0.170121313 | -0.123460995 | -0.275910106 | 1 |

| | | | | | |
|---|---|---|---|---|---|
| SPV_0944 | unknown | -0.069466345 | 0.095619969 | 0.177626287 | 1 |
| SPV_0945 | unknown | 0.121970465 | 0.169730808 | 0.064132646 | 1 |
| SPV_0946 | unknown | -0.035331518 | 0.056169724 | 0.100685956 | 1 |
| SPV_0947 | unknown | 0.107363443 | 0.139696928 | 0.052618683 | 1 |
| SPV_0948 | nikS | -0.004202983 | -0.10372746 | -0.085189973 | 1 |
| SPV_0949 | unknown | -0.035470292 | 0.077135173 | 0.125414896 | 1 |
| SPV_0950 | mefE | 0.023873397 | 0.020272195 | 0.012435707 | 1 |
| SPV_0951 | unknown | 0.107926393 | -0.006481454 | -0.097260237 | 1 |
| SPV_0953,SPV_0952 | ppc,ftsW | -0.928037793 | -0.496415762 | 0.448475612 | 1 |
| SPV_0955 | trpY | 0.129122072 | 0.152097139 | 0.035229477 | 1 |
| SPV_0956 | unknown | 0.369042342 | 0.02534242 | -0.330191547 | 1 |
| SPV_0959,SPV_0957,SPV_0958 | unknown,dnaG,rpoD | -2.98893312 | -2.211085241 | 0.782399073 | 1 |
| SPV_0960 | lafB | -0.666300851 | -0.545968211 | 0.132899936 | 1 |
| SPV_0961 | lafA | -0.305484924 | -0.123999953 | 0.19889402 | 1 |
| SPV_0962 | unknown | 0.030216612 | 0.214787164 | 0.201785467 | 1 |
| SPV_0963 | unknown | -1.029130498 | -0.71623139 | 0.325527644 | 1 |
| SPV_0964 | obgE | -0.621977783 | -0.438527678 | 0.201492836 | 1 |
| SPV_0965 | unknown | -0.62858646 | -0.484716186 | 0.152170785 | 1 |
| SPV_0969 | unknown | 0.250394948 | -0.437116882 | -0.670977229 | 1 |
| SPV_0970 | map | -0.059805742 | -0.719931578 | -0.644915178 | 1 |
| SPV_0973 | pcrA | -0.478359858 | -0.478293809 | 0.014862863 | 1 |
| SPV_0975 | radC | -0.03090732 | 0.11725429 | 0.159431812 | 1 |
| SPV_0976,SPV_0974 | rex,unknown | 0.11946245 | 0.11420837 | 0.015334203 | 1 |
| SPV_0977 | unknown | 0.011807125 | 0.056922322 | 0.053553897 | 1 |
| SPV_0978 | unknown | 0.164638055 | -0.08615241 | -0.231841441 | 1 |
| SPV_0979 | nifS | -1.192938342 | -0.825921126 | 0.382151616 | 1 |
| SPV_0980 | prs2 | -1.335135116 | -0.789909299 | 0.562141121 | 1 |
| SPV_0981 | unknown | 0.060709281 | 0.068992431 | 0.021687717 | 1 |
| SPV_0983,SPV_0984,SPV_0982,SPV_0985 | ppnK,rluD3,unknown,eutD | -2.207262306 | -1.305313739 | 0.913831227 | 1 |
| SPV_0986 | unknown | -0.046301582 | 0.089321663 | 0.146952408 | 1 |
| SPV_0987 | unknown | 0.022389432 | 0.107943066 | 0.09939433 | 1 |
| SPV_0988 | unknown | 0.158682721 | 0.041280784 | -0.104531939 | 1 |
| SPV_0989 | rplU | -3.503024019 | -2.587546163 | 0.928790578 | 1 |
| SPV_0990 | unknown | -3.226857485 | -2.46750071 | 0.773567471 | 1 |
| SPV_0991 | rpmA | -2.933174179 | -1.768044682 | 1.17882172 | 1 |
| SPV_0992 | unknown | 0.281229727 | 0.044108586 | -0.221731702 | 1 |
| SPV_0993 | unknown | -0.04051234 | -0.066281507 | -0.007677541 | 1 |
| SPV_0994 | ribF | -1.568886206 | -0.991191147 | 0.594145705 | 1 |
| SPV_0995 | unknown | -0.045116498 | 0.06130256 | 0.121850432 | 1 |
| SPV_0996 | unknown | -0.032792809 | -0.180026917 | -0.135633853 | 1 |
| SPV_0997 | hlpA | -0.572843177 | -1.614265102 | -1.024279839 | 1 |
| SPV_0999,SPV_0998,SPV_1000 | rggD,unknown,unknown | 0.29276653 | -0.059620369 | -0.334808702 | 1 |

| | | | | | |
|---|---|---|---|---|---|
| SPV_1000 | unknown | 0.023938501 | 0.068452508 | 0.061798895 | 1 |
| SPV_1001,SPV_1002 | ligA,pulA | -0.437690719 | -0.146412245 | 0.306614207 | 1 |
| SPV_1003 | unknown | 0.125084648 | 0.062266583 | -0.050391719 | 1 |
| SPV_1004 | gapN | -0.202415595 | 0.090353288 | 0.306327694 | 1 |
| SPV_1008,SPV_1007 | glgA,glgD | 0.155295103 | -0.025807768 | -0.162333107 | 1 |
| SPV_1008,SPV_1007,SPV_1005,SPV_1006 | glgA,glgD,glgB,glgC | 0.115933769 | 0.100525199 | -0.001740314 | 1 |
| SPV_1009 | serB | 0.091541577 | -0.076204932 | -0.146827404 | 1 |
| SPV_1010 | unknown | 0.110478629 | 0.055531741 | -0.039893579 | 1 |
| SPV_1011 | glxK | -0.026980517 | 0.108189708 | 0.14958452 | 1 |
| SPV_1012 | eno | -2.432016012 | -1.5492504 | 0.893465908 | 1 |
| SPV_1013 | unknown | -0.002809481 | 0.044790821 | 0.0613978 | 1 |
| SPV_1017,SPV_1015,SPV_1016 | unknown,rexB,rexA | -0.096661824 | -0.089917105 | 0.01999105 | 1 |
| SPV_1018 | zmpA | 0.155683145 | 0.044863374 | -0.096871737 | 1 |
| SPV_1020,SPV_1019 | rnhB,rbgA | -0.868948312 | -0.301663803 | 0.582591275 | 1 |
| SPV_1021 | unknown | 0.108756457 | 0.081962878 | -0.00653844 | 1 |
| SPV_1023 | xerS | 0.005233027 | 0.014384841 | 0.026212637 | 1 |
| SPV_1024 | lplA | -0.113947491 | 0.146329447 | 0.272867572 | 1 |
| SPV_1025 | acoL | 0.42460461 | -0.081529604 | -0.487531206 | 1 |
| SPV_1026 | acoC | 0.032173738 | 0.13521504 | 0.1171745 | 1 |
| SPV_1027 | acoB | 0.025701982 | 0.128072718 | 0.117299651 | 1 |
| SPV_1028 | acoA | 0.088129624 | 0.001504556 | -0.071214887 | 1 |
| SPV_1029 | pdrM | -0.209734089 | 0.058377044 | 0.277648815 | 1 |
| SPV_1030,SPV_1032,SPV_1031 | pyrC,ung,mutX | -0.06490864 | -0.032161625 | 0.049496929 | 1 |
| SPV_1034,SPV_1033 | unknown,unknown | 0.049239942 | 0.045950377 | 0.009635222 | 1 |
| SPV_1038 | phtA | 0.063388681 | -0.021919851 | -0.071516337 | 1 |
| SPV_1043 | nrdF | -1.48964232 | -0.643897181 | 0.85963518 | 1 |
| SPV_1044 | lacR | -0.162574987 | 0.064310249 | 0.239576404 | 1 |
| SPV_1046 | lacG-2 | 0.034629207 | 0.005038909 | -0.012629611 | 1 |
| SPV_1047 | lacE-2 | 0.159842883 | 0.093563427 | -0.04983648 | 1 |
| SPV_1048 | lacF-2 | 0.037581184 | 0.032180655 | -0.000259721 | 1 |
| SPV_1049 | lacT | 0.073563881 | 0.02399448 | -0.037141712 | 1 |
| SPV_1053,SPV_1052,SPV_1051,SPV_2301 | lacA,lacB,lacC,lacD | 0.177991141 | 0.039759787 | -0.11923537 | 1 |
| SPV_1054 | unknown | 0.19008383 | -0.016770096 | -0.192914641 | 1 |
| SPV_1057 | unknown | 0.030441386 | 0.052763818 | 0.033336291 | 1 |
| SPV_1059 | unknown | 0.338205238 | 0.120059435 | -0.198288239 | 1 |
| SPV_1060 | lepA | 0.06506946 | 0.02101503 | -0.029416668 | 1 |
| SPV_1061 | pphA | 0.224914135 | 0.026130914 | -0.182257813 | 1 |
| SPV_1062 | recN | -0.45381031 | -0.674861551 | -0.20388555 | 1 |
| SPV_1063 | ahrC | -0.333149395 | -0.196480473 | 0.154897455 | 1 |
| SPV_1064,SPV_1065 | unknown,ispA | -0.939265336 | 0.042146867 | 0.993025888 | 1 |
| SPV_1066 | xseB | -0.83302874 | -0.015046607 | 0.829864971 | 1 |

| | | | | | |
|---|---|---|---|---|---|
| SPV_1067 | xseA | -0.692955732 | -0.123075408 | 0.581749745 | 1 |
| SPV_1068 | udk | 0.198195668 | -0.031498477 | -0.208641323 | 1 |
| SPV_1069 | unknown | -0.156564469 | 0.142650807 | 0.315379375 | 1 |
| SPV_1070 | truB | -0.064653632 | 0.037208734 | 0.117492485 | 1 |
| SPV_1071 | unknown | -0.010423975 | 0.028137421 | 0.053148468 | 1 |
| SPV_1072 | unknown | -0.021110042 | -0.134497189 | -0.10079608 | 1 |
| SPV_1075 | nirC | 0.06338148 | 0.048550696 | -0.005854602 | 1 |
| SPV_1076 | srtA | -0.160638517 | -0.132466316 | 0.041457809 | 1 |
| SPV_1077 | gyrA | -1.69155829 | -0.825057665 | 0.880773186 | 1 |
| SPV_1078 | ldh | -0.425710664 | -0.066800134 | 0.374863043 | 1 |
| SPV_1079 | unknown | -0.026371265 | 0.109426262 | 0.154379294 | 1 |
| SPV_1080,SPV_1079 | unknown,unknown | 0.144622415 | 0.023595229 | -0.103521861 | 1 |
| SPV_1081,SPV_1082,SPV_1080,SPV_1079 | relE2,relB2,unknown,unknown | 0.29695257 | -0.173640659 | -0.457489929 | 1 |
| SPV_1083,SPV_1085,SPV_1084 | vicX,vicR,vicK | -2.467502345 | -2.022472795 | 0.460076495 | 1 |
| SPV_1083,SPV_1085,SPV_1086,SPV_1084 | vicX,vicR,mutY,vicK | -0.080918867 | 0.126225649 | 0.222655345 | 1 |
| SPV_1087 | fhs | -1.104495648 | -0.138351201 | 0.98237122 | 1 |
| SPV_1090,SPV_1088,SPV_1089 | panT,coaB,coaC | -0.032566553 | 0.013807515 | 0.058341568 | 1 |
| SPV_1091,SPV_1093 | niaX,niaR | 0.105423369 | 0.181915929 | 0.094685583 | 1 |
| SPV_1092 | unknown | -0.171107334 | -0.006524045 | 0.178118831 | 1 |
| SPV_1093 | niaR | 0.107875358 | 0.08236371 | -0.01413376 | 1 |
| SPV_1094 | unknown | 0.022953071 | 0.027678989 | 0.018383473 | 1 |
| SPV_1095 | unknown | 0.201523297 | 0.057977766 | -0.12088116 | 1 |
| SPV_1096 | uvrB | 0.328570105 | -0.063907529 | -0.375412886 | 1 |
| SPV_1097 | unknown | 0.049213493 | 0.043027989 | 0.00645022 | 1 |
| SPV_1099,SPV_1098 | glnQ5,glnHP5 | -1.836788605 | -0.559886898 | 1.290759013 | 1 |
| SPV_1100 | zwf | -1.300415541 | -0.460804453 | 0.853495968 | 1 |
| SPV_1101 | ftsY | -0.050009499 | -0.084649249 | -0.018766553 | 1 |
| SPV_1102 | unknown | 0.11770679 | -0.456564038 | -0.560130198 | 1 |
| SPV_1103 | unknown | -0.194946871 | -0.519995315 | -0.309274221 | 1 |
| SPV_1104 | smc | -0.439099133 | -0.671734182 | -0.215414047 | 1 |
| SPV_1105 | rnc | -0.659435802 | -0.341244327 | 0.328358359 | 1 |
| SPV_1107,SPV_1106 | guaC,tRNA-Arg-1 | -0.055321176 | 0.108459893 | 0.17582 | 1 |
| SPV_1108 | unknown | -0.066593605 | 0.080552332 | 0.159743648 | 1 |
| SPV_1109 | unknown | 0.063273357 | -0.081865288 | -0.127535939 | 1 |
| SPV_1110 | unknown | 0.294625335 | 0.076945286 | -0.206961181 | 1 |
| SPV_1111 | unknown | 0.078854075 | 0.016404587 | -0.044012379 | 1 |
| SPV_1112 | unknown | -0.020085338 | 0.042338719 | 0.07639425 | 1 |
| SPV_1113 | unknown | 0.335921362 | 0.018529277 | -0.298038341 | 1 |
| SPV_1115 | leuB | 0.400733773 | -0.014203546 | -0.396582057 | 1 |
| SPV_1116 | leuA | 0.006013635 | 0.001204176 | 0.009952362 | 1 |
| SPV_1117 | unknown | -0.013893253 | 0.051421045 | 0.078131612 | 1 |

| | | | | | |
|---|---|---|---|---|---|
| SPV_1118 | cutC | 0.061486948 | 0.003265079 | -0.043166572 | 1 |
| SPV_1118,SPV_1119 | cutC,unknown | 0.421719288 | -0.049971151 | -0.458277766 | 1 |
| SPV_1120 | topA | -0.321312659 | -0.543444546 | -0.206252971 | 1 |
| SPV_1121 | unknown | -0.033123536 | 0.1612624 | 0.201254543 | 1 |
| SPV_1122 | dprA | 0.057618386 | 0.142868884 | 0.093804049 | 1 |
| SPV_1130 | licD2 | 0.006957232 | 0.246820653 | 0.25948117 | 1 |
| SPV_1134,SPV_1131,SPV_1132,SPV_1133 | pyrR,carB,carA,pyrB | -0.048355211 | 0.043307813 | 0.105151329 | 1 |
| SPV_1135 | nth | 0.307594406 | -0.001286428 | -0.293425954 | 1 |
| SPV_1136 | unknown | 0.037325328 | -0.084688796 | -0.107616793 | 1 |
| SPV_1137 | unknown | 0.444021398 | -0.029622267 | -0.461352272 | 1 |
| SPV_1139,SPV_1138 | lemA,htpX | 0.206241159 | 0.041967105 | -0.148278088 | 1 |
| SPV_1140 | rsmG | -0.041047745 | 0.011265584 | 0.066484857 | 1 |
| SPV_1141 | uraA | -0.059847983 | 0.010442413 | 0.083295406 | 1 |
| SPV_1142 | ffh | -0.403951361 | -0.351876441 | 0.063930997 | 1 |
| SPV_1143 | unknown | -0.327368925 | -0.385591724 | -0.04555944 | 1 |
| SPV_1144 | unknown | 0.19441116 | -0.077277975 | -0.256797575 | 1 |
| SPV_1146,SPV_1145 | yidA,unknown | 0.011505976 | 0.205865625 | 0.206972037 | 1 |
| SPV_1147 | tRNA-Arg-2 | 0.031769095 | 0.102397397 | 0.089921586 | 1 |
| SPV_1148 | rplS | -3.316669668 | -2.618834349 | 0.70763298 | 1 |
| SPV_1151,SPV_1150,SPV_1149 | unknown,crcB2,crcB1 | 0.226369123 | 0.013195405 | -0.201532584 | 1 |
| SPV_1152 | fld | -0.102580135 | -0.034300737 | 0.082872035 | 1 |
| SPV_1153 | pde2 | -0.027062391 | 0.005035447 | 0.043007637 | 1 |
| SPV_1154 | rpmE2 | -1.476665984 | -1.766519849 | -0.272713431 | 1 |
| SPV_1155 | efeU | 0.155885728 | -0.046089512 | -0.186174848 | 1 |
| SPV_1156 | efeB | 0.009085421 | 0.034355769 | 0.041182328 | 1 |
| SPV_1158 | gdhA | -0.092044369 | -0.051752134 | 0.051408108 | 1 |
| SPV_1159 | unknown | 0.081615079 | 0.077078971 | 0.01249956 | 1 |
| SPV_1160 | unknown | 0.163314028 | 0.020717924 | -0.127718755 | 1 |
| SPV_1162 | unknown | 0.078650471 | 0.023227044 | -0.043111693 | 1 |
| SPV_1164 | cdd-2 | 0.041304666 | -0.03892222 | -0.067461425 | 1 |
| SPV_1165 | unknown | 0.243724588 | -0.072832483 | -0.305043915 | 1 |
| SPV_1166 | unknown | -0.027159339 | 0.138225791 | 0.179003374 | 1 |
| SPV_1167 | appD | 0.249106421 | 0.029443743 | -0.203433856 | 1 |
| SPV_1168 | appC | -0.144081827 | 0.083232909 | 0.239971975 | 1 |
| SPV_1169 | appB | -0.07936388 | 0.157452612 | 0.253811275 | 1 |
| SPV_1173,SPV_1170,SPV_1171,SPV_1172 | unknown,appA,unknown,nanE-2 | 0.088936667 | 0.028626175 | -0.049811289 | 1 |
| SPV_1174 | unknown | -0.036382115 | -0.045366809 | 0.004017188 | 1 |
| SPV_1175 | unknown | 0.186460273 | 0.062285917 | -0.111132655 | 1 |
| SPV_1176 | unknown | 0.04742884 | 0.027370664 | -0.008282889 | 1 |
| SPV_1177 | unknown | 0.097169136 | -0.018633039 | -0.100338825 | 1 |
| SPV_1178 | ptrB | 0.35730905 | 0.282121223 | -0.063140177 | 1 |

| | | | | | |
|---|---|---|---|---|---|
| SPV_1179 | lanL | -0.20624992 | 0.051983845 | 0.275429278 | 1 |
| SPV_1182,SPV_2329,SPV_2327,SPV_1185,SPV_1180,SPV_2328 | unknown,unknown,unknown,unknown,unknown,unknown | -0.056795863 | 0.043073047 | 0.111364348 | 1 |
| SPV_1186 | unknown | -0.147562148 | 0.115457266 | 0.276329152 | 1 |
| SPV_1187 | rplL | -0.278821141 | -0.238756308 | 0.053444996 | 1 |
| SPV_1188 | rplJ | -2.248782962 | -0.951395421 | 1.306179619 | 1 |
| SPV_1189 | unknown | -0.018738497 | 0.071959603 | 0.10886508 | 1 |
| SPV_1190 | trzA | 0.009387248 | 0.253333726 | 0.260485009 | 1 |
| SPV_1191 | unknown | 0.015224271 | -0.081674318 | -0.088550967 | 1 |
| SPV_1192 | unknown | -0.048570332 | -0.008452954 | 0.056436339 | 1 |
| SPV_1193 | msrAB1 | 0.04858765 | -0.022079781 | -0.053549449 | 1 |
| SPV_1195,SPV_1193,SPV_2331,SPV_1194 | hom,msrAB1,unknown,thrB | -0.114768694 | 0.320945491 | 0.454898905 | 1 |
| SPV_1196 | mecA | 0.080062051 | -0.211045617 | -0.275096165 | 1 |
| SPV_1205,SPV_1203,SPV_1204,SPV_1202 | aroA,pheA,aroK,psr | -0.01993443 | 0.169464976 | 0.203118026 | 1 |
| SPV_1206 | unknown | 0.086741765 | 0.070877664 | -0.002639473 | 1 |
| SPV_1207 | tyrA | -0.041794807 | 0.148027004 | 0.204813629 | 1 |
| SPV_1208 | aroC | -0.050486692 | 0.167313126 | 0.233272512 | 1 |
| SPV_1209 | aroB | -0.018678694 | 0.072518868 | 0.102946712 | 1 |
| SPV_1210 | aroE | -0.081341348 | -0.016425698 | 0.075387694 | 1 |
| SPV_1211 | aroD | 0.036546685 | -0.252999057 | -0.275110195 | 1 |
| SPV_1212 | ywbD | -0.065852438 | -0.216600521 | -0.136053698 | 1 |
| SPV_1213,SPV_1214 | unknown,unknown | 0.107184973 | 0.11784013 | 0.020125265 | 1 |
| SPV_1215 | amy | 0.053343751 | 0.122169022 | 0.078583973 | 1 |
| SPV_1218,SPV_1220,SPV_1219,SPV_1221,SPV_1222 | potD,potB,potC,potA,murB | -5.686369945 | -5.300920377 | 0.395468649 | 1 |
| SPV_1223 | unknown | -0.15848115 | 0.061750229 | 0.232168941 | 1 |
| SPV_1224 | budA | -0.07278851 | 0.059588107 | 0.145975991 | 1 |
| SPV_1225 | unknown | 0.066971627 | 0.069647709 | 0.018878245 | 1 |
| SPV_1226 | unknown | 0.316203712 | 0.079598995 | -0.217305517 | 1 |
| SPV_1227 | phoU2 | 0.111358099 | 0.023153146 | -0.072098628 | 1 |
| SPV_1228 | pstB2-2 | 0.035729408 | 0.043613331 | 0.023396238 | 1 |
| SPV_1229 | pstB2-1 | 0.148817425 | -0.093147523 | -0.226908853 | 1 |
| SPV_1230 | pstA2 | -0.073263723 | -0.061694771 | 0.027667977 | 1 |
| SPV_1231 | pstC2 | 0.056310125 | 0.043979009 | 0.003796141 | 1 |
| SPV_1232 | pstS2 | -0.01546767 | -0.120096233 | -0.091687734 | 1 |
| SPV_1233 | unknown | 0.113804926 | 0.048152806 | -0.049717069 | 1 |
| SPV_1234,SPV_1235,SPV_1236 | unknown,unknown,spxA1 | 0.034141837 | 0.213529858 | 0.19206631 | 1 |
| SPV_1237 | unknown | -0.03509922 | 0.028603455 | 0.076549006 | 1 |
| SPV_1238,SPV_1240,SPV_1239 | nagD,hemN,unknown | 0.036731717 | 0.113062586 | 0.090188789 | 1 |
| SPV_1241 | unknown | 0.061378951 | 0.115840931 | 0.061132572 | 1 |
| SPV_1242 | unknown | -0.044139214 | 0.120596044 | 0.175787824 | 1 |

| | | | | | |
|---|---|---|---|---|---|
| SPV_1244,SPV_1243 | hprK,lgt | -0.061579454 | 0.173582089 | 0.249870686 | 1 |
| SPV_1245 | rpsU | -1.426693388 | -2.139111079 | -0.696251019 | 1 |
| SPV_1246 | nagB | 0.210567971 | -0.081846019 | -0.276906274 | 1 |
| SPV_1247 | queA | -0.08749174 | -0.00109589 | 0.102233108 | 1 |
| SPV_1248 | cbpM | -0.061709309 | -0.003623176 | 0.075622851 | 1 |
| SPV_1249,SPV_1250,SPV_1251 | unknown,nadE,pncB | -0.420293349 | -0.461790004 | -0.032473351 | 1 |
| SPV_1252 | unknown | 0.195639909 | -0.015873443 | -0.198833924 | 1 |
| SPV_1253 | unknown | 0.140744428 | -0.111826022 | -0.240142688 | 1 |
| SPV_1254 | unknown | -0.190833592 | 0.016934269 | 0.220460076 | 1 |
| SPV_1255 | unknown | -0.001429296 | -0.004983689 | 0.01136797 | 1 |
| SPV_1256 | unknown | 0.223162972 | -0.015144112 | -0.22285476 | 1 |
| SPV_1257 | unknown | 0.037530211 | 0.055133214 | 0.032413843 | 1 |
| SPV_1258 | unknown | 0.091814775 | -0.103597505 | -0.180483437 | 1 |
| SPV_1260,SPV_1259 | unknown,unknown | 0.07581659 | 0.073638641 | 0.008543985 | 1 |
| SPV_1261 | unknown | 0.16447722 | 0.026299571 | -0.121058514 | 1 |
| SPV_1262 | unknown | 0.2033209 | 0.087806673 | -0.10064914 | 1 |
| SPV_1263 | unknown | -0.090134765 | 0.076473927 | 0.181043553 | 1 |
| SPV_1264 | unknown | -0.026271718 | 0.060061778 | 0.098467448 | 1 |
| SPV_1265 | unknown | 0.107478713 | -0.036301874 | -0.128781357 | 1 |
| SPV_1266 | unknown | 0.236490908 | 0.021496395 | -0.20050481 | 1 |
| SPV_1267 | unknown | 0.025001205 | 0.008013726 | -0.001530807 | 1 |
| SPV_1272 | unknown | 0.123073965 | 0.012421462 | -0.098187337 | 1 |
| SPV_1273 | unknown | 0.041854921 | 0.074741472 | 0.053908979 | 1 |
| SPV_1274 | guaA | 0.056718476 | 0.424979502 | 0.382130568 | 1 |
| SPV_1275 | nagR | 0.160171215 | 0.094376146 | -0.045725416 | 1 |
| SPV_1276 | unknown | 0.125333977 | 0.110042695 | -0.00022941 | 1 |
| SPV_1277 | unknown | -0.127864096 | 0.076560277 | 0.217018053 | 1 |
| SPV_1278 | cppA | 0.078913773 | 0.140058038 | 0.082066187 | 1 |
| SPV_1279 | unknown | 0.058567356 | -0.023409994 | -0.069706594 | 1 |
| SPV_1280,SPV_2338 | unknown,capA | 0.241724929 | -0.040348179 | -0.265461894 | 1 |
| SPV_1284 | unknown | 0.032564844 | 0.211489991 | 0.193358926 | 1 |
| SPV_1285 | def1 | 0.083383156 | -0.393240639 | -0.459700565 | 1 |
| SPV_1286 | trmH | 0.189867579 | -0.113566942 | -0.289440357 | 1 |
| SPV_1287 | trxB | -0.690274522 | -0.051068395 | 0.664042072 | 1 |
| SPV_1288 | unknown | -0.056352958 | 0.860685107 | 0.934235511 | 1 |
| SPV_1290,SPV_1289 | unknown,unknown | 0.034810108 | -0.240352616 | -0.260687508 | 1 |
| SPV_1292,SPV_1293,SPV_1291 | ogt,unknown,unknown | 0.053258766 | -0.034318214 | -0.07341709 | 1 |
| SPV_1294,SPV_1295 | unknown,unknown | 0.068864478 | 0.111550793 | 0.054390402 | 1 |
| SPV_1296 | pdxT | -0.126667654 | 0.065319885 | 0.204907775 | 1 |
| SPV_1297 | pdxS | 0.031008338 | 0.05531811 | 0.037329854 | 1 |
| SPV_1298 | nox | 0.231262708 | 0.224536335 | 0.008413071 | 1 |
| SPV_1299 | unknown | -0.132157508 | 0.04734989 | 0.193052879 | 1 |
| SPV_1302,SPV_1301,SPV_1300 | unknown,unknown,apbE | 0.028367822 | 0.082223115 | 0.077030331 | 1 |

| | | | | | |
|---|---|---|---|---|---|
| SPV_1308 | unknown | 0.153358299 | 0.021064513 | -0.115606507 | 1 |
| SPV_1309,SPV_1308 | pgdA,unknown | -0.041966389 | -0.276681119 | -0.224873902 | 1 |
| SPV_1311 | mocA | 0.137563964 | -0.09693398 | -0.205652875 | 1 |
| SPV_1312 | yfmL | -0.014402924 | -0.041475981 | -0.013824537 | 1 |
| SPV_1315 | unknown | 0.039561331 | -0.06923223 | -0.09279877 | 1 |
| SPV_1317 | unknown | 0.105600823 | 0.105642722 | 0.009846818 | 1 |
| SPV_1318 | tuf | -3.540406783 | -2.424162794 | 1.129317057 | 1 |
| SPV_1319 | unknown | 0.046257122 | 0.150598923 | 0.117879783 | 1 |
| SPV_1320 | unknown | -0.347116169 | 0.354535883 | 0.718697854 | 1 |
| SPV_1321 | mucB | 0.167089065 | -0.037493586 | -0.187059721 | 1 |
| SPV_1322 | unknown | 0.15347969 | -0.020368561 | -0.152646595 | 1 |
| SPV_1323 | unknown | -0.044034512 | 0.082838364 | 0.143889481 | 1 |
| SPV_1326 | pgm | -0.051243675 | 0.348773079 | 0.41564127 | 1 |
| SPV_1327 | bta | 0.05005107 | 0.976288167 | 0.93912999 | 1 |
| SPV_1329,SPV_1328,SPV_1330 | glnQ6,glnH6,glnP6 | 0.229192516 | 0.142748912 | -0.080093185 | 1 |
| SPV_1331,SPV_1332 | unknown,unknown | 0.208304434 | 0.172801329 | -0.023585703 | 1 |
| SPV_1333 | unknown | 0.012482458 | 0.037968223 | 0.040272667 | 1 |
| SPV_1339,SPV_1340,SPV_1338,SPV_1336,SPV_1337,SPV_1341,SPV_1335,SPV_1334 | atpF,atpB,atpH,atpG,atpA,atpE,atpD,atpC | -1.579570735 | -0.71994777 | 0.882471589 | 1 |
| SPV_1343 | unknown | 0.308646435 | 0.065312425 | -0.23167756 | 1 |
| SPV_1344 | unknown | 0.102384464 | 0.141038716 | 0.053986148 | 1 |
| SPV_1346,SPV_1345 | mltG,greA | -0.178904478 | 0.196172015 | 0.392177139 | 1 |
| SPV_1347 | unknown | -0.210077147 | 0.441765778 | 0.668180659 | 1 |
| SPV_1348 | unknown | -5.868878204 | -4.742987546 | 1.13604635 | 1 |
| SPV_1349 | murC | -5.811270469 | -4.810097964 | 1.025590035 | 1 |
| SPV_1350 | unknown | -6.209813239 | -4.907872333 | 1.316478883 | 1 |
| SPV_1351 | snf | 0.041053448 | 0.752644512 | 0.726926464 | 1 |
| SPV_1353,SPV_1352 | metB,patB2 | 0.354237389 | 0.111634453 | -0.232192628 | 1 |
| SPV_1357 | aliB | 0.112569758 | -0.022450725 | -0.124130413 | 1 |
| SPV_1359 | murE | -5.272344754 | -4.215208335 | 1.063028069 | 1 |
| SPV_1360 | unknown | 0.103867498 | 0.001241117 | -0.088492444 | 1 |
| SPV_1361 | unknown | 0.070710454 | 0.043726404 | -0.015799725 | 1 |
| SPV_1362 | unknown | 0.101661276 | 0.095889224 | 0.01462142 | 1 |
| SPV_1363 | ppaC | -1.233570957 | -0.505375524 | 0.746637303 | 1 |
| SPV_1365,SPV_1364 | unknown,unknown | 0.006573844 | 0.156680579 | 0.162018066 | 1 |
| SPV_1367,SPV_1365,SPV_1366,SPV_1364 | unknown,unknown,unknown,unknown | -0.126469308 | 0.424716512 | 0.564716818 | 1 |
| SPV_1369,SPV_1368 | ssbA,rpsR | -3.194795221 | -2.335230496 | 0.872422887 | 1 |
| SPV_1370 | rpsF | -4.063938228 | -3.193608232 | 0.885133914 | 1 |
| SPV_1371 | asnC | -2.676223116 | -1.567219789 | 1.12111493 | 1 |
| SPV_1372 | unknown | -2.571246448 | -1.337945142 | 1.249250994 | 1 |
| SPV_1375 | unknown | 0.183258595 | 0.072016744 | -0.093967245 | 1 |

| | | | | | |
|---|---|---|---|---|---|
| SPV_1376 | pclA | 0.281124944 | 0.105450227 | -0.163661211 | 1 |
| SPV_1377 | mga1 | 0.037675597 | 0.018145823 | -0.003410692 | 1 |
| SPV_1378 | yaaA | 0.096789459 | 0.042255998 | -0.037859007 | 1 |
| SPV_1379 | unknown | 0.175964365 | -0.024180711 | -0.190308805 | 1 |
| SPV_1381,SPV_1380,SPV_1379 | def2,unknown,unknown | 0.056806161 | -0.027223175 | -0.068354604 | 1 |
| SPV_1383,SPV_1382 | pacL,unknown | 0.156540615 | 0.134378546 | -0.008409667 | 1 |
| SPV_1384 | mntE | -0.055908119 | 0.05402317 | 0.124328133 | 1 |
| SPV_1385,SPV_1389,SPV_1388,SPV_1387,SPV_1386 | unknown,unknown,unknown,dapB,cca | -0.654939681 | -0.420664123 | 0.248459245 | 1 |
| SPV_1390,SPV_1391,SPV_1392 | glmM,unknown,disA | -1.400208811 | -0.142166914 | 1.274294669 | 1 |
| SPV_1393 | unknown | 0.249988492 | -0.249244937 | -0.484074798 | 1 |
| SPV_1394,SPV_1396,SPV_1395 | unknown,unknown,unknown | -0.002191873 | -0.047520606 | -0.032426022 | 1 |
| SPV_1397 | aldR | 0.028349867 | -0.03113384 | -0.047975479 | 1 |
| SPV_1398 | engB | -0.929441176 | -0.728701802 | 0.214674955 | 1 |
| SPV_1399 | clpX | -2.056977152 | -1.059221255 | 1.012699888 | 1 |
| SPV_1400 | unknown | -0.249905148 | -0.240150879 | 0.02370055 | 1 |
| SPV_1401 | folA | -2.907834565 | -1.661787631 | 1.264976208 | 1 |
| SPV_1402 | dpr | -0.127509059 | 0.056215908 | 0.197144229 | 1 |
| SPV_1404,SPV_1403 | tpiA,lytC | -1.41125382 | -1.540582431 | -0.113894486 | 1 |
| SPV_1405,SPV_1406 | dnaD,metA | -2.654112112 | -2.485171557 | 0.185246992 | 1 |
| SPV_1407 | apt | -0.118989289 | -0.033743666 | 0.098828821 | 1 |
| SPV_1408 | unknown | 0.015287707 | -0.047419468 | -0.070518967 | 1 |
| SPV_1409 | msmK | 0.200400663 | 0.052046769 | -0.134672465 | 1 |
| SPV_1410 | tRNA-Leu-1 | -0.162591969 | -0.062723839 | 0.114969731 | 1 |
| SPV_1411,SPV_1412 | unknown,codY | 0.109287422 | 0.1539221 | 0.056605744 | 1 |
| SPV_1413 | cshA | -0.040236274 | 0.18358138 | 0.235841639 | 1 |
| SPV_1414 | oxlT | 0.027959637 | -0.018945212 | -0.032307956 | 1 |
| SPV_1415 | merA | 0.247974323 | 0.105049779 | -0.128321587 | 1 |
| SPV_1419,SPV_1418 | unknown,pepQ | 0.037304555 | 0.073758966 | 0.050958913 | 1 |
| SPV_1420 | unknown | 0.178223431 | 0.033460385 | -0.12881515 | 1 |
| SPV_1423,SPV_1424,SPV_1422 | pdxK,truA,pdxU2 | -1.573450835 | -0.854327415 | 0.732948517 | 1 |
| SPV_1425 | unknown | 0.177876152 | -0.029100934 | -0.188303169 | 1 |
| SPV_1426 | unknown | 0.056507594 | 0.102128724 | 0.055535646 | 1 |
| SPV_1427 | phnA | -0.037550183 | 0.195017546 | 0.25095525 | 1 |
| SPV_1429 | unknown | -1.068540357 | 0.073001704 | 1.155959975 | 1 |
| SPV_1430 | fer | -0.519425108 | 0.26558266 | 0.797394391 | 1 |
| SPV_1431,SPV_1432 | gtrB,galE-1 | 0.104538681 | 0.32946543 | 0.249271804 | 1 |
| SPV_1433 | unknown | 0.154877803 | -0.004946815 | -0.143666509 | 1 |
| SPV_1434 | ybgI | -0.247754576 | -0.0545832 | 0.206766947 | 1 |
| SPV_1435 | unknown | -0.099002174 | -0.00727421 | 0.11280027 | 1 |
| SPV_1436 | ctpE | 0.168656135 | 0.305277945 | 0.161317884 | 1 |
| SPV_1437 | plsC | -0.794424618 | 0.155079249 | 0.963771812 | 1 |
| SPV_1438 | cadD | -0.154527968 | 0.110625871 | 0.27780881 | 1 |

| | | | | | |
|---|---|---|---|---|---|
| SPV_1439 | rpsO | 0.115963191 | 0.026030534 | -0.073809688 | 1 |
| SPV_1440 | unknown | 0.025050275 | -0.017020117 | -0.028600753 | 1 |
| SPV_1441 | unknown | 0.163815961 | 0.080876763 | -0.069902234 | 1 |
| SPV_1446,SPV_1445 | unknown,unknown | 0.081618907 | 0.090159299 | 0.023852989 | 1 |
| SPV_1447 | unknown | 0.098614633 | -0.040834831 | -0.124440292 | 1 |
| SPV_1448 | unknown | -0.064484407 | -0.211784917 | -0.139014379 | 1 |
| SPV_1449 | unknown | 0.086466468 | 0.056650864 | -0.013431623 | 1 |
| SPV_1451,SPV_1450 | unknown,mntR | -0.183378422 | -0.216009035 | -0.019936132 | 1 |
| SPV_1454 | unknown | -0.070030359 | -0.022569792 | 0.066757448 | 1 |
| SPV_1457 | dtd | 0.438244922 | -0.030165847 | -0.450626415 | 1 |
| SPV_1458 | relA | 0.230612618 | 0.003559075 | -0.212501951 | 1 |
| SPV_1459 | unknown | 0.039200546 | 0.069735911 | 0.047950885 | 1 |
| SPV_1460 | pepO | 0.347412267 | 0.027751751 | -0.304798857 | 1 |
| SPV_1461,SPV_1463,SPV_1462 | psaB,psaA,psaC | -0.472186419 | -0.056610849 | 0.432002827 | 1 |
| SPV_1464 | psaD | -0.111075067 | -0.052670671 | 0.069055403 | 1 |
| SPV_1465 | unknown | 0.156515457 | 0.119187273 | -0.018225652 | 1 |
| SPV_1466 | unknown | 0.042787042 | 0.026388995 | 0.001481431 | 1 |
| SPV_1467 | unknown | 0.012051748 | 0.03955962 | 0.042623416 | 1 |
| SPV_1468 | gpmA | -1.413663822 | -1.00117457 | 0.425244395 | 1 |
| SPV_1469 | unknown | -0.018096457 | 0.266601169 | 0.296415647 | 1 |
| SPV_1474 | divIVA | -1.918729283 | -1.137643408 | 0.792799775 | 1 |
| SPV_1478 | ylmE | -0.080385129 | 0.270995521 | 0.366862009 | 1 |
| SPV_1480,SPV_1479 | ftsA,ftsZ | -4.613265025 | -3.298091227 | 1.329484348 | 1 |
| SPV_1481 | unknown | -0.057272993 | 0.252251408 | 0.32818745 | 1 |
| SPV_1482 | unknown | -0.024347304 | 0.500476715 | 0.537475031 | 1 |
| SPV_1483 | murF | -3.130615335 | -1.855288785 | 1.28901686 | 1 |
| SPV_1485 | recR | 0.229309692 | 0.400985711 | 0.18351438 | 1 |
| SPV_1486 | pbp2b | 0.196940375 | -0.06353304 | -0.246274912 | 1 |
| SPV_1487 | nanR | -0.014730513 | 0.187959145 | 0.218703315 | 1 |
| SPV_1488 | nanK | 0.030796026 | 0.042957612 | 0.022067047 | 1 |
| SPV_1490 | unknown | -0.140236121 | 0.08219366 | 0.230967714 | 1 |
| SPV_1491 | unknown | 0.089529221 | 0.086822836 | 0.014403385 | 1 |
| SPV_1492 | yjgK | 0.184051802 | 0.003954777 | -0.16805633 | 1 |
| SPV_1493 | satC | -0.09218115 | 0.019275217 | 0.130685282 | 1 |
| SPV_1494 | satB | -0.169808546 | 0.169901791 | 0.351693223 | 1 |
| SPV_1495 | satA | -0.085958519 | -0.007764998 | 0.092203053 | 1 |
| SPV_1496 | nanP | 0.261714859 | -0.071946644 | -0.327610143 | 1 |
| SPV_1497 | nanE-1 | -0.035939523 | 0.101168149 | 0.141366195 | 1 |
| SPV_1498 | unknown | -0.006648499 | 0.077874228 | 0.09870686 | 1 |
| SPV_1499 | nanB | -0.053077939 | 0.024205712 | 0.090977644 | 1 |
| SPV_1500 | unknown | -0.036368285 | 0.039923776 | 0.08877377 | 1 |
| SPV_1501 | ycjO | 0.078485954 | 0.017111546 | -0.043787229 | 1 |
| SPV_1502 | unknown | -0.132415322 | 0.102948808 | 0.248641916 | 1 |

| | | | | | |
|---|---|---|---|---|---|
| SPV_1503 | unknown | 0.110636931 | 0.073052294 | -0.024166101 | 1 |
| SPV_1504,SPV_1505 | nanA,unknown | 0.07299385 | 0.007289126 | -0.052110037 | 1 |
| SPV_1506 | axe1 | -0.131601833 | 0.113674284 | 0.257824499 | 1 |
| SPV_1507 | recG | 0.23998675 | 0.131172041 | -0.094149331 | 1 |
| SPV_1508 | alr | -2.23272193 | -1.220747948 | 1.022483446 | 1 |
| SPV_1509 | acpS | -1.864910119 | -1.110272937 | 0.769926052 | 1 |
| SPV_1510 | aroF | -2.058977094 | -1.077874791 | 0.996493387 | 1 |
| SPV_1511 | aroG | -1.324119872 | -0.791082808 | 0.554098991 | 1 |
| SPV_1512 | secA | -1.641068964 | -0.712923115 | 0.948291817 | 1 |
| SPV_1513 | unknown | -0.105042171 | 0.08740867 | 0.206328483 | 1 |
| SPV_1514 | unknown | 0.01662582 | 0.18906893 | 0.186564118 | 1 |
| SPV_1515 | unknown | 0.221911262 | 0.121338058 | -0.089814559 | 1 |
| SPV_1516 | unknown | 0.094782635 | 0.052984154 | -0.026592093 | 1 |
| SPV_1517 | unknown | 0.244832716 | 0.080272177 | -0.151099749 | 1 |
| SPV_1518 | unknown | -0.004201551 | 0.152887677 | 0.168792291 | 1 |
| SPV_1519 | engA | -1.008541271 | -0.486701882 | 0.535836064 | 1 |
| SPV_1520 | frp | -2.616858359 | -1.511610618 | 1.12046161 | 1 |
| SPV_1521 | dnaI | -3.493709096 | -2.276449466 | 1.234187384 | 1 |
| SPV_1522,SPV_1523 | dnaB,nrdR | -3.960508679 | -2.663870992 | 1.310364587 | 1 |
| SPV_1526,SPV_1524,SPV_1525 | unknown,gntR,unknown | -0.058131704 | 0.172354987 | 0.247497904 | 1 |
| SPV_1527,SPV_1528,SPV_2383 | qsrB,qsrA,srf-22 | 0.117197322 | -0.019152699 | -0.121070909 | 1 |
| SPV_1529 | unknown | 0.19379341 | 0.068562594 | -0.11012813 | 1 |
| SPV_1530 | unknown | 0.056114524 | 0.099361536 | 0.058912552 | 1 |
| SPV_1531 | scrK | -0.050625856 | 0.194666903 | 0.261849026 | 1 |
| SPV_1532 | scrA | -0.19658711 | 0.057319098 | 0.268232741 | 1 |
| SPV_1533 | unknown | -0.167561933 | 0.015996952 | 0.195851897 | 1 |
| SPV_1534,SPV_1535 | scrB,scrR | -1.232112549 | -1.261300052 | -0.01525465 | 1 |
| SPV_1536,SPV_1537 | mvaA,mvaS | -0.686051981 | 0.599565072 | 1.298412299 | 1 |
| SPV_1538 | unknown | -0.069769618 | 0.113643382 | 0.200642538 | 1 |
| SPV_1539 | unknown | 0.034481434 | 0.156959223 | 0.135503688 | 1 |
| SPV_1540 | unknown | 0.299532657 | 0.018319466 | -0.263615663 | 1 |
| SPV_1541 | unknown | 0.069839133 | 0.083622556 | 0.02735039 | 1 |
| SPV_1543,SPV_1542 | phpP,stkP | -0.261550445 | -0.316106777 | -0.040922533 | 1 |
| SPV_1544 | sun | 0.082983231 | -0.034927866 | -0.102814252 | 1 |
| SPV_1545 | fmt | -0.406723616 | -0.121030845 | 0.302151892 | 1 |
| SPV_1546 | priA | -0.495885229 | -0.558306193 | -0.049778514 | 1 |
| SPV_1549 | rny | -0.232490636 | -0.011211532 | 0.237894096 | 1 |
| SPV_1551,SPV_1550 | yefM,yoeB | -0.052071907 | -0.215391251 | -0.147047049 | 1 |
| SPV_1552 | unknown | -0.334936704 | -0.226672692 | 0.124138287 | 1 |
| SPV_1553 | unknown | -0.102457449 | -0.176474536 | -0.057845535 | 1 |
| SPV_1554 | rsfA | -0.362826275 | 0.023772777 | 0.401772447 | 1 |

| | | | | | |
|---|---|---|---|---|---|
| SPV_1557,SPV_1556,SPV_1558,SPV_1559,SPV_1561,SPV_1555,SPV_1560 | nadD,yqeK,unknown,yqeH,corA2,unknown,unknown | -2.847487077 | -1.414607182 | 1.446409346 | 1 |
| SPV_1562,SPV_1557,SPV_1556,SPV_1558,SPV_1559,SPV_1561,SPV_1555,SPV_1560 | unknown,nadD,yqeK,unknown,yqeH,corA2,unknown,unknown | 0.056423598 | -0.034750248 | -0.078751186 | 1 |
| SPV_1564 | unknown | -0.042932209 | -0.032061588 | 0.022865228 | 1 |
| SPV_1565 | unknown | 0.009926172 | 0.052014176 | 0.056302623 | 1 |
| SPV_1567,SPV_1566 | trxA,unknown | 0.210192091 | 0.029859923 | -0.16670189 | 1 |
| SPV_1568 | queF | 0.151908507 | 0.041784421 | -0.094415887 | 1 |
| SPV_1569 | aqpZ | 0.133757747 | 0.078407138 | -0.04135668 | 1 |
| SPV_1570 | unknown | 0.075743198 | 0.022250212 | -0.042411308 | 1 |
| SPV_1571 | pepF2 | 0.233386665 | -0.008124085 | -0.22903383 | 1 |
| SPV_1572 | unknown | 0.121743285 | -0.006063798 | -0.113261525 | 1 |
| SPV_1573 | prmA | 0.137921681 | 0.005750657 | -0.116764794 | 1 |
| SPV_1574 | unknown | 0.115185702 | 0.084084011 | -0.012980971 | 1 |
| SPV_1575 | unknown | -0.089119195 | 0.110374673 | 0.213398825 | 1 |
| SPV_1576 | unknown | 0.141440069 | 0.02213685 | -0.105131439 | 1 |
| SPV_1577 | hicB | 0.136055942 | 0.005712094 | -0.107929876 | 1 |
| SPV_1578,SPV_1579 | unknown,unknown | 0.006626991 | 0.128044469 | 0.131163684 | 1 |
| SPV_1580,SPV_2392,SPV_1581 | unknown,ssrS,tRNA-Lys-1 | -0.025786322 | 0.019180076 | 0.059192173 | 1 |
| SPV_1583,SPV_1582,SPV_1585,SPV_1584 | unknown,sacA,unknown,unknown | 0.110719352 | 0.037751698 | -0.059589938 | 1 |
| SPV_1586 | unknown | 0.026589798 | 0.003731061 | -0.008700759 | 1 |
| SPV_1587 | mga2 | -0.100200621 | 0.049982751 | 0.17028039 | 1 |
| SPV_1590,SPV_2393,SPV_1591,SPV_1589,SPV_1588 | gls24,unknown,unknown,unknown,unknown | 0.140388228 | -0.04741203 | -0.169278946 | 1 |
| SPV_1592 | unknown | 0.221152231 | 0.014336779 | -0.195642399 | 1 |
| SPV_1593 | cclA | 0.17651276 | 0.037494667 | -0.120218557 | 1 |
| SPV_1595,SPV_1594 | unknown,unknown | -2.009011107 | -1.341546259 | 0.680943127 | 1 |
| SPV_1597,SPV_1598,SPV_1600,SPV_1596,SPV_1602,SPV_1599,SPV_1601 | trpB,trpF,trpD,trpA,trpE,trpC,trpG | 0.066817527 | 0.095713704 | 0.044214766 | 1 |
| SPV_1603 | unknown | 0.216854262 | 0.062963253 | -0.134659134 | 1 |
| SPV_1605 | unknown | 0.193172732 | -0.107705631 | -0.283746505 | 1 |
| SPV_1609,SPV_1610,SPV_1608,SPV_1606,SPV_1607 | unknown,unknown,unknown,mgtC,sfuB | -0.061054927 | 0.07768169 | 0.149967352 | 1 |
| SPV_1609,SPV_1617,SPV_2396,SPV_1613,SPV_1610,SPV_1608,SPV_1606,SPV_1614,SPV_1612,SPV_1607 | unknown,pfbA,unknown,galT-1,unknown,unknown,mgtC,phoU3,galE-2,sfuB | 0.119322755 | 0.068825083 | -0.038439548 | 1 |
| SPV_1618 | unknown | 0.010589687 | 0.025932697 | 0.02988369 | 1 |
| SPV_1620,SPV_1619 | unknown,aatB | -0.770173209 | 0.13845641 | 0.926503154 | 1 |
| SPV_1621 | unknown | 0.115632546 | -0.003012277 | -0.106856477 | 1 |

| | | | | | |
|---|---|---|---|---|---|
| SPV_1622 | unknown | -0.013181502 | 0.156914568 | 0.180731513 | 1 |
| SPV_1626 | exoA | 0.305531213 | -0.063307192 | -0.354980209 | 1 |
| SPV_1627 | unknown | -0.017309146 | 0.024305318 | 0.058097708 | 1 |
| SPV_1630,SPV_1631 | dpnD,dpnC | 0.205551428 | 0.117724994 | -0.080116236 | 1 |
| SPV_1632 | paaI | -0.064095353 | 0.095519394 | 0.170408762 | 1 |
| SPV_1633 | galT-2 | -0.040712012 | 0.019767161 | 0.081457118 | 1 |
| SPV_1634 | galK | 0.343452051 | 0.022295426 | -0.311996517 | 1 |
| SPV_1635 | galR | 0.108083946 | 0.034146993 | -0.060658818 | 1 |
| SPV_1637,SPV_1636 | nmlR,adhB | -0.127730281 | -0.046965244 | 0.099297801 | 1 |
| SPV_1637,SPV_1636,SPV_1638 | nmlR,adhB,czcD | 0.203006131 | 0.008598115 | -0.182317673 | 1 |
| SPV_1640 | pnuC | 0.315373757 | -0.006943553 | -0.305562461 | 1 |
| SPV_1643,SPV_1644,SPV_1645,SPV_1642 | proV,unknown,unknown,proWX | -0.092772236 | 0.088031665 | 0.19432675 | 1 |
| SPV_1647,SPV_1646 | pepA,unknown | 0.206375483 | 0.097098594 | -0.090784792 | 1 |
| SPV_1648 | unknown | 0.185319812 | -0.075275539 | -0.24791126 | 1 |
| SPV_1649,SPV_1652,SPV_1650,SPV_1651 | piuB,piuA,piuC,piuD | 0.158536254 | 0.034757698 | -0.110283761 | 1 |
| SPV_1653 | yidD | 0.207126375 | 0.179416825 | -0.011142814 | 1 |
| SPV_1654 | rluB | 0.002875658 | -0.001313248 | 0.006196317 | 1 |
| SPV_1655 | scpB | -0.099147723 | -0.247947919 | -0.134826942 | 1 |
| SPV_1656 | scpA | -0.213668396 | -0.21968383 | 0.006449589 | 1 |
| SPV_1657 | xerD | -0.009044258 | -0.157587576 | -0.135774594 | 1 |
| SPV_1658,SPV_1659,SPV_1661,SPV_1660 | unknown,unknown,murI,rdgB | -0.642502638 | 0.383907394 | 1.041749316 | 1 |
| SPV_1662 | unknown | -0.22710028 | 0.531062594 | 0.771889608 | 1 |
| SPV_1663 | treC | 0.382038522 | -0.025014891 | -0.390008065 | 1 |
| SPV_1664 | treP | -0.063609424 | 0.056099032 | 0.132807289 | 1 |
| SPV_1665 | treR | 0.063345857 | 0.025144717 | -0.028574052 | 1 |
| SPV_1667 | amiF | -0.857521791 | -1.056106693 | -0.183708502 | 1 |
| SPV_1668 | amiE | -1.092172178 | -1.097679198 | 0.005654319 | 1 |
| SPV_1669 | amiD | -0.911115778 | -1.096897057 | -0.171816548 | 1 |
| SPV_1670 | amiC | -1.234878949 | -0.969436268 | 0.28015427 | 1 |
| SPV_1671 | amiA | -1.360074328 | -0.808706813 | 0.567636945 | 1 |
| SPV_1672 | tacL | -0.096939642 | 0.878487842 | 0.989543526 | 1 |
| SPV_1673 | gtfA | 0.050892878 | 0.033636878 | -0.009045065 | 1 |
| SPV_1674 | unknown | -0.446938019 | -0.513017834 | -0.05122259 | 1 |
| SPV_1675 | msmG | 0.030565435 | 0.125276121 | 0.11302414 | 1 |
| SPV_1676 | msmF | -0.06987134 | 0.028304332 | 0.110948947 | 1 |
| SPV_1677 | msmE | 0.096138156 | 0.087011032 | 0.007294074 | 1 |
| SPV_1678 | agaN | 0.195084369 | -0.112858197 | -0.305775415 | 1 |
| SPV_1679,SPV_1680 | msmR,birA | 0.005395659 | 0.114120553 | 0.125465631 | 1 |
| SPV_1682 | tRNA-Ser-2 | -2.889568098 | -1.81952245 | 1.081837078 | 1 |
| SPV_1683,SPV_1682 | tRNA-Ile-1,tRNA-Ser-2 | -3.655017667 | -2.315606359 | 1.332324036 | 1 |

| | | | | | |
|---|---|---|---|---|---|
| SPV_1697 | tRNA-Asp-1 | -3.777329606 | -2.592657037 | 1.195308616 | 1 |
| SPV_1698 | tRNA-Val-1 | -0.263425458 | -0.068713462 | 0.208502321 | 1 |
| SPV_1703 | tRNA-Glu-2 | -3.986884544 | -3.182174977 | 0.824756262 | 1 |
| SPV_1704 | rumA-2 | 0.190744502 | 0.048327478 | -0.132297645 | 1 |
| SPV_1705 | recX | 0.125140422 | -0.007930076 | -0.118349727 | 1 |
| SPV_1706 | unknown | -0.307516454 | 0.548348514 | 0.868480414 | 1 |
| SPV_1707 | unknown | 0.076754786 | 0.121135307 | 0.056816087 | 1 |
| SPV_1708 | unknown | 0.191370735 | 0.02425698 | -0.154987808 | 1 |
| SPV_1710,SPV_1709 | groES,groEL | -0.023843075 | -0.070897128 | -0.031233781 | 1 |
| SPV_1711 | ssbB | 0.039598767 | 0.088318242 | 0.063690589 | 1 |
| SPV_1712 | ydfG | 0.105595268 | 0.033897784 | -0.05920365 | 1 |
| SPV_1713 | unknown | -0.012434619 | -0.126649301 | -0.099367602 | 1 |
| SPV_1714 | unknown | 0.025863345 | 0.082425974 | 0.07332964 | 1 |
| SPV_1715 | unknown | 0.150356773 | 0.139688651 | 0.007317134 | 1 |
| SPV_1717,SPV_1718,SPV_1716 | unknown,unknown,unknown | -0.250113053 | 0.02584713 | 0.282374103 | 1 |
| SPV_1719,SPV_1720 | unknown,unknown | 0.202799422 | 0.042313084 | -0.146292354 | 1 |
| SPV_1725 | yeeN | 0.006649571 | -0.019446165 | -0.011501706 | 1 |
| SPV_1727,SPV_1729,SPV_1726,SPV_1728 | unknown,unknown,ply,unknown | 0.225360832 | -0.046712663 | -0.255553021 | 1 |
| SPV_1731 | unknown | -0.013438087 | 0.104272184 | 0.133606435 | 1 |
| SPV_1732 | unknown | -0.112135741 | 0.08987772 | 0.208814427 | 1 |
| SPV_1736 | unknown | 0.168018359 | 0.078288038 | -0.078348548 | 1 |
| SPV_1737 | lytA | -0.493059232 | -0.089429645 | 0.415414836 | 1 |
| SPV_1738,SPV_1737 | dinF,lytA | -0.066393458 | -0.028423244 | 0.053989775 | 1 |
| SPV_1738,SPV_1739,SPV_1737 | dinF,recA,lytA | -0.105280529 | 0.067128776 | 0.183087051 | 1 |
| SPV_1738,SPV_1739,SPV_1740,SPV_1737 | dinF,recA,cinA,lytA | 0.028027266 | 0.224299313 | 0.211432303 | 1 |
| SPV_1743,SPV_1741,SPV_1742 | tsaE,lytR,unknown | -1.092787387 | 0.080110061 | 1.19142338 | 1 |
| SPV_1743,SPV_1744,SPV_1741,SPV_1742 | tsaE,comM,lytR,unknown | -0.11999913 | 0.083106371 | 0.217117955 | 1 |
| SPV_1745 | plcR | 0.073315076 | -0.047988588 | -0.10864711 | 1 |
| SPV_1751,SPV_1755,SPV_1752,SPV_1754,SPV_1750,SPV_1746,SPV_1747,SPV_1748,SPV_2420,SPV_1749,SPV_1753 | unknown,unknown,clyB,unknown,wrbA,unknown,pneA1,pneA2,unknown,lanM,unknown | 0.066898682 | 0.165443585 | 0.110510434 | 1 |
| SPV_1757 | ndk | 0.398987715 | -0.153951569 | -0.534446641 | 1 |
| SPV_1758 | rpoC | -2.310985889 | -1.246688286 | 1.084373408 | 1 |
| SPV_1759 | rpoB | -2.514447822 | -1.511412401 | 1.017110545 | 1 |
| SPV_1760 | tRNA-Cys-1 | -2.560825977 | -1.890309725 | 0.68323523 | 1 |
| SPV_1761 | hlyX | 0.147716709 | -0.035274043 | -0.169194407 | 1 |
| SPV_1762 | endA | -0.003063581 | 0.119164237 | 0.13854387 | 1 |
| SPV_1763 | epuA | -0.011905689 | 0.027267622 | 0.056560521 | 1 |
| SPV_1764 | murA-2 | -0.161900838 | 0.01207357 | 0.186165307 | 1 |

| | | | | | |
|---|---|---|---|---|---|
| SPV_1765,SPV_1767,SPV_1766 | unknown,unknown,coaD | -0.027797581 | 0.044247104 | 0.086161932 | 1 |
| SPV_1768 | asnA | -0.006787452 | 0.112527156 | 0.131247279 | 1 |
| SPV_1769 | unknown | 0.032874713 | 0.019018134 | -0.00197871 | 1 |
| SPV_1771 | yjfA | 0.151258348 | 0.047482482 | -0.086379921 | 1 |
| SPV_1772 | acyP | 0.279474301 | 0.025562984 | -0.237916639 | 1 |
| SPV_1773 | yidC1 | -0.319623069 | 0.262328867 | 0.594602993 | 1 |
| SPV_1774 | pflA | -0.06351138 | 0.162229253 | 0.242482198 | 1 |
| SPV_1775 | lysA | 0.352132335 | 0.111044776 | -0.218132343 | 1 |
| SPV_1776 | purR | 0.023033949 | -0.31366057 | -0.320218602 | 1 |
| SPV_1778,SPV_1777 | rmuC,yhaM | -0.05832999 | 0.097585367 | 0.16661121 | 1 |
| SPV_1780,SPV_1782,SPV_1779,SPV_1781 | rpe,ksgA,unknown,rsgA | -0.818602606 | -0.740973821 | 0.095693306 | 1 |
| SPV_1785,SPV_1784,SPV_1783 | unknown,unknown,unknown | 0.127118332 | 0.052835902 | -0.056313824 | 1 |
| SPV_1786 | unknown | 0.161183571 | 0.029848002 | -0.119030916 | 1 |
| SPV_1787,SPV_1788 | rnmV,tatD | 0.348814802 | -0.091596498 | -0.428165122 | 1 |
| SPV_1789 | diiA | -0.069369401 | 0.014627056 | 0.094791879 | 1 |
| SPV_1792 | unknown | -0.02352423 | 0.112165053 | 0.151300196 | 1 |
| SPV_1793 | unknown | 0.111042219 | -0.038413675 | -0.135568078 | 1 |
| SPV_1794 | unknown | 0.184111947 | 0.01475936 | -0.154741678 | 1 |
| SPV_1795 | unknown | 0.216855541 | 0.121060524 | -0.082239279 | 1 |
| SPV_1796 | unknown | -0.156381779 | 0.112072916 | 0.284938555 | 1 |
| SPV_1797 | ccpA | -0.44299673 | -0.205938647 | 0.253321113 | 1 |
| SPV_1803,SPV_1800,SPV_1801,SPV_1799,SPV_1798,SPV_1802 | unknown,unknown,unknown,unknown,unknown,unknown | 0.068793728 | 0.073651783 | 0.016732943 | 1 |
| SPV_1819 | nusG | -0.848421336 | -0.632266369 | 0.228358118 | 1 |
| SPV_1820 | secE | -1.438888548 | -0.923988155 | 0.535695139 | 1 |
| SPV_1821 | pbp2a | -0.196539934 | 0.203938747 | 0.416150568 | 1 |
| SPV_1822 | rluD2 | -0.07502001 | 0.03713834 | 0.125631974 | 1 |
| SPV_1823 | gap | -2.025384649 | -1.424644236 | 0.614880964 | 1 |
| SPV_1824 | unknown | 0.192640165 | 0.0611119 | -0.118197805 | 1 |
| SPV_1826 | nadC | -0.009666471 | 0.094991926 | 0.118915476 | 1 |
| SPV_1827,SPV_2426 | unknown,unknown | 0.100114491 | 0.008940547 | -0.077617817 | 1 |
| SPV_1829 | bguR | 0.085203533 | -0.156747983 | -0.231953284 | 1 |
| SPV_1833,SPV_1831,SPV_1830,SPV_1832 | bguC,bguD,bguA,bguB | 0.136507886 | -0.037866028 | -0.154728945 | 1 |
| SPV_1834 | adhE | -0.072088281 | 0.030144144 | 0.115600498 | 1 |
| SPV_1836 | unknown | 0.237628704 | 0.085815549 | -0.139482515 | 1 |
| SPV_1837 | unknown | -0.03125637 | 0.1285974 | 0.174233505 | 1 |
| SPV_1838 | yajC | 0.019855364 | 0.118690506 | 0.11297017 | 1 |
| SPV_1839,SPV_1838 | ulaH,yajC | -0.199243558 | -0.32278565 | -0.10394674 | 1 |
| SPV_1840,SPV_1841 | ulaG,ulaR | -0.09552784 | -0.039069885 | 0.071282187 | 1 |
| SPV_1842 | ulaF | 0.053916567 | -0.082809683 | -0.122406516 | 1 |
| SPV_1843 | ulaE | 0.211440513 | 0.052667633 | -0.143445972 | 1 |

| | | | | | |
|---|---|---|---|---|---|
| SPV_1844 | ulaD | 0.030111811 | 0.140053159 | 0.123599014 | 1 |
| SPV_1845 | ulaC | 0.096585478 | 0.137434229 | 0.055279584 | 1 |
| SPV_1846 | ulaB | 0.071951507 | 0.194510987 | 0.139021311 | 1 |
| SPV_1847 | ulaA | -0.100538481 | 0.066228765 | 0.178899339 | 1 |
| SPV_1848 | unknown | 0.184073696 | -0.011262675 | -0.179309639 | 1 |
| SPV_1849 | eloR | -0.18090323 | -0.979817109 | -0.784981794 | 1 |
| SPV_1850 | yidC2 | -0.175365005 | -1.417469701 | -1.227216045 | 1 |
| SPV_1851 | rnpA | -2.285559681 | -2.045538271 | 0.247576415 | 1 |
| SPV_1852 | unknown | 0.063539169 | -0.103102488 | -0.151210911 | 1 |
| SPV_1853 | ackA | -0.00619002 | -0.382672715 | -0.362420653 | 1 |
| SPV_1854 | unknown | -0.013852327 | 0.047184975 | 0.07559014 | 1 |
| SPV_1863,SPV_1861,SPV_1858,SPV_1859,SPV_1862,SPV_2427,SPV_1857,SPV_1860 | comGA,comGC,comGF,comGE,comGB,unknown,comGG,comGD | 0.186284083 | -0.060611586 | -0.229565812 | 1 |
| SPV_1864 | unknown | 0.278335516 | 0.089842693 | -0.172839194 | 1 |
| SPV_1865 | adh | 0.180400406 | 0.045724683 | -0.122108172 | 1 |
| SPV_1866 | nagA | 0.33979125 | -0.005429671 | -0.33313923 | 1 |
| SPV_1867 | adr | 0.152462105 | 0.037150706 | -0.101815669 | 1 |
| SPV_1868 | tgt | -0.084728022 | 0.083946202 | 0.182645031 | 1 |
| SPV_1869 | unknown | -0.007820299 | 0.127358432 | 0.150927162 | 1 |
| SPV_1870 | pcp2 | 0.429824572 | 0.136285752 | -0.275041914 | 1 |
| SPV_1871,SPV_1872,SPV_1870 | unknown,unknown,pcp2 | 0.098962832 | 0.070235391 | -0.016185194 | 1 |
| SPV_1873 | unknown | -0.077739616 | 0.0976073 | 0.186695494 | 1 |
| SPV_1874 | unknown | 0.116820851 | 0.074697611 | -0.021240779 | 1 |
| SPV_1875 | unknown | 0.548950501 | -0.04259044 | -0.567345872 | 1 |
| SPV_1876 | unknown | -0.012085443 | 0.087633174 | 0.113492823 | 1 |
| SPV_1877 | thrC | 0.19001445 | 0.113440218 | -0.068660468 | 1 |
| SPV_1878 | unknown | 0.228099775 | 0.015816567 | -0.195773514 | 1 |
| SPV_1879 | tRNA-Leu-6 | -1.166768799 | -0.885555764 | 0.295125255 | 1 |
| SPV_1880,SPV_1879,SPV_1881 | tRNA-Gln-2,tRNA-Leu-6,tRNA-His-1 | -3.053860317 | -2.422148704 | 0.645052811 | 1 |
| SPV_1882 | tRNA-Trp-1 | -0.798637287 | -0.058236335 | 0.756790844 | 1 |
| SPV_1883 | tRNA-Tyr-2 | -2.36869296 | -1.367616575 | 1.022318152 | 1 |
| SPV_1884 | tRNA-Phe-2 | -0.064434604 | 0.00720358 | 0.089837485 | 1 |
| SPV_1888 | tRNA-Gly-4 | -3.790641164 | -2.515301854 | 1.286711688 | 1 |
| SPV_1895 | unknown | -2.189379336 | -1.211157661 | 0.997667747 | 1 |
| SPV_1896 | gltX | -2.784453485 | -1.700513551 | 1.105585356 | 1 |
| SPV_1897 | pgi | -2.57631199 | -1.817397169 | 0.777934732 | 1 |
| SPV_1899,SPV_1898 | unknown,unknown | -0.156065318 | 0.077803942 | 0.247394461 | 1 |
| SPV_1900 | patB | 0.109352166 | -0.001221039 | -0.091554567 | 1 |
| SPV_1903 | hexA | 0.135002127 | 0.0633337 | -0.054458354 | 1 |
| SPV_1904 | argR1 | -2.276298397 | -0.971252806 | 1.313557444 | 1 |
| SPV_1905 | argS | -0.440116127 | -0.173460951 | 0.278960615 | 1 |
| SPV_1906 | unknown | -0.020287283 | 0.037314008 | 0.072511071 | 1 |

| | | | | | |
|---|---|---|---|---|---|
| SPV_1907 | unknown | 0.076976343 | -0.097976219 | -0.157789398 | 1 |
| SPV_1909,SPV_1908 | pnpS,pnpR | 0.075101902 | 0.024501667 | -0.037292568 | 1 |
| SPV_1910 | pstS1 | 0.044027181 | 0.031686128 | 0.002954247 | 1 |
| SPV_1911 | pstC1 | 0.061781528 | 0.179323758 | 0.131884355 | 1 |
| SPV_1912 | pstA1 | -0.001884553 | -0.043229505 | -0.028497241 | 1 |
| SPV_1913 | pstB1 | 0.183097731 | 0.100209441 | -0.063788852 | 1 |
| SPV_1914 | phoU1 | 0.291475268 | 0.023893402 | -0.254877439 | 1 |
| SPV_1915 | unknown | 0.293841187 | 0.100652244 | -0.174922807 | 1 |
| SPV_1916 | unknown | -0.060588307 | -0.094015797 | -0.022414733 | 1 |
| SPV_1917 | unknown | -0.434699438 | -0.230170721 | 0.219088411 | 1 |
| SPV_1918,SPV_1919 | gpsA,galU | -0.19177065 | -0.33597876 | -0.130133642 | 1 |
| SPV_1920 | unknown | -0.017196799 | 0.094599459 | 0.130083227 | 1 |
| SPV_1921 | unknown | 0.033578262 | 0.04749633 | 0.027848632 | 1 |
| SPV_1922 | hipO | -1.110297239 | -0.453887124 | 0.667321507 | 1 |
| SPV_1923 | dapD | -0.027378425 | 0.107757337 | 0.156995558 | 1 |
| SPV_1924 | unknown | 0.061178397 | 0.123059392 | 0.076393589 | 1 |
| SPV_1925 | pbp1b | 0.074731689 | -0.163771066 | -0.223792251 | 1 |
| SPV_1926 | tyrS | 0.06535133 | 0.03194983 | -0.021613681 | 1 |
| SPV_1927,SPV_1928 | ctpC,unknown | 0.066377421 | 0.091996854 | 0.039957979 | 1 |
| SPV_1929 | rrmA | -0.089953961 | -0.014044408 | 0.084594832 | 1 |
| SPV_1930 | unknown | 0.167058056 | 0.05003242 | -0.102103887 | 1 |
| SPV_1931 | unknown | 0.119934768 | 0.100270902 | -0.003540332 | 1 |
| SPV_1933,SPV_1932 | malQ,malP | 0.063935651 | -0.00866111 | -0.059704807 | 1 |
| SPV_1934 | malX | -0.026960107 | 0.030987614 | 0.078427826 | 1 |
| SPV_1935 | malC | 0.094731544 | 0.026443947 | -0.061699785 | 1 |
| SPV_1936 | malD | 0.048855429 | -0.020077722 | -0.053328208 | 1 |
| SPV_1938,SPV_1937 | malR,malA | -0.183438031 | -0.012122137 | 0.186592759 | 1 |
| SPV_1943 | unknown | -0.020368932 | 0.014583638 | 0.047743809 | 1 |
| SPV_1944 | unknown | 0.131312261 | -0.039961006 | -0.156754008 | 1 |
| SPV_1945 | unknown | 0.137647703 | -0.024114289 | -0.137432924 | 1 |
| SPV_1946 | unknown | 0.101308716 | 0.042328577 | -0.036416492 | 1 |
| SPV_1947 | unknown | -0.192809034 | 0.146248478 | 0.35084116 | 1 |
| SPV_1948 | unknown | -0.188952301 | 0.044290131 | 0.245572233 | 1 |
| SPV_1950 | hisS | -2.076383324 | -1.242660661 | 0.846642636 | 1 |
| SPV_1951 | unknown | 0.077912987 | 0.06613759 | 0.003252954 | 1 |
| SPV_1952 | rgg | 0.166383307 | -0.010507889 | -0.152943977 | 1 |
| SPV_1953 | unknown | -0.114528134 | -0.036808351 | 0.089906842 | 1 |
| SPV_1954 | unknown | 0.003407248 | 0.039053346 | 0.047785513 | 1 |
| SPV_1955 | unknown | 0.394456105 | 0.115864943 | -0.261008147 | 1 |
| SPV_1956 | ilvD | 0.176678366 | 0.057930714 | -0.105445034 | 1 |
| SPV_1957 | tktC | 0.299355251 | 0.145943895 | -0.137886767 | 1 |
| SPV_1958 | tktN | 0.015255741 | 0.029846022 | 0.029410234 | 1 |
| SPV_1959 | ulaA2 | -0.061219244 | 0.031274686 | 0.105541317 | 1 |

| | | | | | |
|---|---|---|---|---|---|
| SPV_1960 | ulaB2 | 0.074491657 | 0.044002614 | -0.016418067 | 1 |
| SPV_1961 | ulaR2 | 0.222552311 | 0.097403492 | -0.106975859 | 1 |
| SPV_1962,SPV_2438 | unknown,unknown | 0.097407458 | 0.095897992 | 0.012664129 | 1 |
| SPV_1963,SPV_1964 | rpmF,rpmG3 | -2.479692227 | -1.388056395 | 1.105535764 | 1 |
| SPV_1965 | cbpN | 0.409556387 | -0.007008061 | -0.393796002 | 1 |
| SPV_1972,SPV_1970,SPV_1969,SPV_2439,SPV_1971 | unknown,unknown,unknown,unknown,unknown | 0.249051528 | 0.029020054 | -0.203664689 | 1 |
| SPV_1973 | unknown | 0.144048519 | 0.079861384 | -0.048861773 | 1 |
| SPV_1975,SPV_2442 | arcA,srf-26 | -0.038650551 | 0.094460423 | 0.14280076 | 1 |
| SPV_1976 | arcB | 0.133933526 | 0.048105294 | -0.079086514 | 1 |
| SPV_1977 | arcC | 0.211956401 | 0.053571038 | -0.13970937 | 1 |
| SPV_1978 | arcD | -0.094140846 | 0.043250133 | 0.167429115 | 1 |
| SPV_1979 | unknown | 0.0273161 | 0.073905153 | 0.061316213 | 1 |
| SPV_1981 | unknown | 0.146084321 | 0.132162737 | -5.17909E-05 | 1 |
| SPV_1983 | unknown | 0.048466528 | -0.027243368 | -0.066031572 | 1 |
| SPV_1984 | ybbK | 0.099322572 | -0.000110212 | -0.08169501 | 1 |
| SPV_1988,SPV_1992,SPV_1987,SPV_1989,SPV_1986,SPV_1990,SPV_1995,SPV_1994,SPV_1993,SPV_1985,SPV_1991 | fucY,unknown,fucL,unknown,fucI,unknown,fucK,fucA,fucU,adh2,unknown | 0.099927992 | 0.032444961 | -0.057199962 | 1 |
| SPV_1996 | fucR | 0.010394129 | 0.052914347 | 0.055031315 | 1 |
| SPV_1997,SPV_1999,SPV_2000,SPV_1998 | adcA,adcC,adcR,adcB | -0.299314317 | -0.692011378 | -0.379206356 | 1 |
| SPV_2002 | dltD | 0.154325993 | 0.028406449 | -0.110704906 | 1 |
| SPV_2003 | dltC | 0.0403476 | 0.020626747 | -0.005172476 | 1 |
| SPV_2004 | dltB | -0.031267339 | 0.123325317 | 0.166633312 | 1 |
| SPV_2005 | dltA | 0.045491928 | 0.009367674 | -0.027698402 | 1 |
| SPV_2006 | unknown | 0.332427201 | -0.021319507 | -0.331565286 | 1 |
| SPV_2007 | unknown | -0.026876921 | 0.004903593 | 0.045844675 | 1 |
| SPV_2008 | unknown | -0.138399333 | 0.051049999 | 0.202526637 | 1 |
| SPV_2009 | unknown | -0.051938293 | 0.121410453 | 0.18312747 | 1 |
| SPV_2010,SPV_2011,SPV_2012,SPV_2013 | unknown,glpF,glpO,glpK | -0.053046686 | -0.01772556 | 0.04676379 | 1 |
| SPV_2014,SPV_2010,SPV_2011,SPV_2012,SPV_2013 | unknown,unknown,glpF,glpO,glpK | -0.010833455 | 0.150486758 | 0.173695549 | 1 |
| SPV_2015,SPV_2016 | hslO,unknown | 0.002391968 | -0.277822314 | -0.264150431 | 1 |
| SPV_2018,SPV_2017 | unknown,cbpA | -0.041361211 | 0.074822891 | 0.126250215 | 1 |
| SPV_2019 | unknown | -0.01379196 | -0.048176785 | -0.019106183 | 1 |
| SPV_2020 | unknown | 0.000666734 | 0.064614938 | 0.079782619 | 1 |
| SPV_2021 | unknown | 0.358048448 | -0.145298505 | -0.484392555 | 1 |
| SPV_2023,SPV_2022 | ctsR,clpC | -0.067243749 | -0.141977294 | -0.057496734 | 1 |
| SPV_2024,SPV_2026,SPV_2027,SPV_2025 | thiZ,thiX,unknown,thiY | 0.08260542 | -0.019702074 | -0.092138134 | 1 |

| | | | | | |
|---|---|---|---|---|---|
| SPV_2029 | unknown | -0.936751128 | 0.114340162 | 1.065189226 | 1 |
| SPV_2030 | dnaC | -2.57752606 | -1.441032291 | 1.156828266 | 1 |
| SPV_2031,SPV_2032 | rplI,pde1 | -3.095414856 | -1.894748661 | 1.216546354 | 1 |
| SPV_2033 | hpf | -0.138422622 | 0.033445597 | 0.186340771 | 1 |
| SPV_2034,SPV_2035 | comFC,comFA | 0.267286018 | 0.076726485 | -0.176092325 | 1 |
| SPV_2036 | unknown | -0.154659722 | 0.010052506 | 0.175593341 | 1 |
| SPV_2037 | cysK | -0.017949707 | 0.080926278 | 0.116739776 | 1 |
| SPV_2039 | unknown | -0.010664016 | 0.046539515 | 0.073820723 | 1 |
| SPV_2040 | unknown | 0.022330221 | 0.083048936 | 0.076372568 | 1 |
| SPV_2041 | tsf | -2.947425955 | -1.939900453 | 1.024711723 | 1 |
| SPV_2042,SPV_2458 | rpsB,srf-30 | -3.639200165 | -2.481218996 | 1.171855115 | 1 |
| SPV_2043 | pcsB | -5.035403672 | -3.389337264 | 1.656578228 | 1 |
| SPV_2044 | mreD | 0.197917598 | -0.022909556 | -0.206433748 | 1 |
| SPV_2045 | mreC | 0.226161719 | 0.13503528 | -0.079475042 | 1 |
| SPV_2046 | cbiQ | 0.316485009 | -0.079687515 | -0.378728719 | 1 |
| SPV_2047 | cbiO2 | 0.042655478 | -0.053201422 | -0.084257638 | 1 |
| SPV_2048 | cbiO1 | -0.050282867 | -0.238610617 | -0.185447883 | 1 |
| SPV_2050,SPV_2051,SPV_2052,SPV_2049 | rodZ,unknown,unknown,pgsA | -0.040125772 | -0.059076646 | -0.00475261 | 1 |
| SPV_2054,SPV_2053 | recF,unknown | -0.198500171 | -0.070769377 | 0.144760353 | 1 |
| SPV_2055 | guaB | 0.159606089 | 0.043249937 | -0.102192039 | 1 |
| SPV_2056 | trpS | -1.929105204 | -1.333847409 | 0.611462237 | 1 |
| SPV_2057 | unknown | -0.12205689 | 0.020922138 | 0.153338972 | 1 |
| SPV_2058 | unknown | 0.209180589 | 0.022802403 | -0.177947807 | 1 |
| SPV_2059 | unknown | 0.065041735 | 0.085131467 | 0.031529408 | 1 |
| SPV_2060 | pipR | 0.217117579 | 0.141899664 | -0.061622235 | 1 |
| SPV_2061 | tRNA-Asn-2 | -0.480570638 | -0.212732559 | 0.28422936 | 1 |
| SPV_2062,SPV_2064,SPV_2063,SPV_2065 | tRNA-Glu-5,comD,comE,comC1 | 0.161615929 | 0.07672631 | -0.069113498 | 1 |
| SPV_2067,SPV_2066 | rlmH,tRNA-Arg-5 | 0.184136772 | 0.010262426 | -0.155597726 | 1 |
| SPV_2068,SPV_2069 | htrA,parB | 0.040141353 | 0.064624515 | 0.041047716 | 1 |
| SPV_2069 | parB | 0.110990454 | 0.097254329 | -0.000588894 | 1 |
| SPV_2070 | unknown | -0.225904266 | 0.042441702 | 0.285418481 | 1 |
| SPV_2073 | unknown | 0.020695533 | -0.037887103 | -0.040035265 | 1 |
| SPV_2076 | unknown | 0.064125763 | 0.037726315 | -0.012731168 | 1 |
| SPV_2078 | ccnC | -0.15048584 | 0.284809037 | 0.442132109 | 1 |
| SPV_2078,SPV_0024 | ccnC,purA | 0.171328572 | 0.05361714 | -0.104395132 | 1 |
| SPV_2078,SPV_0024,SPV_0023 | ccnC,purA,comW | -0.067835435 | 0.062490951 | 0.145562459 | 1 |
| SPV_2081 | srf-01 | 0.097934541 | -0.046127238 | -0.126029504 | 1 |
| SPV_2082 | unknown | -0.082223294 | -0.044305985 | 0.0533539 | 1 |
| SPV_2084 | srf-02 | 0.14185617 | 0.063364389 | -0.063914538 | 1 |
| SPV_2085 | unknown | -0.094439941 | 0.099756922 | 0.211555811 | 1 |
| SPV_2086,SPV_0049,SPV_0050 | srf-03,comA,comB | 0.060140917 | -0.022765212 | -0.061713611 | 1 |

| | | | | | |
|---|---|---|---|---|---|
| SPV_2087 | unknown | -0.030759629 | 0.125360663 | 0.169102491 | 1 |
| SPV_2088 | unknown | 0.094421699 | 0.010909106 | -0.066473752 | 1 |
| SPV_2089 | unknown | 0.098781052 | 0.080003352 | -0.005147283 | 1 |
| SPV_2090 | unknown | 0.016790843 | 0.005712573 | -0.002121412 | 1 |
| SPV_2091 | unknown | 0.095215375 | 0.053333199 | -0.023628626 | 1 |
| SPV_2092 | unknown | 0.25479312 | 0.067689281 | -0.179520991 | 1 |
| SPV_2096 | unknown | 0.088979795 | -0.01189518 | -0.083299105 | 1 |
| SPV_2100 | unknown | 0.323245348 | 0.024470086 | -0.285366009 | 1 |
| SPV_2102,SPV_0120,SPV_0123,SPV_0119,SPV_2103,SPV_0122,SPV_0121,SPV_0124 | unknown,unknown,unknown,unknown,unknown,unknown,unknown,unknown | 0.055789739 | 0.02536023 | -0.013556703 | 1 |
| SPV_2104 | unknown | 0.113926827 | 0.033581286 | -0.065593627 | 1 |
| SPV_2109,SPV_0132 | cibC,cibB | -0.017383639 | 0.048628621 | 0.079616422 | 1 |
| SPV_2109,SPV_0132,SPV_0133 | cibC,cibB,cibA | -0.029591029 | 0.035551937 | 0.077129931 | 1 |
| SPV_2110 | unknown | 0.075431082 | -0.049470581 | -0.1089447 | 1 |
| SPV_2113 | unknown | 0.010460509 | -0.010661178 | -0.003666381 | 1 |
| SPV_2115 | unknown | 0.365382492 | 0.018801915 | -0.328011377 | 1 |
| SPV_2116 | unknown | -0.010020995 | 0.205566373 | 0.228470118 | 1 |
| SPV_2117 | unknown | -0.231191257 | 0.034496882 | 0.276233019 | 1 |
| SPV_2119 | srf-05 | 0.216930329 | 0.012122711 | -0.193568796 | 1 |
| SPV_2120 | srf-06 | -0.211508299 | 0.088624618 | 0.312026401 | 1 |
| SPV_2121,SPV_0186 | unknown,unknown | 0.058980155 | -0.198724832 | -0.249391363 | 1 |
| SPV_2125 | ccnE | 0.195756379 | 0.144415246 | -0.035725134 | 1 |
| SPV_2126 | unknown | -0.017034451 | 0.064203087 | 0.095822014 | 1 |
| SPV_2127 | unknown | -0.131010389 | 0.107428874 | 0.25479672 | 1 |
| SPV_2129 | ccnA | 0.113790285 | 0.072063528 | -0.023962486 | 1 |
| SPV_2130 | ccnB | -0.091676801 | 0.097522408 | 0.202183183 | 1 |
| SPV_2131 | srf-07 | 0.180183756 | 0.025703935 | -0.138260539 | 1 |
| SPV_2132 | unknown | -0.006824709 | 0.043142067 | 0.060232134 | 1 |
| SPV_2133 | ccnD | 0.156062079 | 0.044341965 | -0.096967115 | 1 |
| SPV_2134 | unknown | -0.007585179 | 0.022504921 | 0.040807463 | 1 |
| SPV_2135 | unknown | 0.228007195 | 0.038171906 | -0.174751865 | 1 |
| SPV_2137 | unknown | 0.306416131 | 0.063041104 | -0.226758464 | 1 |
| SPV_2139 | srf-08 | -0.05177296 | 0.140594766 | 0.203960726 | 1 |
| SPV_2140,SPV_0288 | NA,unknown | 0.470233242 | 0.009833559 | -0.445833224 | 1 |
| SPV_2141 | unknown | -0.059832906 | 0.187400632 | 0.267870179 | 1 |
| SPV_2142 | unknown | 0.01728739 | 0.126543669 | 0.12165689 | 1 |
| SPV_2144 | unknown | -0.310894721 | -0.460962214 | -0.132440951 | 1 |
| SPV_2146 | unknown | -0.128722321 | -0.418711328 | -0.270579502 | 1 |
| SPV_2157,SPV_0391 | ydiL,briC | 0.126305622 | 0.031977556 | -0.075794741 | 1 |
| SPV_2158 | unknown | -0.019041729 | 0.126600037 | 0.161936182 | 1 |
| SPV_2160 | unknown | 0.111359982 | 0.147012537 | 0.04959592 | 1 |
| SPV_2163 | unknown | 0.080460531 | 0.099369452 | 0.035151897 | 1 |

| | | | | | |
|---|---|---|---|---|---|
| SPV_2164 | unknown | 0.053988903 | 0.161741481 | 0.121734246 | 1 |
| SPV_2165 | unknown | 0.125531269 | 0.039256449 | -0.070274107 | 1 |
| SPV_2166 | unknown | -0.254762378 | 0.292743909 | 0.620935884 | 1 |
| SPV_2167 | srf-09 | -0.301980657 | -0.116285396 | 0.204083311 | 1 |
| SPV_2168 | unknown | 0.029019768 | 0.015946798 | 0.001241593 | 1 |
| SPV_2169 | unknown | 0.117796764 | -0.008195297 | -0.118980213 | 1 |
| SPV_2178 | unknown | -0.113911603 | -0.08321116 | 0.041776182 | 1 |
| SPV_2185 | srf-10 | 0.147437289 | 0.027119317 | -0.110424154 | 1 |
| SPV_2185,SPV_0500 | srf-10,unknown | -0.0869623 | 0.080425135 | 0.183452552 | 1 |
| SPV_2188 | unknown | 0.325078284 | 0.016779564 | -0.298455672 | 1 |
| SPV_2189 | unknown | 0.122182948 | 0.041540863 | -0.06549424 | 1 |
| SPV_2192 | unknown | 0.135403721 | 0.064578042 | -0.05632848 | 1 |
| SPV_2193 | unknown | 0.250185751 | 0.140533604 | -0.094435261 | 1 |
| SPV_2200 | srf-11 | 0.13382513 | 0.053675813 | -0.07053999 | 1 |
| SPV_2201 | unknown | -0.114983722 | -0.01446033 | 0.112869944 | 1 |
| SPV_2202 | unknown | 0.207441156 | -0.005634276 | -0.197965866 | 1 |
| SPV_2203 | unknown | 0.089023901 | 0.008356068 | -0.062532283 | 1 |
| SPV_2205 | unknown | -0.152117024 | 0.051578467 | 0.218975688 | 1 |
| SPV_2213 | unknown | -0.320360682 | -0.242011449 | 0.091907088 | 1 |
| SPV_2218,SPV_0673 | unknown,unknown | -0.279565635 | 0.09509315 | 0.38346235 | 1 |
| SPV_2219 | unknown | -0.033954038 | 0.041289702 | 0.085734208 | 1 |
| SPV_2220 | unknown | -0.139513954 | -0.168107106 | -0.013887494 | 1 |
| SPV_2226 | srf-12 | 0.019675932 | 0.031036642 | 0.024916151 | 1 |
| SPV_2227 | unknown | 0.048594058 | 0.006957715 | -0.028343879 | 1 |
| SPV_2238,SPV_0781,SPV_0778 | unknown,unknown,unknown | 0.018959234 | 0.026212386 | 0.018995765 | 1 |
| SPV_2239 | unknown | -0.013094093 | -0.042838263 | -0.017536855 | 1 |
| SPV_2240 | unknown | 0.230594595 | 0.032507752 | -0.176558991 | 1 |
| SPV_2242 | unknown | 0.176659975 | 0.005922562 | -0.15849269 | 1 |
| SPV_2244 | unknown | 0.165276934 | 0.030980061 | -0.121325577 | 1 |
| SPV_2245 | unknown | 0.104784194 | 0.071658827 | -0.017548205 | 1 |
| SPV_2246 | unknown | 0.155326174 | 0.084947889 | -0.061552188 | 1 |
| SPV_2247 | srf-13 | 0.16417072 | -0.0516649 | -0.206961733 | 1 |
| SPV_2249,SPV_0817 | unknown,unknown | 0.202157815 | 0.030220913 | -0.157901255 | 1 |
| SPV_2250 | unknown | -0.19150273 | 0.061914447 | 0.267651699 | 1 |
| SPV_2251 | unknown | -0.067963898 | 0.080973016 | 0.161595695 | 1 |
| SPV_2257,SPV_0846,SPV_0844,SPV_2256,SPV_0843 | unknown,unknown,comEC,unknown,comEA | -0.007865085 | 0.046963544 | 0.054310483 | 1 |
| SPV_2257,SPV_0846,SPV_2256 | unknown,unknown,unknown | 0.034326397 | 0.078884368 | 0.063713099 | 1 |
| SPV_2258 | srf-14 | -3.710147217 | -2.609074779 | 1.115069226 | 1 |
| SPV_2259 | rpmG1 | -0.053004988 | -0.085547921 | -0.016465651 | 1 |
| SPV_2260 | unknown | 0.09863049 | 0.006260552 | -0.073738225 | 1 |
| SPV_2265,SPV_0888 | phtD,lmb | -0.069768477 | 0.131782884 | 0.214115433 | 1 |
| SPV_2266 | dhaM | 0.309491929 | 0.012617569 | -0.282188556 | 1 |

| | | | | | |
|---|---|---|---|---|---|
| SPV_2267 | unknown | -0.08881107 | 0.017831192 | 0.119553629 | 1 |
| SPV_2270 | srf-15 | -3.818852405 | -2.686776468 | 1.142699471 | 1 |
| SPV_2274 | unknown | 0.212047741 | -0.058051248 | -0.256211198 | 1 |
| SPV_2278 | unknown | 0.094319874 | 0.075773279 | -0.001797403 | 1 |
| SPV_2279 | unknown | 0.079081724 | 0.019634243 | -0.041310583 | 1 |
| SPV_2282 | shp | 0.138888984 | 0.139338782 | 0.01568429 | 1 |
| SPV_2283 | unknown | 0.22420056 | 0.060336824 | -0.149428232 | 1 |
| SPV_2285,SPV_0931,SPV_0930 | unknown,pezT,pezA | -0.494153067 | 0.315231973 | 0.82403531 | 1 |
| SPV_2287,SPV_2285,SPV_0931,SPV_2286,SPV_0930 | unknown,unknown,pezT,unknown,pezA | 0.372586896 | -0.01259337 | -0.366515599 | 1 |
| SPV_2287,SPV_2285,SPV_0931,SPV_2286,SPV_0930,SPV_0927 | unknown,unknown,pezT,unknown,pezA,nplT | 0.144914252 | 0.076978329 | -0.052056417 | 1 |
| SPV_2288,SPV_0918,SPV_0919,SPV_0917,SPV_0915,SPV_0916 | unknown,piaD,unknown,piaC,piaA,piaB | 0.218256794 | 0.092415947 | -0.115151728 | 1 |
| SPV_2289 | unknown | -0.033830752 | 0.038772616 | 0.086341789 | 1 |
| SPV_2290 | unknown | 0.004610359 | 0.069666651 | 0.080311115 | 1 |
| SPV_2291 | srf-16 | -0.144603354 | 0.037865264 | 0.193062886 | 1 |
| SPV_2292 | srf-17 | 0.117026676 | 0.042648495 | -0.065785117 | 1 |
| SPV_2293 | phtF | 0.183537287 | 0.056671244 | -0.110063035 | 1 |
| SPV_2297 | phtB | 0.307799065 | 0.025891966 | -0.269631412 | 1 |
| SPV_2298 | unknown | 0.069305059 | 0.024023755 | -0.030846936 | 1 |
| SPV_2299 | unknown | 0.09628622 | -0.00168156 | -0.08419694 | 1 |
| SPV_2304 | unknown | 0.201706929 | 0.018136333 | -0.18141644 | 1 |
| SPV_2309 | unknown | 0.003713791 | 0.07993682 | 0.08356692 | 1 |
| SPV_2310 | unknown | 0.228226827 | 0.013835148 | -0.200416318 | 1 |
| SPV_2311 | unknown | 0.092611757 | -0.040829105 | -0.120431021 | 1 |
| SPV_2314 | unknown | -0.113573437 | -0.06839446 | 0.059113885 | 1 |
| SPV_2315 | unknown | 0.358072164 | -0.030868899 | -0.367036267 | 1 |
| SPV_2317 | srf-19 | 0.113919777 | 0.157298248 | 0.058036102 | 1 |
| SPV_2318 | unknown | 0.41899069 | 0.068558791 | -0.339944674 | 1 |
| SPV_2319 | unknown | 0.241292186 | -0.015163711 | -0.242732207 | 1 |
| SPV_2320 | unknown | 0.016473929 | -0.015303189 | -0.017080681 | 1 |
| SPV_2325 | unknown | 0.375187525 | 0.004638153 | -0.354561416 | 1 |
| SPV_2326 | unknown | 0.278506431 | 0.014931514 | -0.248999566 | 1 |
| SPV_2332 | unknown | 0.267672219 | -0.087782413 | -0.341384662 | 1 |
| SPV_2333 | unknown | -0.170101137 | -0.067925463 | 0.115077285 | 1 |
| SPV_2337 | unknown | 0.12716204 | 0.0641765 | -0.047027527 | 1 |
| SPV_2340,SPV_1309,SPV_1308 | unknown,pgdA,unknown | 0.136327518 | -0.02679612 | -0.15177092 | 1 |
| SPV_2349 | unknown | 0.084636898 | 0.041619087 | -0.02754069 | 1 |
| SPV_2351 | msbA | -0.046088084 | 0.126702407 | 0.182173099 | 1 |
| SPV_2362 | unknown | 0.146659898 | 0.015426954 | -0.115778283 | 1 |
| SPV_2363 | unknown | 0.199889748 | 0.018468379 | -0.164619595 | 1 |
| SPV_2364 | unknown | 0.103352227 | 0.170706836 | 0.081147226 | 1 |

| | | | | | |
|---|---|---|---|---|---|
| SPV_2365,SPV_1444 | unknown,thrS | -3.213583831 | -1.925983875 | 1.302509352 | 1 |
| SPV_2366 | unknown | 0.107143881 | 0.019805943 | -0.073355896 | 1 |
| SPV_2367 | unknown | -0.084021765 | 0.051414237 | 0.14568797 | 1 |
| SPV_2370,SPV_2369 | unknown,unknown | 0.076548143 | 0.139151151 | 0.083214123 | 1 |
| SPV_2371 | unknown | 0.060616458 | 0.150958337 | 0.105211714 | 1 |
| SPV_2372 | unknown | -0.029252258 | -0.02607345 | 0.016548987 | 1 |
| SPV_2373 | unknown | 0.105148857 | 0.080116583 | -0.01442445 | 1 |
| SPV_2377 | unknown | 0.076755972 | 0.032182262 | -0.032176532 | 1 |
| SPV_2378 | srf-21 | 0.357447305 | 0.105587142 | -0.235273845 | 1 |
| SPV_2381 | unknown | 0.229518145 | -0.055434365 | -0.271644112 | 1 |
| SPV_2382 | unknown | 0.157254039 | 0.004269768 | -0.134533488 | 1 |
| SPV_2384 | unknown | 0.396226155 | -0.036282468 | -0.414659845 | 1 |
| SPV_2388 | unknown | -0.134209873 | -0.018968723 | 0.128206156 | 1 |
| SPV_2389 | unknown | 0.411944484 | -0.14950707 | -0.530904504 | 1 |
| SPV_2391 | hicA | 0.338310565 | 0.045324224 | -0.276821385 | 1 |
| SPV_2392,SPV_1581 | ssrS,tRNA-Lys-1 | -0.019238588 | 0.164245077 | 0.196631548 | 1 |
| SPV_2395 | unknown | 0.273359692 | -0.001120611 | -0.260682781 | 1 |
| SPV_2398,SPV_2397 | unknown,unknown | 0.063501394 | 0.054645835 | 0.00476298 | 1 |
| SPV_2407 | unknown | 0.188007364 | 0.038316716 | -0.136701008 | 1 |
| SPV_2408 | unknown | -0.058118269 | 0.0702641 | 0.140082404 | 1 |
| SPV_2411 | unknown | 0.209027182 | 0.102702914 | -0.094382601 | 1 |
| SPV_2414 | unknown | 0.071086976 | -0.015469842 | -0.075414749 | 1 |
| SPV_2416 | unknown | 0.039910888 | -0.022221084 | -0.048056741 | 1 |
| SPV_2417 | unknown | 0.240161915 | 0.040851198 | -0.176442765 | 1 |
| SPV_2418 | unknown | -0.010267311 | 0.104363927 | 0.126108571 | 1 |
| SPV_2419 | unknown | 0.129817535 | 0.113859407 | -0.004602659 | 1 |
| SPV_2422 | unknown | -0.016200201 | -0.119031635 | -0.092715629 | 1 |
| SPV_2423 | rpmG2 | -1.608968659 | -1.066101374 | 0.560207026 | 1 |
| SPV_2432 | unknown | 0.005054722 | -0.007429499 | 0.002157333 | 1 |
| SPV_2433,SPV_1902 | srf-24,patA | 0.236312627 | -0.018112237 | -0.239732484 | 1 |
| SPV_2434 | unknown | 0.27610763 | -0.004552769 | -0.252484638 | 1 |
| SPV_2436 | srf-25 | 0.146353622 | -0.010366289 | -0.143968592 | 1 |
| SPV_2437 | unknown | -0.121558304 | 0.013253258 | 0.14386767 | 1 |
| SPV_2441,SPV_1974 | unknown,unknown | 0.108637314 | -0.011706333 | -0.097782362 | 1 |
| SPV_2446 | unknown | 0.304214237 | 0.045217756 | -0.24504357 | 1 |
| SPV_2448 | unknown | -0.05683224 | 0.099467521 | 0.172207414 | 1 |
| SPV_2450 | unknown | -0.093389137 | -0.00071721 | 0.104654122 | 1 |
| SPV_2453 | unknown | -0.023339341 | -0.080153363 | -0.041971692 | 1 |
| SPV_2454,SPV_2028 | srf-29,cbpD | 0.236996843 | 0.001373187 | -0.220927118 | 1 |
| SPV_2455 | unknown | 0.130763735 | 0.101565176 | -0.013698894 | 1 |
